# Human CD8-iTreg are potent GVHD suppressors and tumoricidal effectors by release of Granzyme-K^+^ Supramolecular Attack Particles

**DOI:** 10.64898/2026.06.18.731665

**Authors:** Jemma H. Larson, Ewoud B. Compeer, Phillip R. Dougherty, Kyle Smith, Michael C. Zaiken, Olga Margaritaki, Benjamin Kopp, Sujeong Jin, Maria Harkiolaki, Lina Chen, Salvatore Valvo, Claire Staton, Nagaja Capitani, Chiara Cassioli, Nathaniel Payne, Sara Bolivar Wagers, Sophia Hani, Bailey Houle, Yiyun Peng, Cosima T. Baldari, Leslie S. Kean, Harvey Cantor, Glenn Dranoff, Cameron McDonald-Hyman, Keli H. Hippen, Michael L. Dustin, Bruce R. Blazar

## Abstract

Regulatory CD8^+^ T-cells (CD8^+^ Treg) are a distinct yet understudied T-cell subset capable of simultaneous immunosuppression and cytolysis. Here, we characterized induced human CD8^+^ Treg (CD8-iTreg) generated from peripheral blood CD8^+^CD25⁻ T-cells using anti-CD3e mAb-loaded artificial antigen presenting cells, IL-2, TGFβ, and Rapamycin. These CD8-iTreg differentiated into a stable, highly proliferative bifunctional population with suppressive activity comparable to CD4-iTreg while retaining cytolytic capacity similar to conventional CD8⁺ cytotoxic T lymphocytes (CTL). Multi-parameter spectral flow cytometry and single-cell RNA-seq revealed a distinct immunoregulatory signature: a predominantly Treg-like profile marked by tissue-residency marker CD103 with increased canonical Treg markers (FoxP3, HELIOS, CD25, CD39, CTLA-4, CCR4, and IL-10) and reduced pro-inflammatory cytokines. A unique cytotoxic program was marked by elevated Granzyme-K (GzmK) and Thrombospondin-4 (Tsp-4), a thrombospondin family extracellular matrix glycoprotein upregulated in activated CD8+ T-cells. Cytolysis was primarily mediated by Perforin (Prf) and multiple Granzymes packaged into Tsp-4⁺ supramolecular attack particles (SMAPs), with GzmK contributing to both cytotoxic and suppressive functions. After anti-CD19scFv CAR (CAR19) transduction, CAR19^+^ CD8-iTreg showed superior *in vivo* anti-tumor efficacy compared with CAR19-CTLs, significantly reducing tumor burden and prolonging survival in a CD19^+^ Nalm-6 human leukemia xenograft model while maintaining low pro-inflammatory cytokine production. In a xenogeneic graft-versus-host disease (GVHD) model with residual human leukemia, CAR19⁺ CD8-iTreg inhibited GVHD lethality and controlled tumor growth without increasing systemic inflammation. Together, these findings support CD8-iTreg–based CAR therapies as a strategy to retain potent anti-leukemic activity while limiting inflammatory toxicities of conventional CAR T-cells, properties particularly beneficial in treating auto- and allo-immune diseases.

**One sentence summary:** CD8-iTreg drive parallel tumoricidal and immunoregulatory functions mediated by releasing Tsp-4^+^ SMAPs containing granzyme K.

## INTRODUCTION

Although post-transplant cyclophosphamide has been shown to reduce the incidence of acute graft-versus-host disease (GVHD), chronic GVHD, infection risk and organ toxicities can pose challenges in patients depending on the allogeneic hematopoietic stem cell transplantation (allo-HSCT) setting (1–3). CD4^+^ Regulatory T-cell (CD4 Treg) based cellular therapies can significantly reduce both acute and chronic graft-versus-host-disease (GVHD) incidence and severity without apparent toxicities, increased infection risk or compromise of a graft-versus-leukemia (GVL) benefit (4, 5).

Despite therapeutic advances, acute lymphoblastic B-cell leukemia (B-ALL) remains a significant cause of cancer-related mortality, with approximately 6540 new cases and estimated 1390 deaths estimated in the US in 2023 (6). Moreover, the 5-year survival in adult patients remains ∼50% (7), highlighting the need for improved treatments. Among these, chimeric antigen receptor (CAR) targeting CD19 (CAR19) T-cell therapy stands out as inducing complete remission in the majority of B-ALL patients (8, 9) although relapse and inflammatory toxicities - including cytokine release syndrome (CRS) and neurotoxicity – can limit long-term efficacy and patient survival (10–14).

Recent advancements have seen the adaptation of CAR Treg therapies in preclinical studies as a promising, targeted approach for preventing and treating GVHD (15). For example, murine CAR19 CD4 Treg adoptive transfer in murine allo-HSCT recipients of conventional T-cell (Tcon) fully protected mice from uniformly lethal acute GVHD (16), whereas in the same experiment, CAR19 CD8^+^ cytotoxic T-cells (CTL) or CAR19 CD4^+^ CTL infusion significantly accelerated GVHD lethality; notably, control EGFR-CD4^+^ Treg confers no GVHD protective advantage (16). Desirable outcomes of allo-HSCT for patients with B-cell malignancy are to spare recipients from GVHD side-effects, while eliminating undesirable tumor cells. Studies have shown that CAR19 CD4^+^ Treg *in vitro* immunosuppression can be mediated by their CTL function (17–20). We reported 100% survival in murine allo-HSCT recipients given CAR19 CD4^+^ Treg, Tcon and CD19^+^ leukemia cells (16). However, *in vivo* human CAR19 CD4^+^ Treg infusion often does not eliminate CAR19 targeted tumor cells in xenogeneic (xeno) recipients (21).

Due to their low circulating frequency and phenotypic marker overlap with senescent and virtual memory T-cells, CD8^+^ regulatory T-cells (CD8^+^ Treg) are a less studied subset with for dual potential suppressive and cytotoxic functions, albeit highly debated (20). *In vivo* murine studies show that murine CD8-iTreg adoptive transfer alone is sufficient to suppress GVHD lethality while maintaining superior GVL activity compared to CD4^+^ iTreg (22). Because *in vitro* generated CD8-iTreg can secrete immunosuppressive cytokines (23–29) and cytotoxic granzymes and perforin (20, 30, 31), we hypothesized that CD8-iTreg could enable adoptive cell therapies that combine tumor cytotoxicity with reduced inflammatory toxicity.

Canonically, cytotoxic CD8^+^ T-cells and NK-cells will polarize and fuse LAMP-1^+^ lytic granules to release perforin-1 (Prf), granulysin (GNLY), and granzymes at the immunological synapse upon T-cell engagement (32). Soluble pore-forming Prf enables soluble granzymes to enter the cytoplasm of the T-cell and trigger apoptosis in a caspase 3-dependent manner (33). Recently, we described a thrombospondin-1 (Tsp-1) dependent pathway orchestrated by supramolecular attack particles (SMAPs), where Prf and/or Granzyme-B (GzmB) are incorporated in non-membranous extracellular particles with a Tsp-1 and Tsp-4 shell (34, 35). Whether cytotoxic Tregs could adopt this pathway, assemble and release SMAPs with similar composition and content, and play a role in an *in vivo* tumoricidal response were unknown.

Here, we report that CD8-iTreg generated from human peripheral blood (PB) CD8⁺CD25⁻ T-cells exhibit both potent *in vitro* and *in vivo* suppressor functions as well as improved cytolytic function and tumor control. We show that CD8-iTreg kill targets through the release of SMAPs that contain multiple granzymes, including Granzyme K (GzmK), and are enriched in Tsp-4, distinguishing CD8-iTreg SMAPs from those produced by conventional CTL, and conferring enhanced tumoricidal capacity. Notably, the dual functionality of CD8-iTreg is primarily mediated by GzmK, indicating a noncanonical mechanism that diverges from established regulatory and effector T-cell programs. This observed dual functionality of CD8-iTreg was further reflected in our *in vivo* studies demonstrating CAR19⁺ CD8-iTreg adoptive transfer successfully controlled CD19^+^ leukemia cells in recipients without exacerbating GVHD lethality or inducing a release of pro-inflammatory cytokines commonly associated with CAR T-cell therapy.

Together, these findings establish GzmK⁺Tsp-4^hi^ CD8-iTreg as a cytotoxic–regulatory T-cell state with therapeutic potential and form a strong rationale for testing GzmK^+^Tsp-4^hi^ CD8-iTreg in future CAR T-cell trials as a next-generation CAR T-cell platform with improved efficacy and safety.

## RESULTS

### TGFβ and Rapamycin induction drives human CD8-iTreg differentiation with dual immunosuppressive and cytolytic potential

A highly proliferative, bifunctional population of human PB–derived CD8-iTreg was generated by adapting our CD4-iTreg induction protocol (36). CD8⁺CD25⁻ T-cells were isolated and stimulated with anti-CD3–loaded KT64/86 artificial antigen-presenting cells, proven to promote expansion of CD4 Treg (37). CD8^+^CD25^-^ T-cells cultured in IL-2 (300U/ml), Rapamycin (109nM) and TGFβ (9ng/ml) generated CD8-iTreg, whereas culture in IL-2 alone (300 U/ml) gave rise to conventional CTLs (**Fig. 1A**). Cultures were restimulated with anti-CD3 mAb-loaded KT64/86 cells every 7-days and maintained at 2.5 × 10⁵ to 5 × 10⁵ cells/ml. After 14-days, CTL expanded 1296 ±342–fold and CD8-iTreg expanded 270 ±87–fold. CD8-iTreg iteratively expanded by 6- to 10-fold per stimulation cycle for up to 5 weeks while retaining their proliferative capacity (**Fig. 1B**).

**Figure 1.**
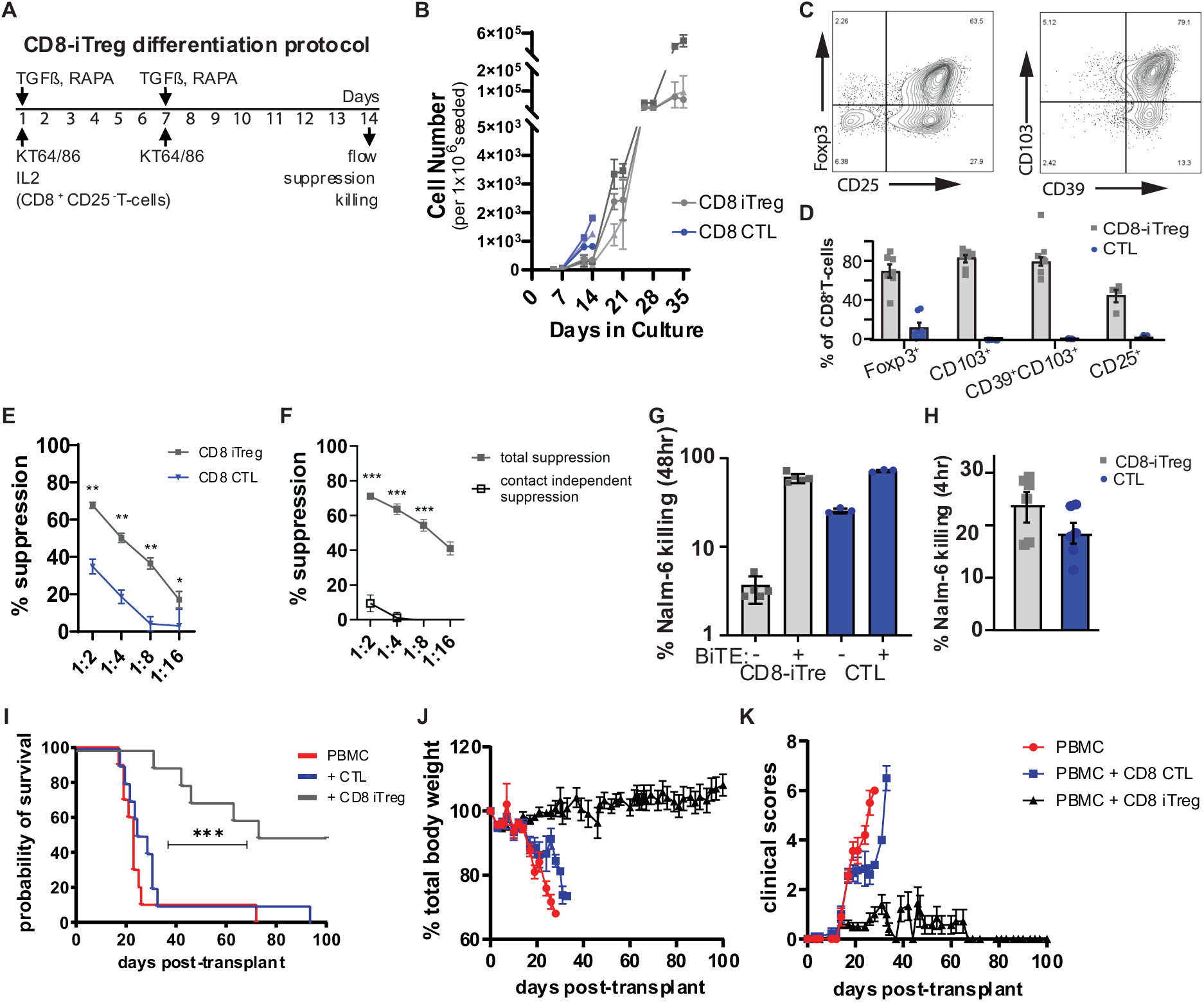
Generation and characterization of bifunctional human PBMC-derived CD103^+^CD8-iTreg. **(A)** Schematic protocol to differentiate CD8-iTreg from CD8^+^CD25^-^ PB T-cells. (**B**) Cumulative cell numbers during differentiation with IL-2(300U/ml), Rapamycin(109nM) and TGFβ(9ng/ml) for CD8-iTreg generation; n=3 representative donors. **(C-D)** Flow cytometry analysis showing FoxP3, CD25, and tissue-residency markers CD103 and CD39 in representative contour plots (**C**) and proportion of CD8^+^ T-cell population for 6 different donors across 2 independent culture experiments (**D**). **(E)** In vitro suppression of activated PBMC proliferation by CTL (cultured with IL-2 alone) or CD8-iTreg at different effector-to-PBMC ratios (1:2, 1:4, 1:8, 1:16) vs PBMC-only controls (0:1), with and without a transwell membrane **(F)**. **(G)** Nalm-6 Percentage killing of Nalm-6 leukemia cells determined by staining with live/dead dye upon 48hr co-culture with CD8-iTreg (Grey) or donor-matched CTL (Blue) at a 5:1 effector-to- target (E:T) ratio with/without αCD3xαCD19 bispecific T-cell engagers (BiTE; 25ng/ml). (**H**) Nalm-6 percentage killing as determined by live/dead dye after 4hrs of co-culturing in 2:1 E:T ratio and with 1ng/ml BiTE. (**I-K**) Xenogeneic GVHD: mice received only human PBMCs (red), PBMCs and CTL (Blue), or PBMCs and CD8-iTreg (Grey) their probability of survival (**I**), weight loss (**J**), and clinical score (**K**). n=10/group; pooled data from 2 independent experiments.

Phenotypically, CD8-iTreg were similar to conventional Treg, gained expression of Foxp3, a dominant transcription factor associated with CD4^+^ and CD8^+^ Treg populations (38, 39), upregulated the high-affinity IL-2 receptor (CD25), and co-expressed tissue resident markers CD103 and CD39 (**Fig. 1C**). Conversely, CD8^+^CD25^-^ T-cells cultured with IL-2 alone (CTL) showed minimal Foxp3 and CD103 expression (4.35 ±1.09% and 0.03 ±0.29%, respectively). CD8-iTreg exhibited increased frequencies of FOXP3⁺ cells (70.0 ±8.8%; P < 0.0110) and CD103⁺CD39⁺ cells (69.5 ±6.6%; *p* <0.0055) relative to donor-matched CTLs (**Fig. 1D**). To assess the suppressive potential of CD8-iTreg, we used an *in vitro* HLA-mismatched CFSE-based proliferation assay (40). CD8-iTreg significantly suppressed human PBMC division at different effector-to-PBMC ratios and were more potent suppressors than donor-matched CTLs (**Fig. 1E**). At a 1:4 ratio, suppression reached 50.2 ±1.7% for CD8-iTreg, comparable to CD4-iTreg (**Fig. S1A**), versus 18.6 ±2.9% suppression in CTL co-culture, establishing the baseline for non-specific inhibition resulting from spatial and metabolic constraints. This suppressive capacity was maintained after prolonged culture, with day-14 and day-35 CD8-iTreg achieving 62% and 59.8% suppression, respectively (**Fig. S1B**). Suppression was primarily contact-dependent since suppression in a transwell system was significantly reduced (**Fig. 1F**).

To compare CD8-iTreg cytotoxic potential to their donor-matched CTLs, we employed a quantitative flow cytometry–based tumor killing assay using αCD3×αCD19 bispecific T-cell engagers (BiTE) to redirect T-cells against CD19⁺ Nalm-6 leukemia cells. With patient-derived Nalm6 tumor cells cultured at a 5:1 effector-to-target ratio for 48hrs, 96.3% and 74.6% of Nalm-6 cells remained viable in the absence of BiTE for CD8-iTreg and CTLs, respectively. Addition of BiTE (25ng/ml) reduced viability to 38.4% and 28.3% for CD8-iTreg and CTLs, respectively (**Fig. 1G**). We hypothesized that the apparent similarity in cytotoxicity at 48hrs could be masked by the high BiTE concentration and the rapid kinetics of BiTE-mediated killing (41). Indeed, at a lower BiTE dose (1ng/ml), within 4hrs CD8-iTreg and CD8 CTLs eliminated ∼24% and ∼19% of Nalm-6 cells, respectively (**Fig. 1H**; n=6/group; p<0.05), suggesting faster early cytotoxic responses by CD8-iTreg.

Considering their notable and sustained cytolytic potential and significant *in vitro* suppressive capabilities, we next assessed whether human CD8-iTreg could suppress T-cell responses *in vivo* using a xenoGVHD model (36, 40). Recipients of human PBMCs alone (2.5×10^6^) uniformly succumbed to GVHD (median survival time (MST) of 23 days), whereas mice treated with CD8-iTreg (4×10^6^) showed a significantly prolonged survival (**Fig. 1I**; MST and long-term survival of 86.5 days; n=10/group; p<0.0001) and reduced weight loss and clinical scores compared to PBMCs and PBMC+CTL recipients (**Fig. 1J and K**). Altogether, these results demonstrate that bifunctional human CD8-iTreg can be robustly expanded *in vitro* and exhibit potent suppressive and cytotoxic activity.

### Human CD8-iTreg are phenotypically distinct from both CD4^+^ and CD8^+^ Treg populations

To define the bifunctional program of human CD8-iTreg, we performed scRNA-seq on CD8⁺CD25⁻ T-cells, from three donors, were cultured 14 days under CTL(+IL-2) or CD8-iTreg(+IL-2, TGFβ, rapamycin) conditions (**Fig. 2A**) from three donors individually studied; data were pooled for analysis. While culture conditions cleanly separated in a dimensionally reduced space, CD103+ iTreg segregated from CD103- counterparts (**Fig. 2B**). RNA Velocity analysis suggested this CD103+ iTreg population to be terminally differentiated as it arose late in latent time analysis (**Fig. S2A and S2B**). This was consistent with the CD103 protein expression observed in CD8iTreg cultures (>80%; **Fig. 1D**).

**Figure 2.**
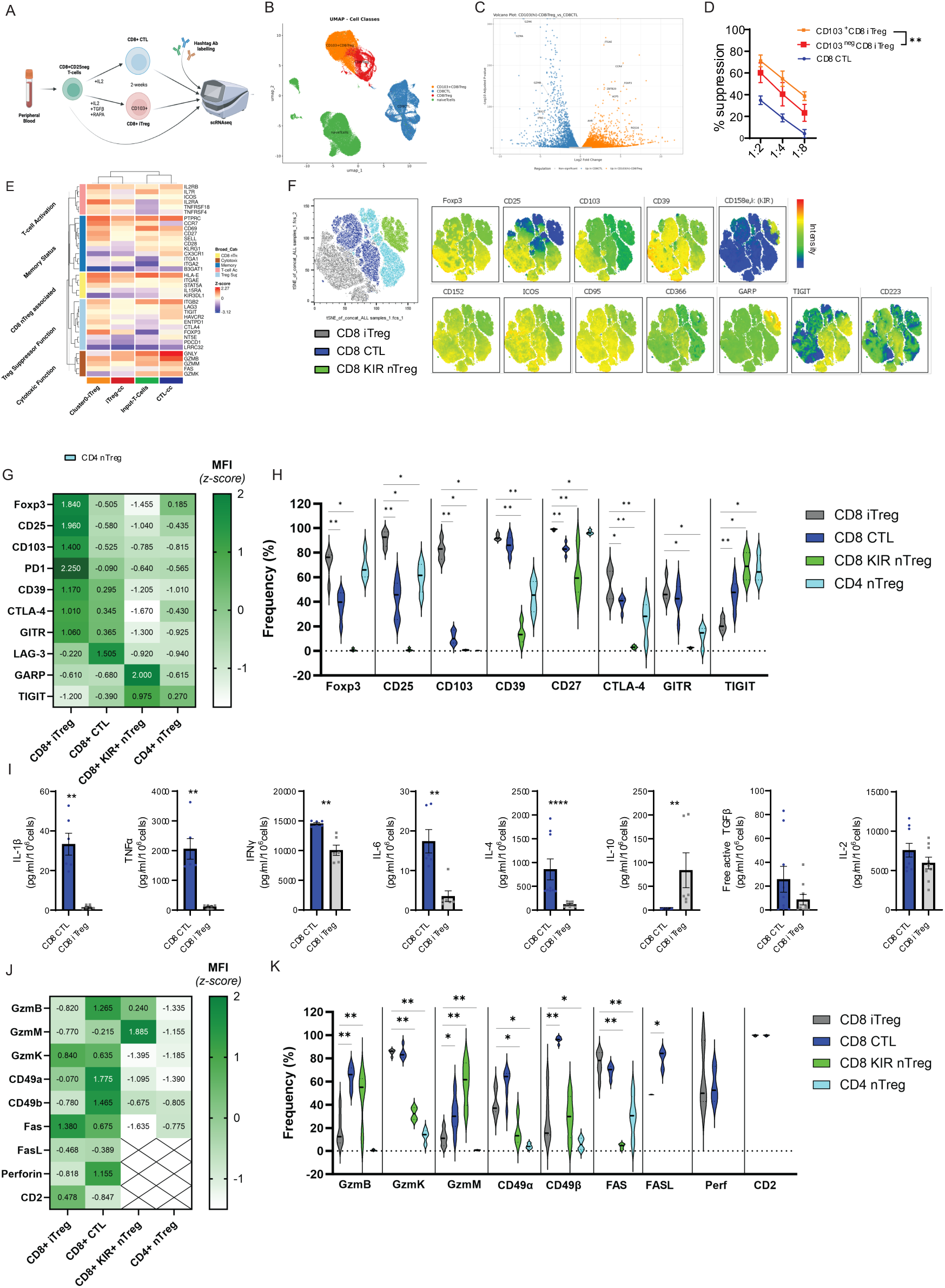
Human CD8-iTreg immunophenotyping identifies cytolytic and suppressor function enrichment in the CD103^+^ subset. CD8^+^CD25^-^ T-cell culture scRNAseq. (**A**) Schema of CD8^+^25^-^ cells cultured from three donors each for two weeks: Input T-cells CD8^+^CD25^-^ T-cells (naïve T-cells), cultured with IL-2 (CD8 CTL), and cultured with IL-2, TGFβ, and Rapamycin (CD8-iTreg) and hashtagged for multiplexed scRNAseq (10x Chromium platform). (**B**) UMAP separation between cultures. CD8iTreg segregated by CD103. (**C**) DEG by functional category. (**D**) Activated PBMC proliferation suppression by CD103^lo^ CD8-iTreg(red), CD103^hi^ CD8-iTreg, CD8 CTL or unsorted CD8-iTreg at effector-to-PBMC ratios (1:2,1:4, 1:8) vs PBMC-only controls (0:1). Multi-parameter spectral flow comparing CD8-iTreg (Grey), CD8 CTL (dark blue), CD8^+^ KIR-expressing Treg (green), and CD4^+^ Treg (light blue). **(E)** DEG volcano plot for CD103^+^ CD8-iTreg and CD8+ CTL cultures. Triplicate samples were pseudo-bulked. DEseq2 was used for DEG analysis (threshold:log2FC >2, p-adj<0.05). Of 1795 DEG, 1,067 were upregulated, 728 downregulated in CD8-iTreg vs CD8 CTL. (**F**) tSNE analysis depicting clusters for each subset showing relative expression of Treg-associated markers; coloring correlates with intensity (Red:high; Blue:low/absent). (**G-H**) Expression (relative MFI z-score) and frequency of Treg-associated markers. (**I**) Cytokines (IL-1β, TNFα, IFNγ, IL-6, IL-4, IL-10, free active-TGFβ, and IL-2) from CD8 CTL (dark blue) vs CD8-iTreg (grey) after PMA/Ionomycin(4h), by LegendPlex^TM^ Panel. (**J-K)** CTL-associated antigens relative expression and frequency.

Pseudo-bulk analysis revealed 1795 DEGs between CD103^+^ CD8-iTreg vs CD8 CTLs (log₂FC >2, adj P <0.05; **Fig. 2C**); 1,067 upregulated and 728 downregulated in CD8-iTreg, respectively. CTLs expressed cytolytic transcripts, including GNLY and Gzm-A, B, K, whereas CD103⁺ CD8-iTreg upregulated *FOXP3, CD25, CTLA4, CD27*, and other Treg-associated genes (**Fig. 2C, S2B**), correlating with enhanced *vitro* suppression (**Fig. 2D**). Notably, CD103⁺ CD8-iTreg also co-expressed cytotoxic genes such as *FAS*, *GZMM*, and *GZMB* alongside Treg-associated transcripts *LAG3*, *CTLA4*, and *TIGIT* (**Fig. 2E**), defining a bifunctional regulatory–cytotoxic program.

Transcriptomic findings were validated by multi-parameter spectral flow cytometry. Donor-matched CD4^+^ natural-Treg (nTreg) characterized by CD4^+^CD25^+^CD45RA^+^CD127^-^ (42), and CD8^+^KIR^+^-Treg (43), were sorted from PBMCs; CD8-iTreg and CTLs were differentiated from CD8⁺CD25⁻ T-cells as above, then cryopreserved. After thawing and overnight resting in RPMI + 5% heat inactivated human-Ab serum (Valley Biomedical) IL-2 (300 U/ml), cells were analyzed across two panels encompassing 32-parameters to evaluate T-cell activation, cytotoxicity, immunosuppression, and canonical Treg CD4^+^ and CD8^+^ nTreg marker expression (**Table S1)**. t-SNE revealed distinct clusters for each cell type, with protein expression overlay distinguishing CD8-iTreg (**Fig. 2F**). CD8-iTreg strongly upregulated Foxp3, CD25, CD103, and CD39, and were enriched in CD4^+^ Treg-associated antigens CTLA-4, ICOS and CD27, while displaying low expression of the canonical CD8^+^KIR^+^-Treg marker KIR3DL2. Quantitative MFI and frequency analysis confirmed increased expression of CD25, CD103, CD39, CD27, CTLA-4, and GITR compared with CD4⁺ Treg, with similar Foxp3 frequencies (**Fig. 2G-H**), consistent with a robust immunosuppressive Treg-like phenotype.

Using a LegendPlex human protein assay, as compared to CD8-CTL, CD8-iTreg produced significantly lower pro-inflammatory cytokine levels of IL-1β (33.33 ±5.54 vs 1.433 ±0.42pg/ml/1M, p=0.002), TNFα (2058 ±347.3 vs 114.3 ±22.84pg/ml/1M, p=0.002), IFNγ (14547 ±184.8 vs 10060 ±861.8pg/ml/1M, p=0.002), IL-6 (17.39 ±2.96 vs 3.503 ±1.38pg/ml/1M, p=0.002), and IL-4 (856.4 ±219.7 vs 119.4 ±16.23pg/ml/1M, p= <0.001), and significantly more IL-10 (2.27 ±0.26 vs 83.72 ±36.83pg/ml/1M, p=0.002) along with lower free active-TGFβ (25.72 ±10.88 vs 8.71 ±4.44pg/ml/1M, p=0.182), and IL-2 (7585 ±870.3 vs 5950 ±738.5pg/ml/1M, p=0.248), respectively, following a 4hr PMA/Ionomycin stimulation (**Fig. 2I**). Similar results were obtained with the RayPlex Human Inflammation Bead Array (**Fig. S2C**).

True to their bifunctional nature, at the protein level, CD8-iTreg co-expressed moderate levels of cytolytic-associated antigens GzmB, GzmM, CD49a, CD49b, FasL, Prf, and notably higher CD95 (Fas), CD2 and GzmK levels compared to donor-matched CTLs (**Fig. 2J**). CD95^+^ cells were present at higher frequency, GzmK^+^ frequencies were similar to CD8-CTLs and frequency of GzmB^+^, GzmM^+^ and Prf^+^ cells were lower. Collectively, these data reflected cytotoxicity and immunoregulatory functions. CD8-iTreg were distinct from canonical CD8⁺KIR⁺ Treg (20, 43, 44), exhibiting low expression and frequency of KIR^+^ Treg-associated antigens HLA-E, CD57, KLRG1, CXCR1, and KIR3DL1/DL2 (**Fig. S2D**) and a unique signature more closely aligned with canonical CD4^+^ Treg and CD8^+^ CTLs; these data exclude PB-derived CD8^+^KIR^+^ Treg contamination.

### CD8-iTreg drive parallel cytotoxic and immunoregulatory responses via Gzm-K

Having defined the unique phenotypic signature of CD8-iTreg, we investigated the mechanisms underlying their dual cytotoxic and suppressive functions. Previous studies of CD8⁺ Treg implicated GzmA, GzmB, and Prf–dependent killing(20). Therefore, we evaluated their specific contributions in CD8-iTreg–mediated cytotoxicity using a Prf inhibitor (CMA, 1μM), pan-Gzms A/B/H inhibitor 3,4-dichloroisocoumarin (DCI; 40μM), GzmB-specific inhibitor (Z-AAD-CMK; 5μg/mL), GzmK inhibitor Bosutinib (10μM, 48hr pre-treatment), or CRISPR-Cas9a targeting GNLY or GzmK, compared with vehicle (DMSO) and non-targeting controls. Human leukemia patient-derived CD19^+^ Nalm-6 leukemia cells were used as targets at a 2:1 E:T ratio and killing was assessed by live/dead staining after 4hrs in the presence of aCD3xCD19 BiTE (1ng/ml); long-term killing was monitored in an Incucyte system using 25ng/ml BiTE.

Both human CD8-iTreg and conventional CTLs were largely Prf- and Gzm-dependent, since CMA or DCI significantly reduced killing in both cell types, compared to vehicle or untreated controls (dashed lines). GzmB inhibition markedly reduced CTL-mediated killing but had minimal impact on CD8-iTreg (**Fig. 3A**), consistent with their low GzmB and higher GzmA/M/K protein expression (**Fig. 2J**). GNLY knockout modestly reduced killing in both cell types, reflecting its low baseline expression (**Fig. 2E**). In long-term killing assays with high BiTE concentrations, Prf (**Fig. 3B**) or Gzm-A/B/H (**Fig. 3C**) inhibition moderately reduced overall CD8-iTreg cytotoxicityCD8.

**Figure 3.**
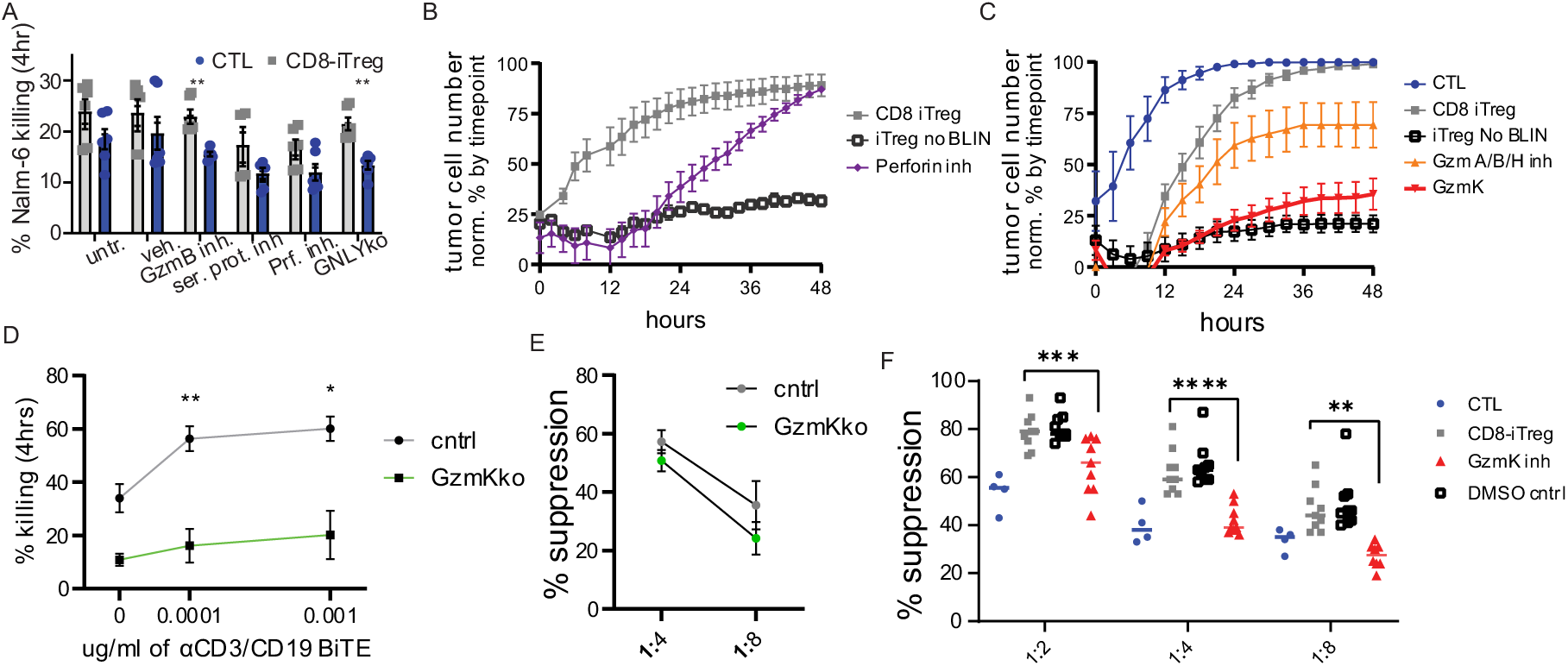
Perforin and Granzyme-K contribute to CD103^+^ CD8-iTreg dual cytotoxic and regulatory functions. (**A**) Percentage of Nalm-6 killing determined by live/dead dye after 4hrs of co-culturing in 2:1 E:T ratio in presence of 1ng/ml BiTE in combination with inhibitors against GzmB, all serine proteases, Prf function, CRISPR-Cas9a targeting tumoricidal granulysin (GNLY) or vehicle controls (PBS; DMSO). (**B-C**) Long-term tumoricidal potential of CD8-iTreg was evaluated by continuous Incucyte killing assay. CD8-iTreg were co-cultured with Nalm-6 cells at 2:1 E:T ratios. **(B)** Prf-dependent cytotoxicity was evaluated by pre-treating CD8-iTreg with 1μM Concanamycin A (CMA) for 2hrs. **(C)** The contribution of Gzm-A, B, and H was assessed by pre-treating CD8-iTreg with 3,4-dichloroisocoumarin (DCI; 40μM) for 3hrs. GzmK inhibition of CD-iTreg by Bosutinib (10 µM; 48h) pretreatment; all inhibitor treatments were compared against equivalent volume DMSO vehicle or untreated controls. **(D) S**hort-term and tumoricidal potential of CD8-iTreg electroporated with GzmK CRISPR/Cas9 RNP vs non-targeting control RNP. **(E-F)** In vitro suppression of activated PBMC proliferation by CD8-iTreg at 1:4 and 1:8 effector-to-PBMC ratio vs PBMC-only controls (0:1), following **(E)** CRISPR/Cas9 RNP GzmK-ko or **(F)** pretreatment with 40μM DCI (4hrs), or 10μM Bosutinib (48hr), vs DMSO vehicle and untreated controls.

Our phenotypic analysis revealed upregulated GzmK in CD8-iTreg relative to CTLs (**Fig. 2J**), suggesting a possible role in their function. As described previously (45), Bosutinib (10μM), an established dual Src and ABL kinase and GzmK inhibitor, treatment significantly reduced GzmK expression (97.7% to 26.6%) and not Prf or GzmB in CD8-iTreg (**Fig. S3A-B**) which resulted in an almost complete loss of tumor killing *in vitro* (**Fig. 3C**). However, given that Bosutinib is a non-specific dual Src/ABL kinase inhibitor, we subsequently confirmed the specific role of GzmK using GzmK-targeting CRISPR/Cas9 ribonucleoproteins (RNPs) in CD8-iTreg, achieving 65% reduction in GzmK expression compared to controls, without affecting Prf, granzymes and GNLY expression (**Fig. S3C**). Loss of GzmK in CD8-iTreg corresponded with a 17% and 20% reduction in Nalm-6 killing after 4hr co-cultures (**Fig. 3D**), supporting that CD8-iTreg employ a combination of Prf and multiple granzymes, with GzmK playing an important role in total cytotoxic potential.

Similarly, CRISPR/Cas9 RNP targeting GzmK, reduced suppression by 11.3% and 38% at 1:4 and 1:8 effector-to-PBMC ratios, respectively (**Fig. 3E**), suggesting a contributing role of GzmK in CD8-iTreg mediated suppression. In parallel, Bosutinib pre-treatment also significantly reduced the suppressive capacity of CD8-iTregs, reducing total suppression from 57.2 ±4.3% in untreated controls to 38.5 ± 2.1% at a 1:4 effector-to-PBMC ratio (**Fig. 3F**). In contrast, inhibition or blockade of GzmA/B/H, Caspase, CD39, IL-10, CTLA-4, or CD25 did not significantly affect CD8-iTreg–mediated suppression (**Fig. S3E**). Altogether, these findings position GzmK as an important mediator of CD8-iTreg dual functionality.

### Human CD8-iTreg generate Thrombospondin 4 (Tsp4)-enriched supramolecular attack particles with superior cytotoxicity

Having previously identified a cytotoxic pathway in CTLs involving SMAPs (34), we investigated whether CD8-iTreg also deploy SMAP-mediated cytotoxic mechanisms. Mass spectrometry analysis of material deposited and captured on activating planar supported lipid bilayers (pSLBs) revealed substantial overlap between proteins released by CD8-iTreg and previously characterized CTL SMAP components (34) (**Table S2).** Canonical SMAP glycoproteins Tsp-1 co-assembly with Tsp-4 is required for effective SMAP formation and CTL cytotoxicity (35). Tsp-1, Tsp-4, and GzmB peptide fragments were significantly enriched on activating pSLBs compared with ICAM-1–only control bilayers (**Fig. 4A**), consistent with CD8-iTreg release of SMAP-associated proteins in a TCR-dependent manner as seen with CD8^+^ CTLs. However, there was a notable difference in peptide fragment abundance in SMAP composition. CTL-derived SMAPs contained relatively higher Tsp-1 levels, whereas CD8-iTreg-derived SMAPs were enriched in Tsp-4. This was corroborated by Tsp-4 CD8-iTreg immunostaining compared with donor-matched CTLs by flow cytometry (**Fig. 4B**) and western blot analysis (**Fig. S4A**). Notably, Tsp-4 expression in CD8-iTreg correlated with increased expression of CD103, Prf, GzmA, GNLY, and reduced expression of GzmB (**Fig. S4B**). These findings suggested that CD8-iTreg may utilize a SMAP program enriched in Tsp-4 that could influence the cytotoxic cargo.

**Figure 4.**
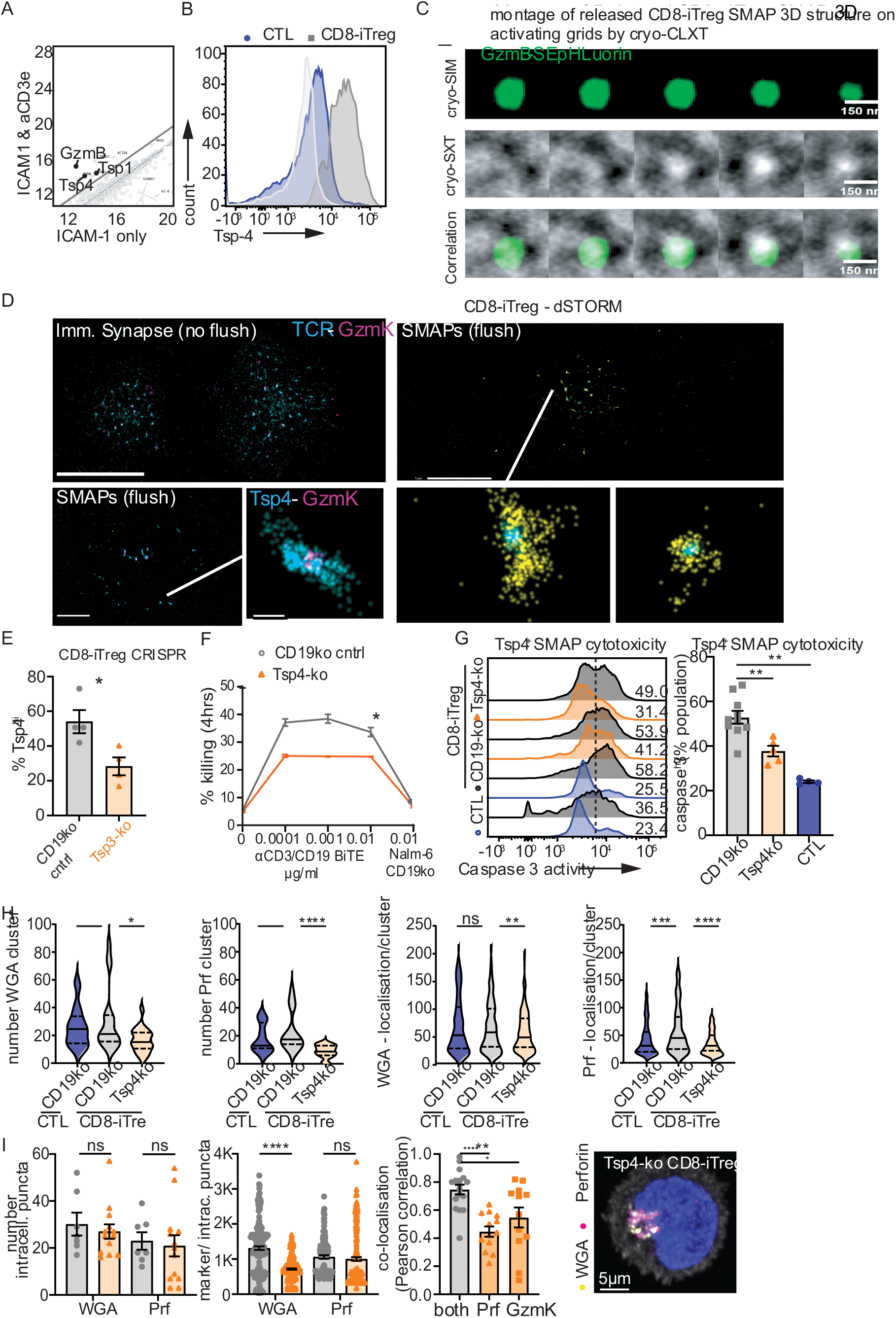
Tsp-4^+^ SMAPs enhance CD8-iTreg-mediated, Prf/GzmK-dependent leukemia cell killing. **(A)** Mass spectrometry comparison of peptide fragments in particles deposited by CD8-iTreg after 90 min on activating (ICAM-1/anti-CD3ε) versus non-stimulating (ICAM-1 only) pSLBs. **(B)** Representative flow cytometry histogram showing total Tsp-4 expression in CD8-iTreg (grey) compared to donor-matched CTLs (blue). **(C)** 3D cryo-correlative SIM and Soft X-ray Tomography reconstruction of a GzmB-mCherry-SEpHluorin-positive CD8-iTreg SMAP released on activating Quantifoil grids. **(D)** dSTORM analysis of immunological synapse (top left) or SMAPs (others) on activating pSLBs stained for Tsp-4, GzmK, Prf, GzmA, and WGA. **(E)** Tsp-4 expression in CD8-iTreg from four donors following CRISPR–Cas9 targeting of Tsp-4 (orange) or CD19 control (grey). **(F)** Cellular Cytotoxicity assay measuring Nalm-6 killing after 4 h co-culture with CD8-iTreg (E 1:2) and αCD3×αCD19 BiTEs following Tsp-4 or CD19 editing. **(G)** SMAP-mediated cytotoxicity measured by caspase-3 activation in Nalm-6 cells exposed to CTL-or CD8-iTreg-derived SMAPs. **(H)** dSTORM quantification of Prf and WGA-positive SMAPs, including DBSCAN particle counts and localizations per cluster. **(I)** 3D Airyscan imaging of CD19- or Tsp-4-knockout CD8-iTreg stained for Prf or GzmK after two-hour incubation with fluorescent WGA to identify SMAP-containing lytic granules.

We previously characterized SMAPs as carbon-dense particles with a heterogeneous size distribution, using cryo-Soft X-ray Tomography (cryo-SXT), averaging 111 ±36 nm as defined by direct Stochastic Optical Reconstruction Microscopy (dSTORM) (*37*). Cryo-SXT is a non-destructive 3D imaging technique that exploits the preferential natural absorption of X-rays at the “water window” of the X-ray spectrum by carbon-rich cellular structures in unstained, vitrified specimens, achieving a resolution up to ∼25 nm. The B24 beamline at Diamond Light Source, offers cryo-SXT, using X-rays from a bending magnet on the synchrotron ring, passed to focusing and dispersive optics before reaching the microscope. This system uses a capillary instead of a zone plate condenser, to deliver X-rays to the sample plane, while they are focused to the imaging plane of a camera with a 25 nm Frensel zone plate (46, 47). Here we determined whether the MS-identified components are released by CD8-iTreg as *bona fide* SMAPs in a correlative setting, combining cryo-SXT with 3D SIM of the same regions (**Fig. S5 and Movie S1**). 3D cryo-CLXT revealed released GzmB-Superecliptic(SE)SEpHluorin-positive structures, consisting of an X-ray-dense partially enclosed “shell”, structurally corresponding to the larger range of SMAPs **(Figs. 4C and S5)**, as previously described by dSTORM (34).

Subsequent dSTORM analysis demonstrated that GzmK, GzmB, GzmA, and Prf-1 co-localized with Tsp-4^+^ SMAPs, derived from both CTLs and CD8-iTreg (**Fig. 4D and S6A**). Wheat germ agglutinin (WGA) staining, which labels SMAP glycoproteins independently of thrombospondin subtype, detected 24.4±3.3 WGA⁺ SMAPs per CD8-iTreg, and Tsp4-positive SMAP counts were similar between CD8-iTregs and CTLs (31±3 and 26.5±2 per cell, respectively) (**Fig. S6B**). Together, these data indicate that both cell types release comparable SMAP numbers. However, paired experiments imaged under identical conditions revealed significantly greater Tsp-4 localizations within CD8-iTreg-derived compared with CTL-derived SMAPs (45±13 vs 38±10; p=0.001). Consistent with this observation, CD8-iTreg secreted substantially more cytotoxic SMAPs, including GzmB⁺ SMAPs (27.7 ±0.9 vs 4.3 ±0.7; p=0.0008) and Prf⁺ SMAPs (32.4 ±8.4 vs 12.1 ±1.8; p=0.0196). Higher Prf density was displayed within CD8-iTreg than donor-matched CTL SMAP structures (71.5 ±19.5 vs 35.7 ±1.1; p=0.0102) (**Fig. S6C-D**). Together, these results indicate that although CD8-iTreg and CTLs release similar SMAP numbers, CD8-iTreg-derived SMAPs are enriched in cytotoxic proteins, consistent with enhanced tumoricidal potential (p=0. 0297; **Fig. 3A**).

Because Tsp-4 represents a major structural CD8-iTreg-derived SMAP component, we next investigated their functional contribution to CD8-iTreg cytotoxicity. Electroporation of Tsp-4 targeting CRISPR/Casp9a RNPs reduced Tsp-4 expression in approximately half of the CD8-iTreg population, whereas an irrelevant control guide RNA targeting CD19 had no effect (**Fig. 4E and S7A**). No significant changes in total intracellular Prf, GzmB, GzmA, GzmK and GNLY expression levels were observed (**Fig. S7B**). Using a short-term αCD3–αCD19 BiTE assisted killing assay (1ng/ml), we observed that reduced Tsp-4 expression corresponded to a proportional decrease in Nalm-6 leukemia cell death after 4hrs (**Fig. 4F**), without killing CD19-negative Nalm- 6 cells (**Fig. S7C**), demonstrating that Tsp-4 expression contributes directly to antigen-specific CD8-iTreg-mediated cytotoxicity. Notably, targeting Tsp-1 alone, or co-targeting Tsp-1 and Tsp-4, produced a similar reduction in cytotoxicity as Tsp-4 depletion alone at this early time point, indicating that both Tsp-1 and Tsp-4 are required for efficient short-term killing. This dependency was observed in both CD8-iTreg and conventional CTLs, suggesting that the underlying SMAP-associated molecular machinery is conserved across distinct CD8⁺ T-cell states (**Fig. S7D**).

To specifically examine SMAP-mediated killing, donor-matched CTLs and CD8-iTreg targeted with Tsp-4 or control CD19 guide RNA were activated on pSLBs containing ICAM-1 and anti-CD3ε for 90 minutes; cells then were removed by PBS flushing to leave deposited SMAP structures on the bilayers (34). dSTORM analysis confirmed the presence of Prf-containing SMAPs on these surfaces. Control CD8-iTreg deposited an average of 31.2 ±5.6 WGA⁺ SMAPs and 23 ±4 Prf⁺ SMAPs, whereas Tsp-4 targeted CD8-iTreg deposited significantly fewer WGA⁺ puncta (16.6 ±1.7) and Prf-containing SMAPs (9 ±1) (**Fig. 4G**), indicating that Tsp-4 expression is required for efficient cytotoxic SMAP release. When Nalm-6 cells were incubated on these surfaces, SMAP structures left behind by CD8-iTreg, Tsp-4 deficient CD8-iTreg and donor-matched CTLs induced 53±2.7% (n=10), 37.2±2.2% (n=5; p=0.0033), and 24.6±0.4% (n=5; p=0.0007) Nalm-6 cell death, respectively. These findings indicate that loss of Tsp-4 expression in ∼50% of CD8-iTreg population proportionally and significantly reduced SMAP-mediated cytotoxicity.

Previous studies using stimulated emission depletion (STED) microscopy and correlative light-electron microscopy demonstrated that exogenously added WGA is incorporated into a lytic granule subset containing Tsp-1 and/or Tsp-4 that are subsequently released as SMAPs (35, 48), providing a marker for SMAP-forming compartments. To determine whether Tsp-4 affects SMAP formation or cargo composition, CD8-iTreg were incubated with exogenous fluorescent WGA for 2hrs at 37°C prior to fixation and staining for Prf and Tsp-4. Tsp-4 staining. Three-dimensional Airyscan imaging revealed no significant differences in the intracellular WGA⁺ Prf⁺ puncta number between control and Tsp-4-deficient CD8-iTreg (30 ±5 vs 27 ±3, respectively) (**Fig. 4H**), indicating that SMAP-like compartments can form in the absence of Tsp-4. However, SMAP structures in Tsp-4-deficienT-cells incorporated significantly less WGA, as indicated by reduced mean WGA fluorescence intensity per punctum (1313 ±52.3 vs 714 ±19.5; p<0.0001), suggesting SMAP content changes. Although Prf-positive puncta number and Prf intensity per punctum did not change dramatically (**Fig. 4I**), Prf localization to WGA-positive SMAP compartments was significantly reduced, reflected by a decrease in the Pearson correlation coefficient from 0.7 ±0.03 in control cells to 0.45 ±0.04 in Tsp-4 KO CD8-iTreg. Similarly, GzmK localization to WGA-positive SMAP compartments was markedly reduced from Pearson coefficient 0.75 ±0.03 in control cells to 0.55 ±0.07 in Tsp-4 KO CD8-iTreg (**Fig. 4I**). As thrombospondins are multimeric scaffold proteins capable of organizing supramolecular protein assemblies, we propose that elevated Tsp-4 expression in CD8-iTreg promotes the formation of a structural scaffold that stabilizes or concentrates cytotoxic proteins such as Prf and GzmK within SMAPs prior to TCR signaling-induced deposition, thereby enhancing CD8-iTreg cytolytic capabilities. Together, these findings identify Tsp-4 as a key regulator of CD8-iTreg SMAP composition that enhances the cytotoxic potency of these nanoparticles, contributing to the overall cytotoxic potential of CD8-iTreg.

### Anti-hCD19-CAR engineering preserves CD8-iTreg bifunctional immunosuppression and antigen-specific cytotoxicity

Having established the unique bifunctional properties of CD8-iTreg, we hypothesized that these cells could represent a novel strategy for next-generation CAR T therapies, potentially addressing both the potency and safety limitations of current approaches (49). To explore the therapeutic potential of CD8-iTreg, we engineered CD8-iTreg to express CAR19. Activated PB CD8⁺CD25⁻ T-cells were transduced with a second-generation CAR19 lentivirus comprising an EGFR marker domain and 4-1BB(CD137)–CD3ζ signaling modules (**Fig. 5A**). EGFR⁺ cells were enriched and expanded to generate CAR19⁺CD8^+^ T-cells. Mock-transduced or CAR19-EGFR⁺ CD8⁺ T-cells were subsequently differentiated into CD8-iTreg or conventional CTLs using the culture conditions described above.

**Figure 5.**
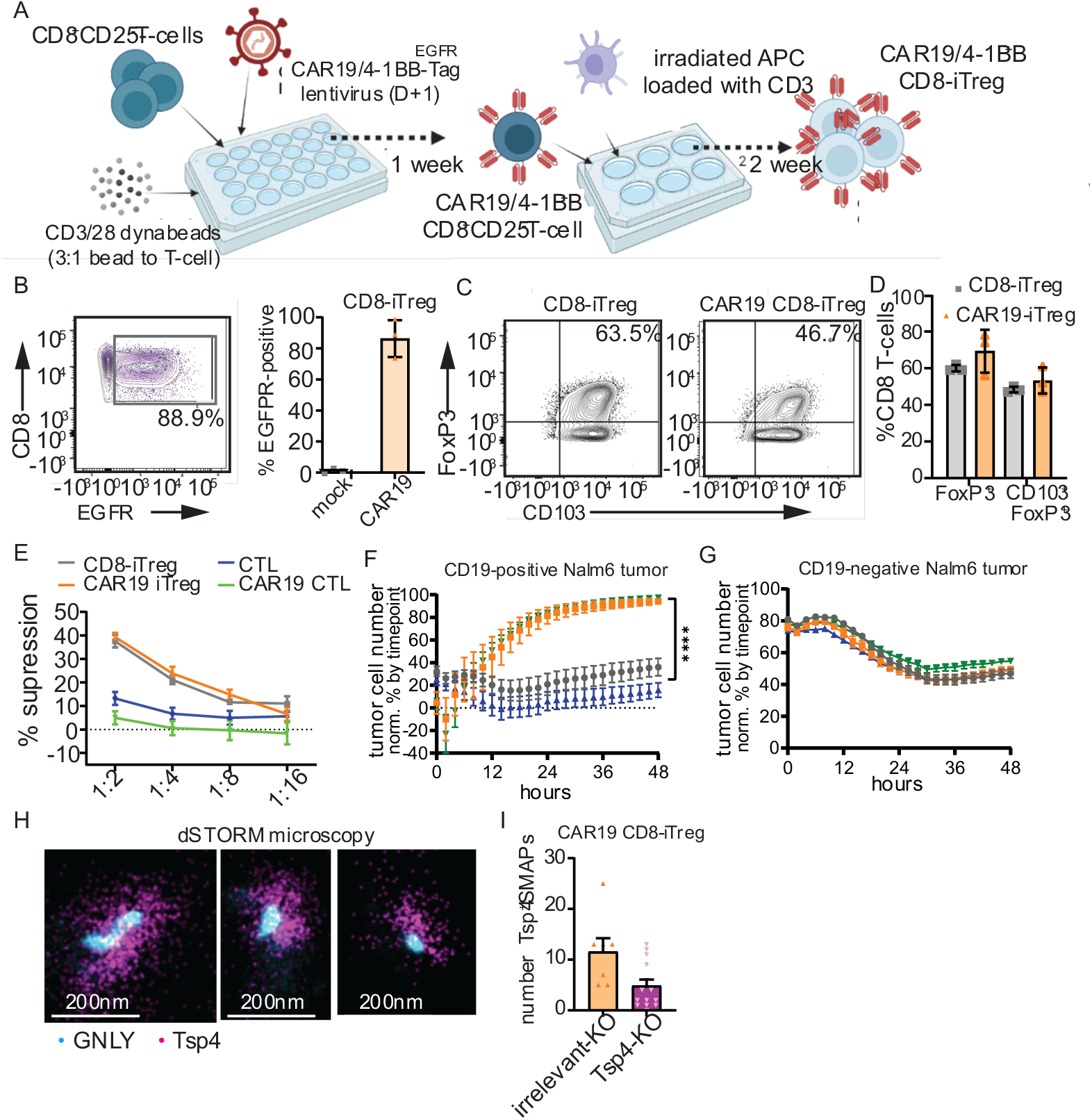
CAR19^+^ CD8-iTreg retain stable dual-functional signature. **(A)** Illustration of protocol to generate CAR19+ CD8-iTreg. **(B)** Representative contour plot and frequency of CAR19-positive CD8-iTreg through detection of EGFPR tag in CAR19 lentivirus. (**C**) Representative CD103 and FoxP3 contour plots and in mock vs CAR19 transduced CD8-iTreg **(D)**. **(E)** In vitro suppression of activated PBMC proliferation by mock vs CAR19 CD8-iTreg vs CD8 CTLs at various effector-to-PBMC ratios. **(F-G)** 48hr continuous Incucyte killing assay of CD8-Treg (Grey), CAR19^+^ CD8-iTreg (orange), donor-matched CTL (Blue) and CAR19 + CTL (Green) co-cultured with wither CD19^+^ **(F)** or CD19-ko **(G)** Nalm-6 tumor cells at 5:1 ratio effector to tumor ratio. (**H**) dSTORM analysis of particles left behind after 90 minutes incubation on activating pSLBs with CD19 and ICAM-1 post flushing with PBS, stained with antibodies against Tsp-4 and GNLY, shown as representatives. (**I**) Quantification of Tsp-4 positive SMAPs released by CD8-iTreg from 3 different donors upon electroporation of RNP complexes with gRNA against Tsp-4 (purple) or irrelevant marker CD19 (orange) as control.

After the two-week differentiation period, CAR19⁺ CD8-iTreg retained stable CAR expression, with >80% of cells remaining CAR19-EGFR⁺ (**Fig. 5B**). CAR expression did not alter the CD8-iTreg phenotypic characteristics, as these cells maintained comparable FoxP3 and CD103 levels and frequencies (**Fig. 5C, D**). Likewise, CAR19 expression did not affect the *in vitro* suppressive capacity of CD8-iTreg or the functional activity of CTLs (**Fig. 5E**).

We next evaluated the tumoricidal activity of CAR19-engineered cells. Both CAR19⁺ CD8-iTreg and CTLs exhibited comparable cytotoxic activity against CD19⁺ Nalm-6 leukemia cells in a continuous Incucyte™ killing assay at a 5:1 effector-to-target ratio (**Fig. 5F**). No cytotoxicity was observed against CD19-knockout (CD19-KO) Nalm-6 cells, confirming CAR-mediated killing antigen-specificity (**Fig. 5G**).

We next examined whether CAR19 expression altered the cytotoxic secretion machinery of CD8-iTreg. Engagement of CAR19⁺ CD8-iTreg with CD19 and ICAM-1 proteins on pSLBs enabled formation of a functional immunological synapse (**Fig. S8**). dSTORM imaging of Tsp-4^hi^ SMAPs with confirmed presence of cytotoxic cargo, including depicted GNLY, deposited by CAR19⁺ CD8-iTreg on CD19⁺ICAM-1⁺ pSLBs (**Fig. 5H**; shown as 3 representative images). To determine the contribution of Tsp-4 to SMAP release in CAR19⁺ CD8-iTreg, CAR19^+^ CD8-iTreg cells were targeted with Tsp-4 gRNA. Disruption of Tsp-4 expression significantly reduced SMAP deposition following ICAM-1 and CD19 engagement, decreasing the number of Tsp-4^hi^ SMAPs released from 11 ±3 per cell by control CAR19⁺ CD8-iTreg to 4 ±1 per cell by Tsp-4 KO CAR19⁺ CD8-iTreg (**Fig. 5I**, p = 0.002). Together, this data demonstrates that engineering CD8-iTreg to express CAR19 does not affect their potent cytotoxic activities through Tsp-4^+^ SMAPs.

### CAR19^+^ CD8-iTreg exhibit superior tumoricidal activity in vivo

We next evaluated CAR19^+^ CD8 iTreg vs CTL *in vivo* efficacy in a xenoGVL model. NSG recipients (n= 5/group) were given CD19^+^ Nalm-6-GFP/firefly-luciferase (luc) tumor cells (1x10^6^; day-0), followed by adoptive transfer of non-transduced (mock), CAR19-4/1BBexpressing CD8-iTreg or CTLs (1x10^7^), or no T-cells on day-7 (**Fig. 6A)**. In mice given a lower number of tumor cells (luc-GFP^+^ Nalm-6, 5x10^5^), no survival or tumor burden differences were seen between CAR19^+^ CD8-iTreg and CTLs conferred comparable survival and tumor burden (**Fig. S9A**). However, at a higher tumor cell number infused (1x10^6^), mice given CAR19^+^ CD8-iTreg had a significantly improved survival as compared to CAR19^+^ CTL (MST 81 vs 53-days; p<0.0018; **Fig. 6B**). In contrast, mice receiving Nalm-6 tumor alone, or mock-transduced iTreg or CTLs, showed rapid disease progression (MST 16-18 days). Both CAR19^+^ iTreg and CTLs exhibited no anti-leukemic effect against CD19-KO Nalm-6 targets *in vivo*, confirming antigen specificity (**Fig. S9B**). CAR19^+^ CD8-iTreg recipients also had a significant improvement in early weight loss (**Fig. 6C**) compared to CAR19^+^ CTL recipients. This correlated with a rapid and significant Nalm-6 tumor burden reduction *in vivo* (**Figs. 6D-E**), 7-days post T-cell adoptive transfer (14-days post-tumor infusion). CAR19^+^ CD8-iTreg recipients maintained undetectable levels of luc^+^ Nalm-6 tumor *in vivo* for 42-days post T-cell infusion, at which point we identified a resurgence of luc^+^ Nalm-6 tumor cells in CAR19^+^ CD8-iTreg recipients (**Fig. 6E**). Altogether, these data demonstrate that CAR19^+^ CD8-iTreg have a superior capacity to reduce tumor load and tumor-related mortality *in vivo* compared to conventional CAR T-cells that could be advantageous to CAR-CTLs immunotherapy.

**Figure 6.**
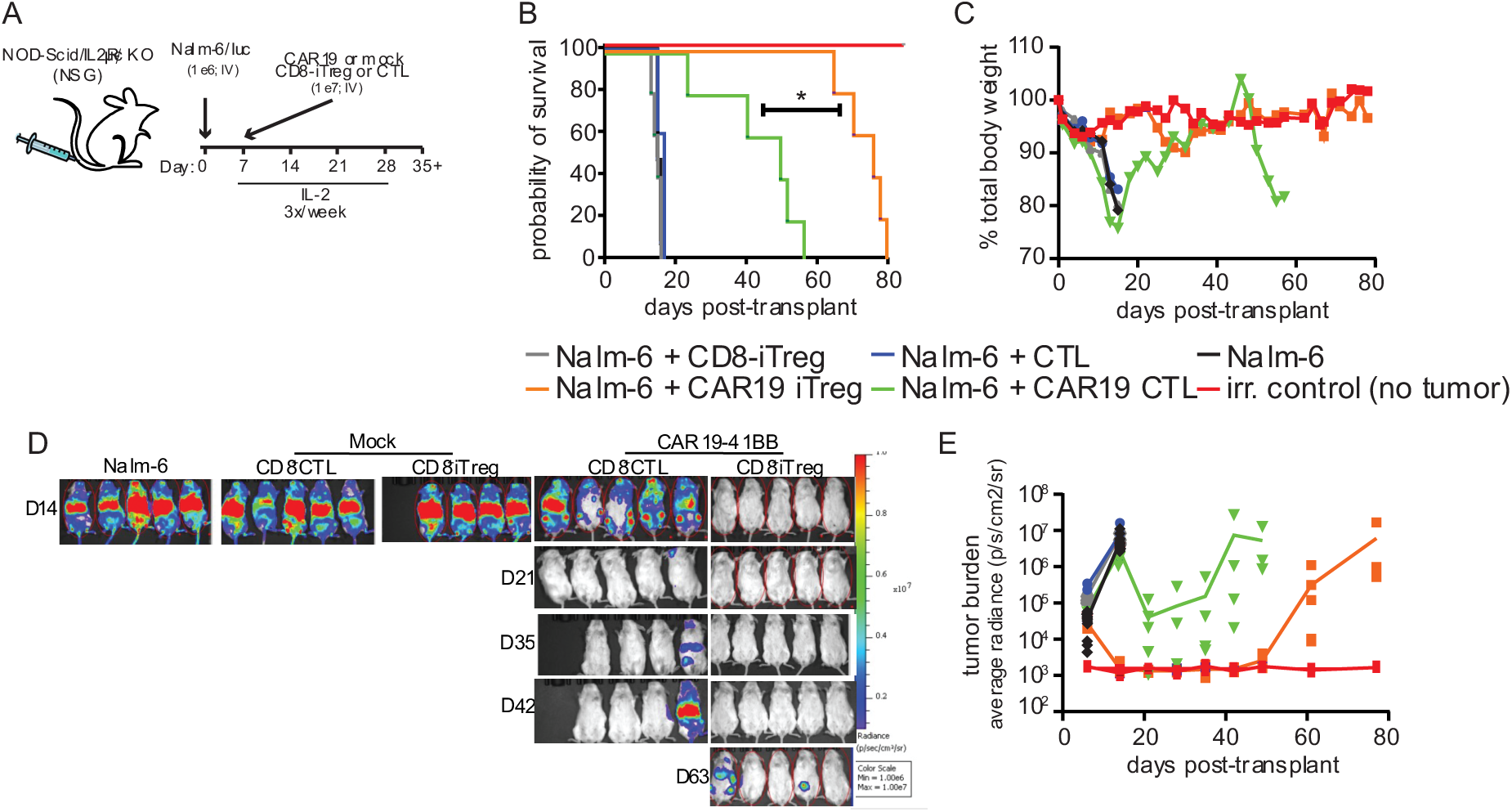
CAR19-4/1BB CD8-iTreg have superior anti-tumor activity against CD19^+^ Nalm-6 tumor in vivo compared to CAR19 CTLs. (**A**) Schematic depiction of the luciferase-positive Nalm-6 tumor model with supplemental IL-2 (250kIU/mouse; 3x/week for 21-days). Probability of survival by log-rank analysis (**B**), body weights (**C**), with representative in vitro imagine system (IVIS) images **(D)** and average tumor burden as determined by BLI, shown as average radiance, (**E**) from mice receiving no tumor cells(Black) or tumor cells alone(Red), or tumor cells with CD8-iTreg(Grey), CAR19^+^ CD8-iTreg(orange), CTL(blue) or CAR19^+^ CTL(Green). n=5/group; representative of 3 independent experiments.

Since clinical administration of IL-2 can be associated with severe side-effects and toxicities, including systemic inflammation and exacerbation of GVHD (49), in parallel, we evaluated the necessity of exogenous IL-2 administration of CAR19^+^ CD8-iTreg success *in vivo*. Importantly, in absence of exogenous IL-2, CAR19⁺ CD8-iTreg maintained substantial, albeit reduced, anti-leukemic activity (**Fig. S9C**), demonstrating their ability to persist in the absence of supplemental IL-2. Therefore, all subsequent *in vivo* experiments were executed without administration of endogenous IL-2.

We next evaluated *in vivo* CAR19^+^ CTL frequency and persistence relative to the infused number of Nalm-6 tumor cells. Irradiated NSG mice were engrafted with 1x10^6^ Nalm-6 tumor cells, followed by adoptive transfer on day-7 of CAR19-expressing luc-GFP⁺ CTLs or CD8-iTreg (1×10⁷ each) (**Fig. 7A**). Longitudinal total body BLI imaging (IVIS) revealed that luc-GFP^+^ CAR19^+^ CD8-iTreg exhibited significant post-tumor survival compared to luc-GFP^+^ CAR19^+^ CTLs (MST 40-days vs 20-days, respectively, **Fig. 7B**; p <0.0018). Monitoring the average radiance (p/s/cm2) over time revealed luc-GFP^+^ CAR19^+^ CTL to reach significantly higher peak expansion *in vivo* compared to CAR19^+^ CD8-iTreg 10-14 days post T-cell infusion (**Fig. 7C**). This expansion corresponded with a rapid reduction in Nalm-6 tumor burden in CAR19^+^ CTL recipients ∼14-day post T-cell infusion (**Fig. 6D-E**). In contrast, luc-GFP^+^ CAR19^+^ CD8-iTreg displayed lower initial expansion but remained detectable *in vivo* for a longer duration, persisting up to 28-days post T-cell infusion (**Fig. 7C-D**). These data suggest that improved outcomes are driven by a sustained, moderate anti-tumor response rather than rapid, transient cytotoxicity.

**Figure 7.**
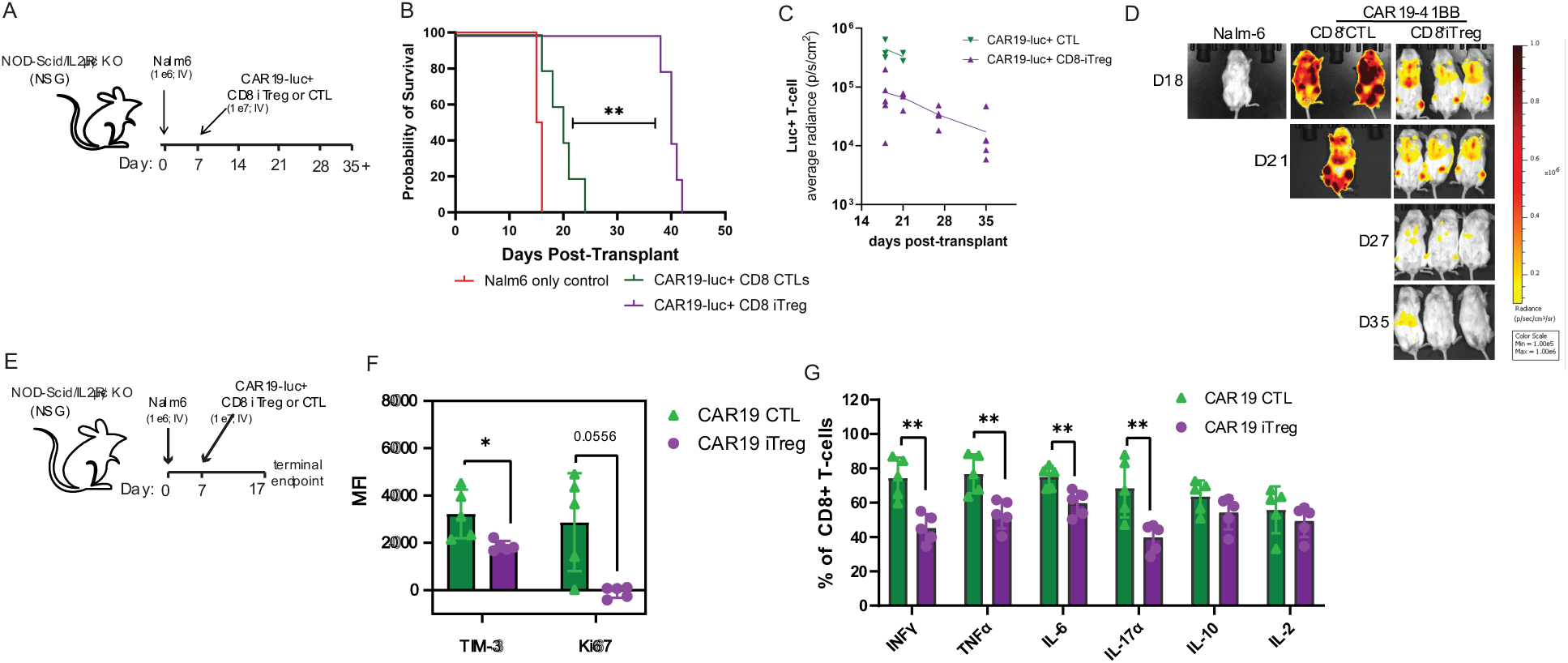
CAR19-4/1BB CD8-iTreg exhibit enhanced longevity in vivo with reduced pro-inflammatory cytokine profile. (**A**) Schematic depiction of the luc^+^ CAR19^+^ T-cell model with luc^-^Nalm-6 tumor cells and no supplemental IL-2. (**B**) Probability of survival of CAR19^+^ iTreg (purple) vs CAR19^+^ CTLs (green) vs tumor-only recipients (red), with (**C-D**) detectable frequency of CAR19^+^ iTreg vs CAR19^+^ CTLs in vivo, shown as (**C**) average radiance and (**D**) representative IVIS images. n=5/group; representative of 2 independent experiments (**E**) Schematic depiction of the luciferase positive CAR19^+^ T-cell model with luc-GFP expressing Nalm-6 tumor cells and day-17 terminal endpoint. (**F-G**) Spleens were isolated from CAR19^+^ iTreg and CAR19^+^ CTL recipient mice 10-days post-transfer into tumor bearing recipients (n=6/group). (**F**) Expression of TIM-3 and Ki67 and (**G**) frequency IFNγ, TNFα, IL-6, IL-17α, IL-10, and IL-2 expressing CD3^+^ cells isolated from recipient spleen.

Given the clinical limitations of CAR T therapy, including CRS and neurotoxicity (50, 51), our promising *in vitro* results (**Fig. 2I**), and suggestion that Treg-based therapies could be as a safer therapeutic alternative (16), we next assessed CD8-iTreg for *in vivo* inflammatory cytokine profile. Here, irradiated NSG mice received 1x10^6^ Nalm-6 tumor cells, followed by CAR19^+^ CTLs or CD8-iTreg (1×10⁷ each) on day-7, without exogenous IL-2, with terminal endpoint 10-days post-transfer (**Fig. 7E**). CAR19⁺ CD8-iTreg were confirmed to express lower levels of T-cell exhaustions marker TIM-3 and T-cell proliferation marker Ki67 vs CAR19⁺ CD8 CTLs (**Fig. 7F**). In parallel, of splenic CD3^+^ cells isolated on day 10 post-transfer were stimulated with ex vivo PMA/Ionomycin for 4hrs to evaluate cellular cytokine production. Subsequent flow-based cytokine analysis revealed that CAR19⁺ CD8-iTreg maintained a lower frequency of pro-inflammatory cytokines following *in vivo* tumor challenge, including IFNγ, TNFα, IL-6 and IL-17α, compared to CAR-CTL recipients (**Fig. 7G**); IL-2 and IL-10 expression were similar in both cell types.

### CAR19^+^ CD8-iTreg restrict leukemia progression and preserve GVHD protection

We previously reported that murine CD4^+^ Treg expressing CAR19 effectively reduced GVHD lethality without loss of GVL efficacy, in contrast to GVHD lethality seen with conventional CAR19 T-cell recipients (16). To examine the dual-functional potential of human CAR19^+^ CD8-iTreg *in vivo*, we utilized a model of residual leukemia post allo-HSCT by injecting NSG mice with CD19^+^ Nalm-6 leukemia, followed by sublethal irradiation 6 days later. On day-7, mice were co-injected with human PBMCs with or without CAR19^+^ CTLs, or CAR19^+^ CD8-iTreg, all without exogenous IL-2 (**Fig. 8A**). In the absence of CAR19^+^ T-cell intervention, PBMCs recipients had a marginal increase in MST compared to Nalm-6 leukemia recipients (**Fig. 8B**), with minimal protection against leukemia progression (**Figs. 8C-D**). CAR19^+^ CTL treatment initially produced robust antileukemic activity protecting mice from immediate fatal tumor progression (**Fig. 8B-D**), resulting in an improved MST compared to Nalm-6/PBMC recipients (29- vs 20-days, respectively; *p*<0.0026; **Fig. 8B**). Conversely, mice receiving CAR19^+^ CD8-iTreg intervention maintained significantly prolonged MST compared to CAR19^+^ CTL recipients (43-day ; p<0.0173; **Fig. 8B**), in addition to blunted leukemia progression (**Figs. 8C-D**),and reduced early weight loss compared to CAR19^+^ CTL recipients (**Fig. 8E**), suggesting that CAR19^+^ CD8-iTreg can maintain potent anti-leukemic activity following allo-HSCT, without exacerbating clinically detectable inflammatory disease.

**Figure 8:**
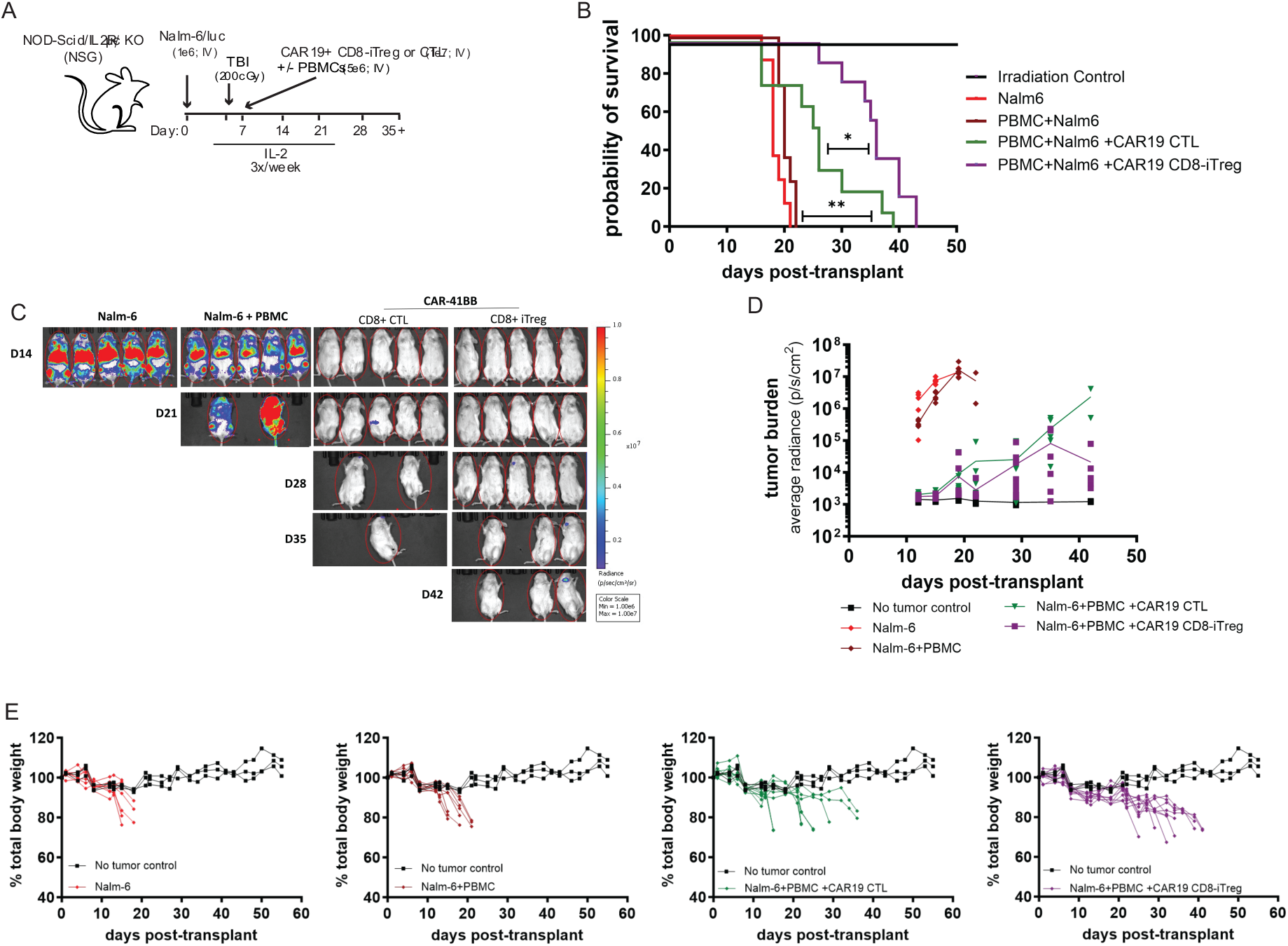
CAR19^+^ CD8-iTreg sustain anti-Tumor efficacy in a model with residual leukemia. Survival probability of survival (**B**), with representative IVIS images (**C**) and average tumor burden determined by BLI, shown as average radiance**, (D)** and total body weight (**E**) of mice receiving no tumor cells (Black) or tumor cells alone (Red), or tumor cells with PBMCs for establishing with residual leukemic (Dark Red) with or without CAR19^+^ CTL (Green) or CAR19^+^ CD8-iTreg (Purple) infusion. n=10/group; pooled from 2 independent experiments.

## DISCUSSION

Here, we developed a strategy to generate a clinically relevant human CD8-iTreg population capable of integrating potent tumoricidal activity while maintaining a restrained inflammatory signature and with robust immunoregulatory function. Single-cell RNA-seq analysis establishes CD8-iTreg as a distinct state, co-expressing canonical CTL and CD4⁺ Treg programs and differing from previously described CD8⁺ Treg populations. This challenges the prevailing view that regulatory differentiation necessarily compromises effector function.

Mechanistically, CD8-iTreg are distinct from previously reported CD8^+^ Treg populations, sharing more markers from both CTL and CD4^+^ Treg simultaneously. Notably, CD8-iTreg deployed a distinct cytotoxic program characterized by the expression and release of GzmK. We found GzmK to be an important contributor in both cytotoxic and suppressive functions. This dual role suggests a previously unrecognized mechanism through which CD8-iTreg integrate effector and regulatory pathways.

Such convergence, similar to that described previously within the context of hemophagocytic lymphohistiocytosis (HLH) pathogenesis (52), challenges the classically contrasting roles of cytotoxic and regulatory T-cell states and raises the possibility that selective deployment of alternative granzyme programs may modulate both T-cell killing and inflammatory output. However, while CRISPR/RNPs GzmK-ko support a modest dual functional role for GzmK, broad pharmaceutical dual-kinase inhibition by Bosutinib also likely contributes to the inhibition of other unidentified CD8-iTreg suppressor pathways independent of GzmK (53). Similarly, we recognized that the use of HLA-mismatched donors across *in vitro* suppression assays throughout our studies could reflect contact-dependent cytolysis of activated PBMCs rather than classical regulatory suppression. Altogether, further mechanistic studies are still necessary to comprehensively identify the core functional pathways.

Importantly, the bifunctional properties of CD8-iTreg are preserved following CAR engineering. CAR19⁺ CD8-iTreg mediate antigen-specific tumor killing while maintaining a reduced inflammatory cytokine profile. *In vivo*, these cells provide superior leukemia control and survival compared to conventional CAR19⁺ CTLs, despite reduced early expansion. Consistent with their regulatory phenotype, CAR19⁺ CD8-iTreg isolated 10-days post-transfer were found to express significantly lower pro-inflammatory cytokine levels compared to CAR19⁺ CTLs isolated from a tumor bearing recipients. While these data do support a reduced pro-inflammatory signature as observed in vitro, these data could also reflect the reduced tumor burden in CAR19^+^ CD8-iTreg recipients— thereby confounding our interpretation. Additional studies are needed to fully characterize the *in vivo* safety profile of CAR-expression CD8-iTreg. Similarly, in a model incorporating both leukemia and GVHD, CAR19⁺ CD8-iTreg again conferred a significant survival advantage relative to CAR19⁺ CTLs. Although both cell types contributed to improved leukemia control in the presence of alloreactive PBMCs, CAR19⁺ CTL recipients frequently exhibited rapid progressive weight loss and clinical signs consistent with the development of GVHD and/or proinflammatory toxicities. In contrast, animals receiving CAR19⁺ CD8-iTreg displayed both delayed mortality and GVHD-associated weight loss without evidence of recurrent tumor burden, suggesting that leukemia remained effectively controlled despite concurrent suppression of GVHD.

Collectively, these findings highlight their potential as a safer alternative to conventional CAR T cells and underscore that therapeutic efficacy depends on qualitative features—such as differentiation state, cytotoxic architecture and *in vivo* persistence—rather than maximal proliferation. This aligns with established CAR T-cell paradigms where the choice of costimulatory domains dictates quality over quantity; CD28 co-stimulation drives rapid expansion but rapid exhaustion, whereas 4/1BBis more slowly expansile but promotes durable persistence (54). Similarly, our data suggest that intrinsic qualitative programs govern the superior fitness and longevity of CD8-iTreg.

Notably, CD8-iTreg were observed to deploy a distinct cytotoxic program characterized by secretion of potent cytotoxic SMAPs, enriched with Tsp-4, Prf and multiple granzymes, including GzmK, that efficiently eliminate leukemia targets. We identified Tsp-4 as a critical scaffold that coordinates cytotoxic cargo loading and supports SMAP formation. The enhanced efficacy of SMAP-mediated killing over soluble effector proteins points to clear functional advantages of scaffolded delivery (34); packaging perforin and granzymes within these particles may maximize their concentration at the immunological synapse, improve extracellular stability, and limit diffusion. We propose that reducing Tsp-4 expression in CD8-iTreg disrupted the localization of GzmK and Prf in lytic granules, corresponding with significantly reduced cytotoxic capacity, supporting an integral role for SMAP-mediated delivery in CD8-iTreg effector function. Although the precise molecular mechanisms remain to be defined, these findings further support an emerging model in which cytotoxic lymphocyte function is critically influenced by higher-order organization of effector molecules.

Although these findings highlight CD8-iTregs as a novel population with distinct clinical potential, several limitations remain. First, while multiple established CD8+ Treg subsets exist, the scope of this analysis was restricted to the broad characterization of a heterogeneous, PB-derived CD8-iTreg population. Future studies are needed to correlate this heterogeneity with function, specifically delineating transient and terminal differentiation states, distinct subtypes, and cellular origins with function. Second, the precise mechanism(s) by which GzmK contributes to tumor control and immune regulation remain incompletely defined. Future studies will utilize proteomic and transcriptomic profiling alongside pathway-targeted and signaling reporter assays to define these downstream mechanisms. Additionally, the molecular determinants governing SMAP stability and target engagement require further investigation to determine how specific structural components of the shell modulate its assembly and kinetics. Because xenogeneic models do not fully recapitulate human immune complexity, future studies will need to evaluate CD8-iTregs in syngeneic and humanized mouse models. These immunocompetent platforms will allow us to more comprehensively assess long-term safety, chronic persistence, tissue homing, and potential off-target toxicities in a physiologically relevant environment.

From a translational perspective, CD8-iTreg represent a promising cellular platform to address the fundamental efficacy–toxicity trade-off in current immunotherapy. By coupling non-conventional cytotoxicity programs with intrinsic immune regulation, CD8-iTreg enable durable tumor control without compromising the desired safety-profile and GVHD attenuation of a Treg-based therapy. Such desirable properties make CD8 iTreg a strong cellular candidate to treat patients with active GVHD and tumor recurrence. More broadly, our findings suggest that engineering T-cells with integrated regulatory circuits—rather than simply enhancing cytotoxicity—may provide a generalizable strategy to improve the therapeutic index of T-cell-based immunotherapies.

## MATERIALS AND METHODS

### Cell purification and culture

CD8^+^CD25^-^ T-cells were enriched from non-mobilized PB apheresis products (Memorial Blood Center, St. Paul, MN, USA) using MACS technology (Miltenyi Biotec) by depleting non-CD8^+^ T-cells with cGMP-grade monoclonal antibody (mAb)-coated microbeads (cocktail of CD8, CD14 and CD19) on an AutoMACS (Depletion 2.1). Unbound cells were washed and CD25^+^ T-cells were subsequently depleted by positive selection using cGMP-grade anti-CD25 microbeads. Bead incubations and washes were carried out as specified by the manufacturer (30’ at room temperature for cGMP-grade beads).Purified cells were stimulated with a K562 cell line (KT) engineered to express CD86 and the high-affinity Fc receptor (CD64) (KT64/86) (1:1 Tcell:KT) (55), irradiated with 10,000 cGray and incubated with anti-CD3 mAb (Orthoclone OKT3; Miltenyi Biotec). Cells were cultured in X-Vivo-15 media (Lonza) supplemented with 10% human AB serum (Valley Biomedical), GlutaMAX (Gibco) and N-acetylcysteine (USP). Recombinant IL-2 (300 IU/ml, Novartis) was added on day-0 and maintained throughout the culture. For iTreg cultures, rapamycin (Tocris) at 109 nM and TGFβ (R&D Systems) at 9ng/ml were added on day-0, and with subsequent media supplementation. Cultures were maintained at 0.25–1.0×10^6^ viable nucleated cells/ml every 2–3 days (36). Complete details for flow cytometry and intracellular cytokine staining can be found in the Supporting information.

### XenoGVHD model

XenoGVHD was used as described (56–59). Briefly, NOD/Scid/γ_c_–/– mice (NSG; Jackson) mice were treated with 0.5 Gy TBI, then injected with PBMCs (2.5x10^6^) with or without CD8^+^ CTLs or iTreg (5x10^6^). Where indicated mice received endogenous rhIL-2 by I.P. injection 3x/week for 21-days following T-cell infusion. Mice were assessed daily for survival, clinical GVHD scores and weights were obtained thrice weekly.

### Xenogeneic CD19^+^ leukemia models

NSG mice were administered with CD19^+^ or CD19-KO (as indicated) luc-GFP expressing Nalm-6 tumor cells (0.5-1×10^6^, as indicated) on day-0, followed by CD8-iTreg or CTLs (5-10×10^6^, as indicated) on day-7. Tumor burden was quantified twice weekly using live total body BLI (IVIS Spectrum). For BLI, D-Luciferin (GoldBio) was reconstituted with DPBS without calcium and magnesium at 30mg/ml. Each mouse was injected with 100uL of reconstituted D-luciferin, and imaged 5 minutes post-injection. Mice were anesthetized with 1.5% Isoflurane for the duration of the imaging. Mice were imaged using the open filter, field of view 22.6 cm x 22.6 cm, *f*/stop=1, with an exposure time automatically determined by IVIS Spectrum software ranging from 4 seconds to 5 minutes. Mice were assessed for clinical GVHD and weighed thrice weekly.

For xenoGVHD with residual leukemia experiments, day-0 NSG recipients were administered with CD19^+^ Nalm-6 tumor cells (1×10^6^), followed by TBI (200rads) on day-6 and I.V. administration of human PBMNCs (2.5×10^6^) with or without CD8-iTreg or CTLs (4-5×10^6^, as indicated) on day-7.

### Study approval

Animal studies were conducted in accordance with a protocol reviewed and approved by the IACUC of the University of Minnesota (protocol# 2312A41590) Human cells utilized for *in vitro* and *in vivo* studies were derived from leukapheresis products purchased from the Memorial Blood Center (St. Paul, MN). No approval was required for these purchased human samples.

### Statistical Analyses

Data are reported as mean values ±SEM. Pairs were compared using unpaired 2-tailed Student’s t-tests. Data sets with ≥3 samples were compared using 1-way ANOVA with multiple comparisons analysis. Data were analyzed by ANOVA or student’s t-test. Probability (p) values ≤0.05 were considered statistically significant. Differences in survival were analyzed by log-rank test. Significance was defined as p<0.05. Statistical analyses were performed using PrismV9 (GraphPad).

## Supporting information

Supplemental Results and Methods

## Abbreviations

ALL: acute lymphoblastic leukemia
Allo-HSCT: allogeneic hematopoietic stem cell transplantation
AML: acute myeloid leukemia
BiTE: bispecific T-cell engager
BLI: bioluminescent imaging
CAR: chimeric antigen receptor
CAR19: CD19-targeting scFv based chimeric antigen receptor
CD8-iTreg: human CD8+ induced Regulatory T-cell
CFSE: Carboxyfluorescein N-succinimidyl ester
CMA: Concanamycin A
CTL: cytotoxic T-cell
CTLA-4: cytotoxic T-lymphocyte-associated protein 4
CRS: cytokine release syndrome
cryo-SXT: cryo-Soft X-ray Tomography
CXCR1: C-X-C motif chemokine receptor 1
dSTORM: direct Stochastic Optical Reconstruction Microscopy
DCI: 3,4-dichloroisocoumarin
DEG: differentially expressed genes
EM: electron microscopy
EGFR: epidermal growth factor receptor
FACS: fluorescence-activated cell sorting
FasL: Fas ligand
FCS: flow cytometry standard
GFP: green fluorescent protein
GI: gastrointestinal tract
GITR: glucocorticoid-Induced TNFR-Related protein
GVL: graft-versus-leukemia
GVHD: graft-versus-host disease
GzmA: granzyme A
GzmB: granzyme-B
GzmK: granzyme-K
GzmM: granzyme-M
GNLY: granulysin
HLA: haploinsufficiency leukocyte antigen
HSCT: hematopoietic stem cell transplantation
ICAM-1: intercellular Adhesion Molecule 1
IFNγ: interferon gamma
IL-6: interleukin 6
IL-10: interleukin 10
i.p.: intraperitoneally
iTreg: induced regulatory T-cell
i.v.: intravenously
IVIS: in vitro imaging system
KIR: killer-cell immunoglobulin-like receptor
KLRG1: killer cell lectin-like receptor G1
KO: knock-out
LAG3: lymphocyte-activation gene 3
LAMP-1: lysosomal-associated membrane protein 1
luc: luciferase
MFI: mean florescence intensity
MHC: major histocompatibility complex
MOI: multiplicity of infection
MST: median survival time
NSG: NOD *scid* gamma
nTreg: natural regulatory T-cells
PB: peripheral blood
PBMNCs: peripheral blood mononuclear cells
Prf: perforin-1
SE: Superecliptic
pSLB: planar supported lipid bilayers
SMAP: supramolecular attack particle
TBI: total body irradiation
Teff: effector T-cells
TGFβ: transforming growth factor beta
THBS: Thrombospondin gene
Tsp-1: thrombospondin-1 protein
Tsp-4: thrombospondin-4 protein
TIGIT: T-cell immunoreceptor with immunoglobulin and ITIM
TNFα: tumor necrosis factor alpha
Treg: regulatory T-cell
Tsp: Thrombospondin protein
tTreg: thymic derived regulatory T-cell
WGA: wheat germin agglutinin

## Data Availability Statement

Datasets generated and analyzed during the present study are available from the corresponding author on reasonable request.

## Acknowledgments

The authors gratefully acknowledge the University of Minnesota University Imaging Centers (UIC) SCR_020997 for their support in imaging analysis, and the staff at the University of Minnesota University Flow Cytometry Facility for cell sorting. Matthew Wood for access to their Oxford Nanoimager (ONI), Jonathan Webber at the Kennedy Institute of Rheumatology Flow Cytometry Facility, and Felix Clanchy and Richard Williams to provide acellular supernatant from RA-patient synovial fibroblasts. We acknowledge Diamond Light Source for time on B24 Cryo Soft X-ray Tomography under proposal BI-32122 (to MLD) and BI-30452 (to EBC).

## Funding Statement

This work was supported by grants from the Children’s Cancer Research Fund, the National Institutes of Health (R01 HL11879; P01 CA065493) and the Hematology Research Training Program T32 HL007062. This work was also supported by ERC Synergy grant 951329 (ATTACK) to MLD and CTB.

## Author Contributions

JHL, EBC, KH, MD, and BRB conceived and designed the experiments; JHL, EBC, KS, PRD, OM, MHT, NCP, NC, CC, BK and BH performed the experiments; JHL, EBC, MCZ, KS, OM, SJ, PRD, CMH, SBW, MHT, MD, KH and BRB, analyzed and interpreted the data; BRB, KH, and MD contributed reagents/materials; and JHL, EBC, KS, PRD, MCZ, OM, SJ, CTB, MD, KH and BRB contributed to the writing and editing of this manuscript. All authors approved the manuscript.

## Competing Interest

B.R.B. receives remuneration as an advisor to Magenta Therapeutics and BlueRock Therapeutics; research funding from BlueRock Therapeutics, Rheos Medicines, Equilibre biopharmaceuticals, and Carisma Therapeutics, Inc.; and is a co-founder of Tmunity Therapeutics. M.L.D. receives remuneration as an advisor to Labgenius. All other authors declare that they have no competing interests.

## SUPPLEMENTAL FIGURES

**Supplemental Figure 1.**
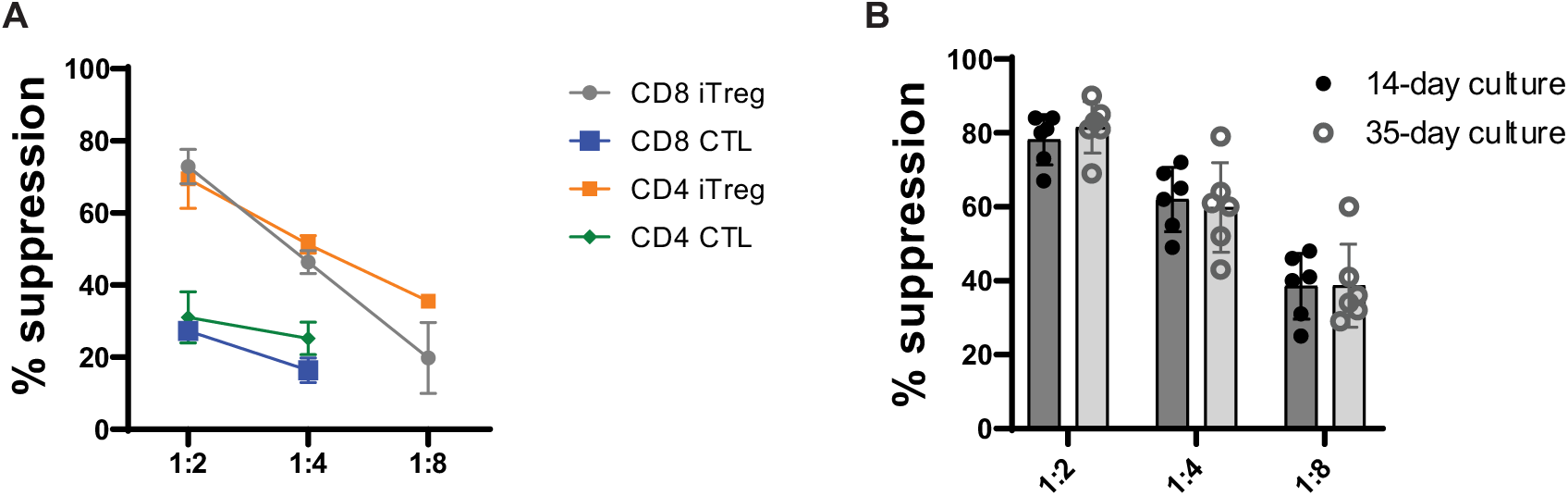
**(A)** In vitro suppression of activated PBMC proliferation by donor-matched CD8-iTreg vs CD8-CTLs vs CD4^+^ iTreg vs CD4^+^ CTLs. CD8 and CD4 iTreg and CTLs were generated from donor-matched PB CD8^+^CD25^-^ or CD4+CD25^-^ T-cells, respectively, stimulated for 14-days with either IL-2 alone or IL-2, Rapamycin, and TGFβ. **(B)** In vitro suppression of activated PBMC proliferation by 14-day vs 35-day cultured CD8-iTreg at different effector-to-PBMC ratios (1:2, 1:4, 1:8) vs PBMC-only controls (0:1).

**Supplemental Table 1.**
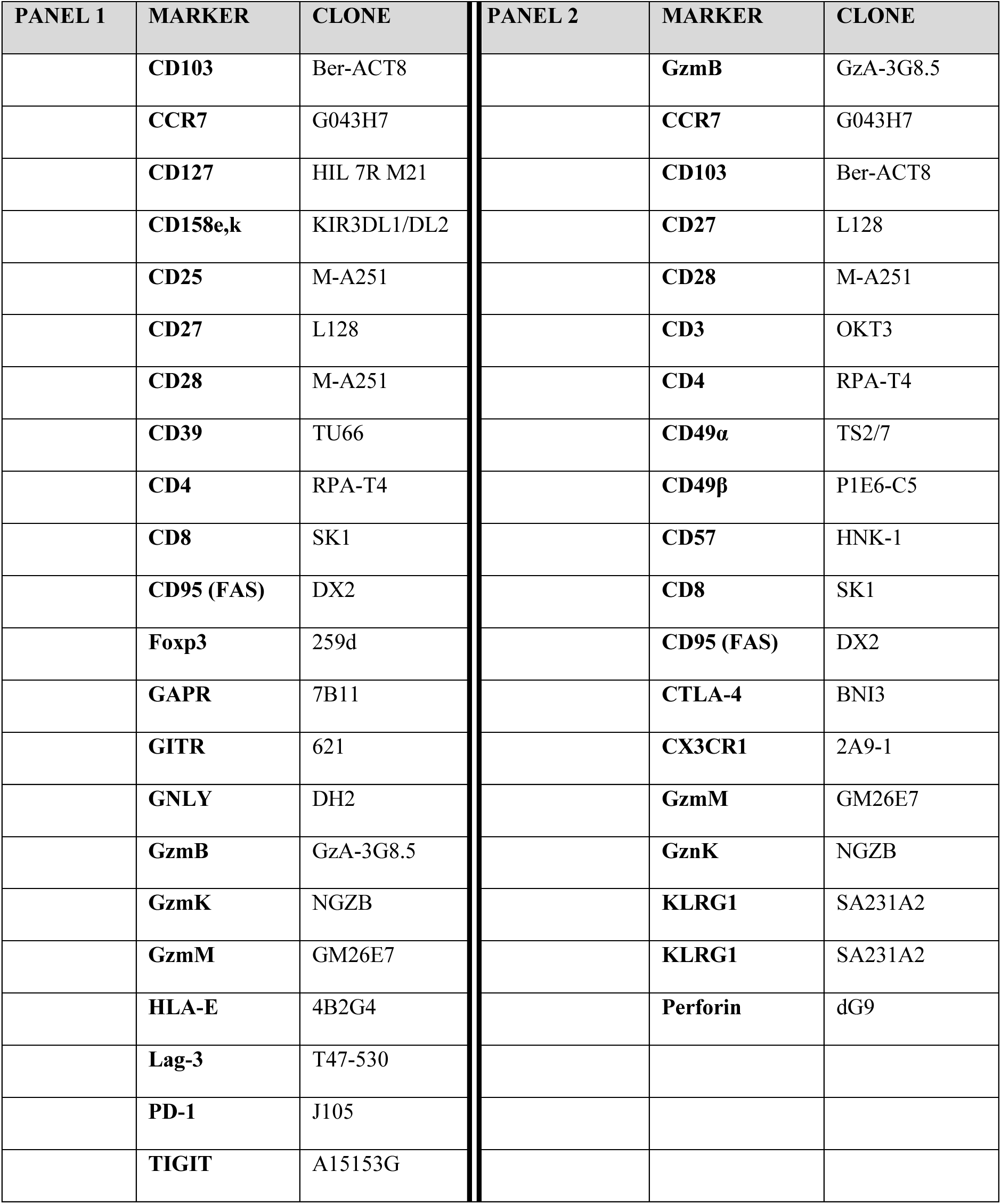
Comprehensive 32-Parameter Spectral Flow Cytometry Panels. List of markers analyzed across two experimental panels evaluating T-cell activation, cytotoxicity, immunosuppression, and canonical CD4^+^ and CD8^+^ nTreg phenotypes.

**Supplemental Figure 2.**
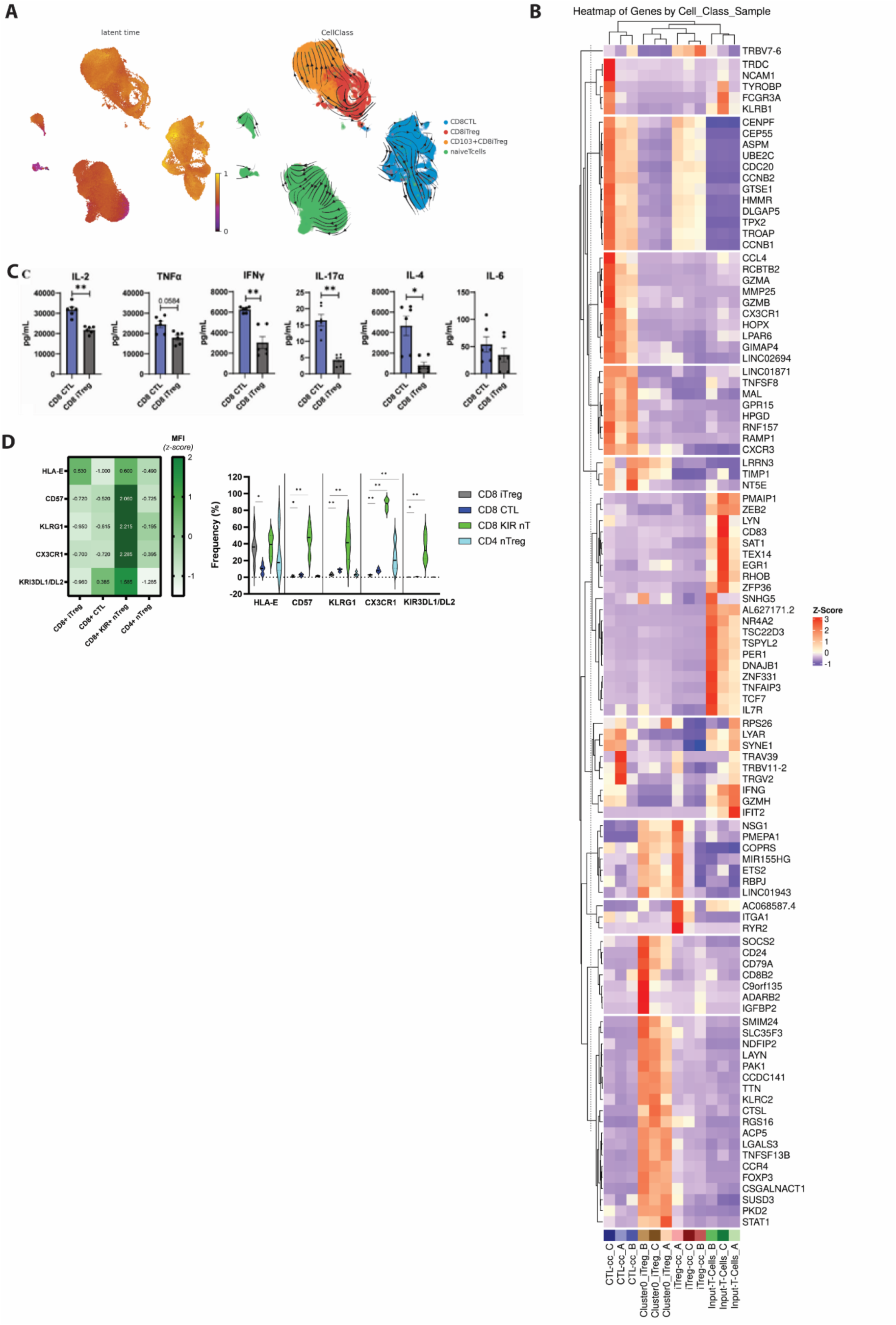
(**A**) Results of scVelo RNA velocity analysis. (Left) Calculated cellular latent time heatmap. Earliest latent time point appear in purple in naïve T-cell associated macroclusters. Latest latent time point appear in CD103^+^ CD8iTreg and CTL macroclusters in yellow. (Right) RNA velocity stream visualization generated from scVelo. (**B**) Heatmap of top 25 DEGs identified for the four major cell types: CD103^hi^ CD8-iTreg (orange) vs CD103^lo^ CD8-iTreg (red) vs CTLs (blue) vs CD8^+^CD25^-^ input cells (green). (**C**) Total cytokine production (IL-2, TNF⍺, INFɣ, IL-17⍺, IL-4, and IL-6) from CD8 CTLs (dark blue) vs CD8 iTreg (grey) after 4hr stimulation with PMA/Ionomycin, quantified by RayPlex® Human Inflammation Bead Array. **(D)** Relative expression and population frequency of KIR^+^ CD8 Treg associated markers in CD8-iTreg compared to donor-matched, CD8^+^ CTLs and hCD4^+^ Treg (light blue).

**Supplemental Figure 3.**
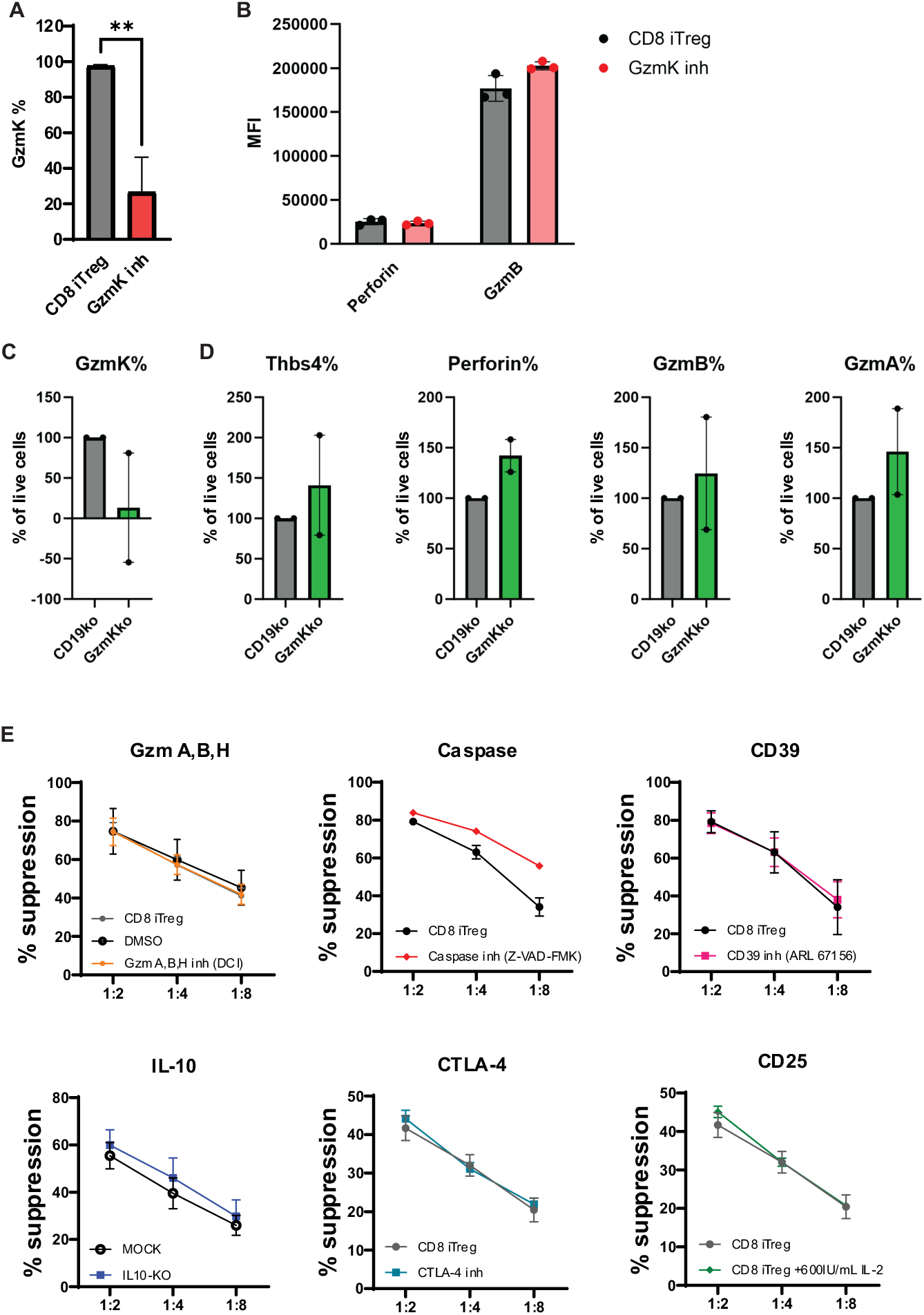
**(A)** Frequency ±SD of GzmK^+^ CD8-iTreg following treatment with 10μM of Bosutinib for 48hrs vs untreated control (n=3 biological replicates). **(B)** Expression of Perforin and Granzyme B in CD8-iTreg after 48hr treatment with 10uM Bosutinib (n=3 biological replicates). Frequency of **(C)** GzmK, and **(D)** Thbs4, Perforin, GzmB and GzmA in CD8-iTreg electroporated with GzmK CRISPR/Cas9 RNPs vs irreverent CD19-ko RNPs (n=2 biological replicates). (**E**) In vitro suppression of activated PBMC proliferation at different effector-to-PBMC ratios (1:2, 1:4, 1:8) vs PBMC-only controls (0:1) by CD8-iTreg (grey) vs CD8-iTreg following inhibition or blockade of GzmA/B/H, Caspase, CD39, IL-10, CTLA-4, or CD25, respectively; shown as mean ±SD for each ratio (n=3 biological replicates).

**Supplemental Figure 4.**
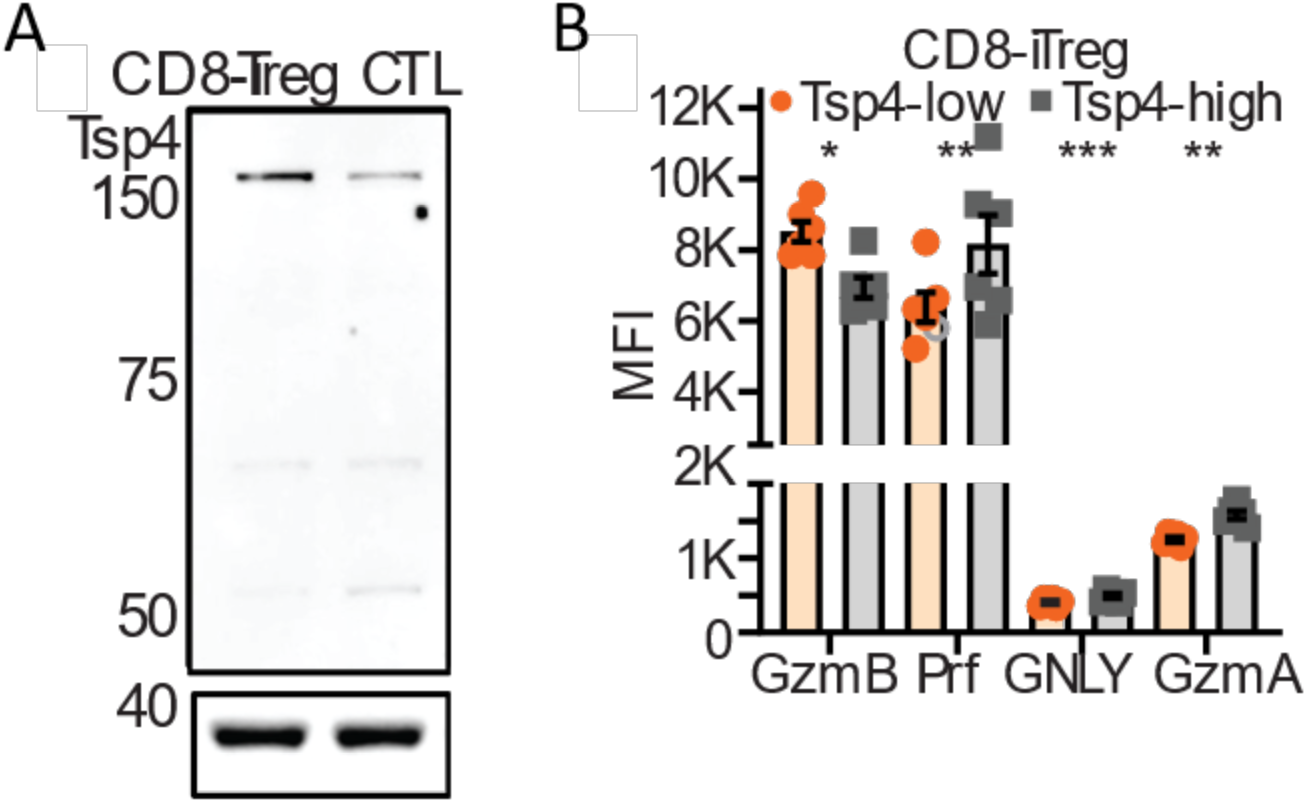
**(A)** Western blot analysis of Thrombospondin-4 expression in CD8-iTreg and donor-matched CTLSMAPs, released on activating Quantifoil grids, coated with b-actin as loading control. (**B**) Flow cytometry analysis reporting mean fluorescent intensity (MFI)poly-l-lysine, anti-CD3ε (ΟΚΤ3) and recombinant human ICAM-1-Fc/CD54; shown as mean ±SD (n=6 biological replicates). GzmB-mCherry-pHluorin mRNA-transfected CD8-iTreg were incubated on the activating Quantifoil grids for 90 min, followed by cell flushing of GzmB, GzmA, perforinthe cells out of the activating grids with ice cold PBS, and GNLY in plunge-freezing of the sample in liquid ethane.

**Supplemental Figure 5.**
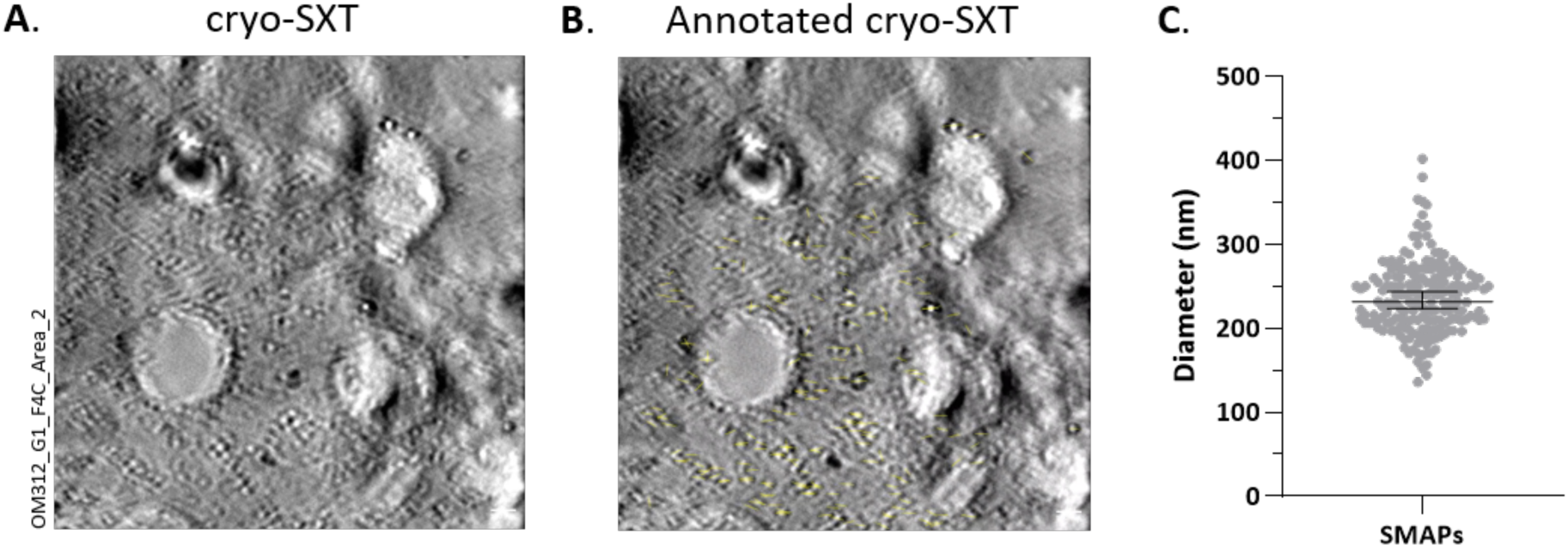
**(A**) Manually reconstructed cryo-tomogram of CD8-iTreg SMAPs, released on activating Quantifoil grids, coated with poly-l-lysine, anti-CD3ε (ΟΚΤ3) and recombinant human ICAM-1-Fc/CD54. GzmB-mCherry-SEpHluorin mRNA-transfected CD8-iTreg were incubated on the activating Quantifoil grids for 90 min, followed by cell flushing of the cells out of the activating grids with ice cold PBS, and plunge-freezing of the sample in liquid ethane. Imaging by correlative cryo-Structured Illumination Microscopy (cryo-SIM) with cryo-SXT followed under liquid nitrogen. Tomogram generation was performed in IMOD, with a manual fiducial model and Simultaneous Iterative Reconstruction Technique (SIRT) tomogram generation method with 12 iterations. (**B**) Annotations of the CD8-iTreg released SMAP diameters. (**C**) SMAPs released on activating Quantifoil grids, CD8-iTreg SMAP diameters by cryo-SXT.

**Supplementary Figure 6.**
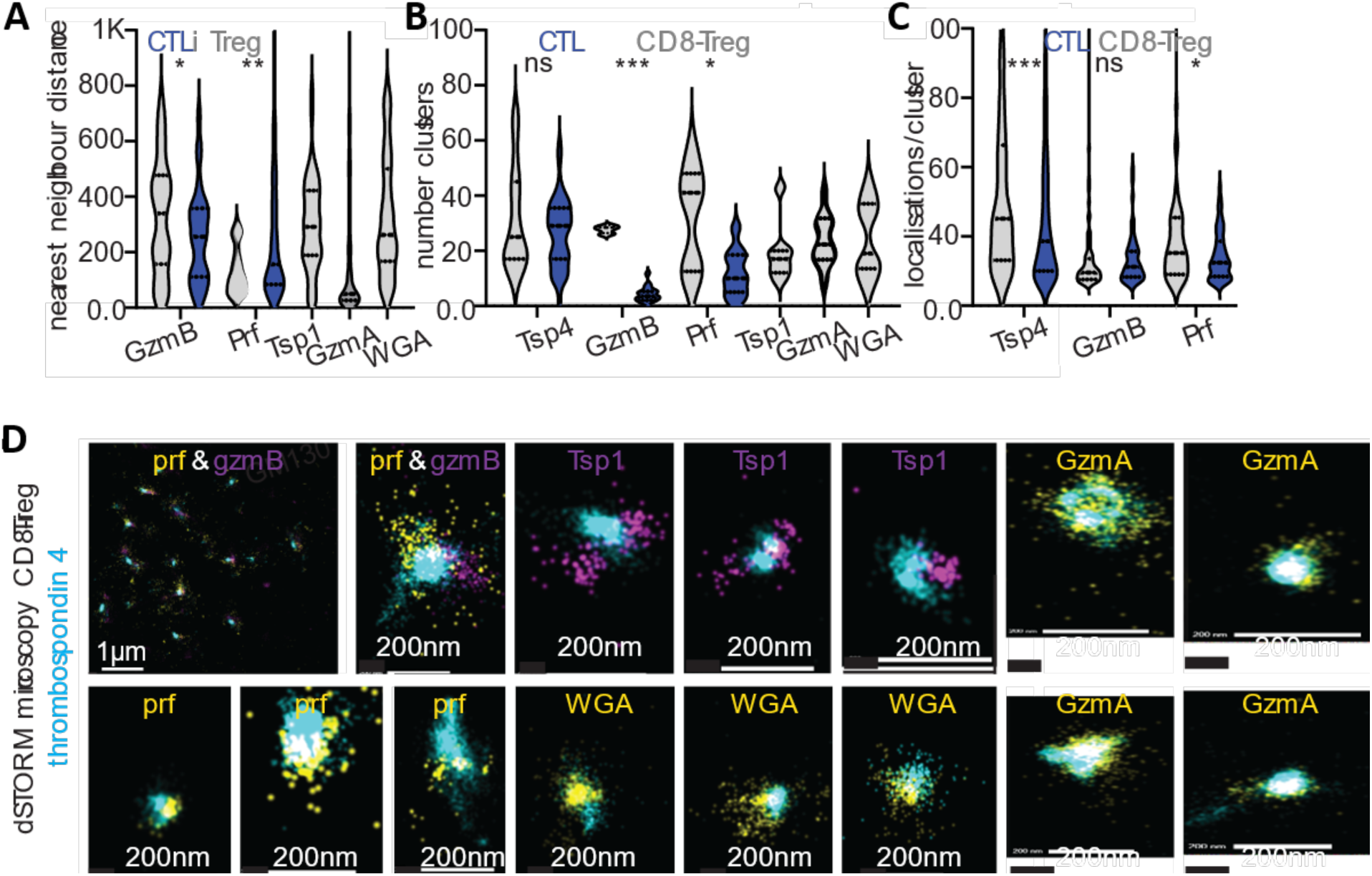
dSTORM analysis of SMAPs. Two- or three-colour dSTORM datasets were analysed using the Oxford NanoImager (ONI) CODI platform. Localization, acquisition, and mapping files were processed with drift correction (DME algorithm; TetraSpeck bead reference). Frames were filtered to remove initial excitation peaks, retaining steady-state blinking events. Localizations were filtered by Gaussian sigma (∼50–250 nm), photon counts (∼300–500), and precision (≤20 nm). SMAP radius and localization number were quantified using clustering within regions of interest at the c-SMAC. Analyses included (**A**) nearest-neighbour distance, (**B**) cluster number (DBSCAN), (**C**) localizations per cluster, and (**D**) representative images with scale bars.

**Supplemental Figure 7.**
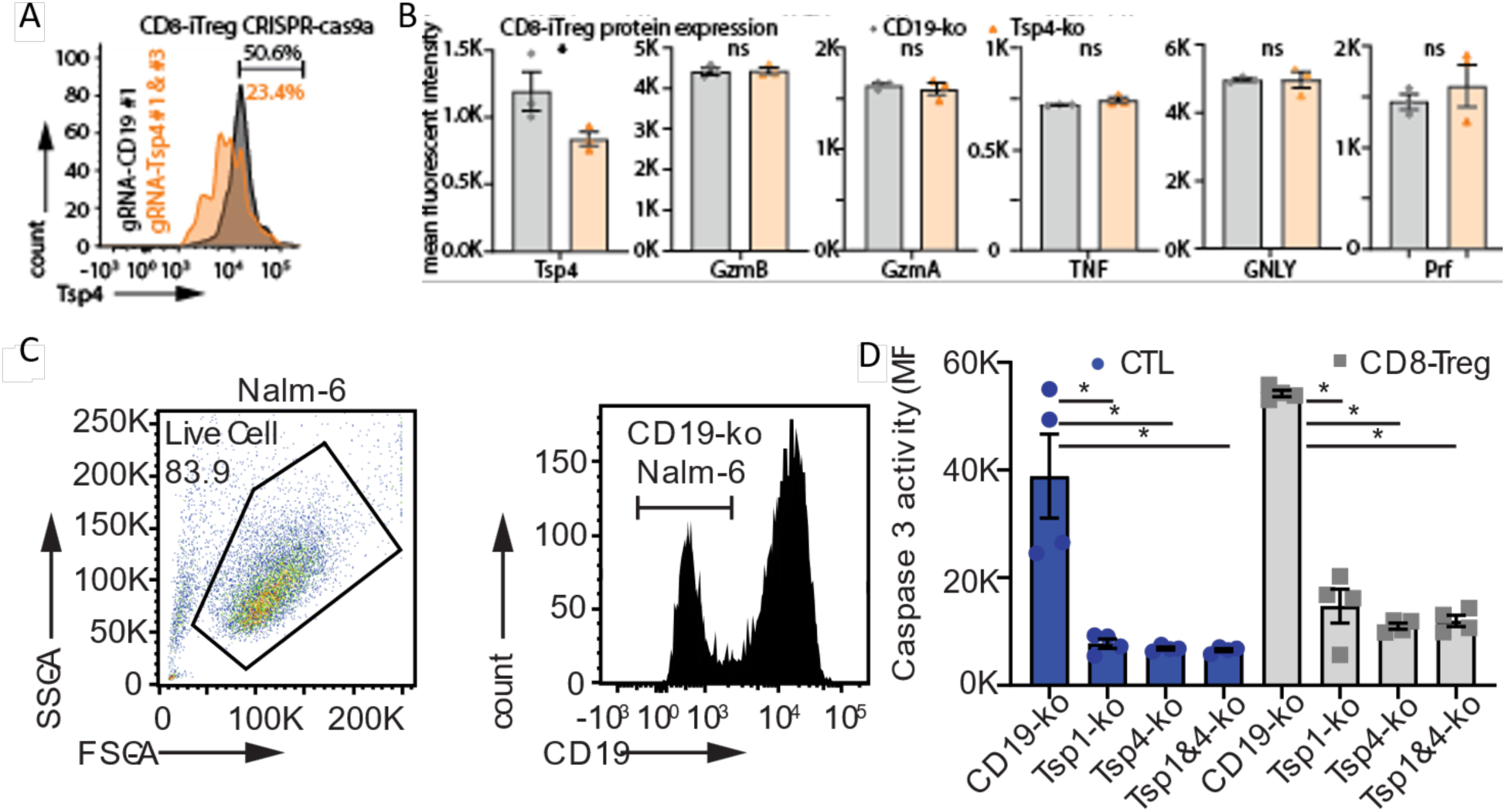
Electroporation of CD8-iTreg with CRISPR/Cas9a Ribonucleoprotein complexes (RNPs) targeting (**A**) Thrombospondin-4 with 2 different gRNAs 24hrs apart, significantly reduces Tsp-4 expression, (**B**) without altering GzmB, A, Prf, TNF, and GNLY expression, compared to once targeted with control CD19 targeting RNP; shown as mean ±SD (n=2 biological replicates). (**C**) CD19 negative Nalm-6 were generated by hitting them once with CD19-RNP, followed by sorting of CD19 negative population. (**D**) Targeting Tsp-1, Tsp-4 or both, in CAR19^+^CD8-iTreg significantly reduces their tumoricidal activity as determined by caspase 3 activity in CD19^+^ Nalm6 4hrs post co-culture.

**Supplemental Figure 8.**
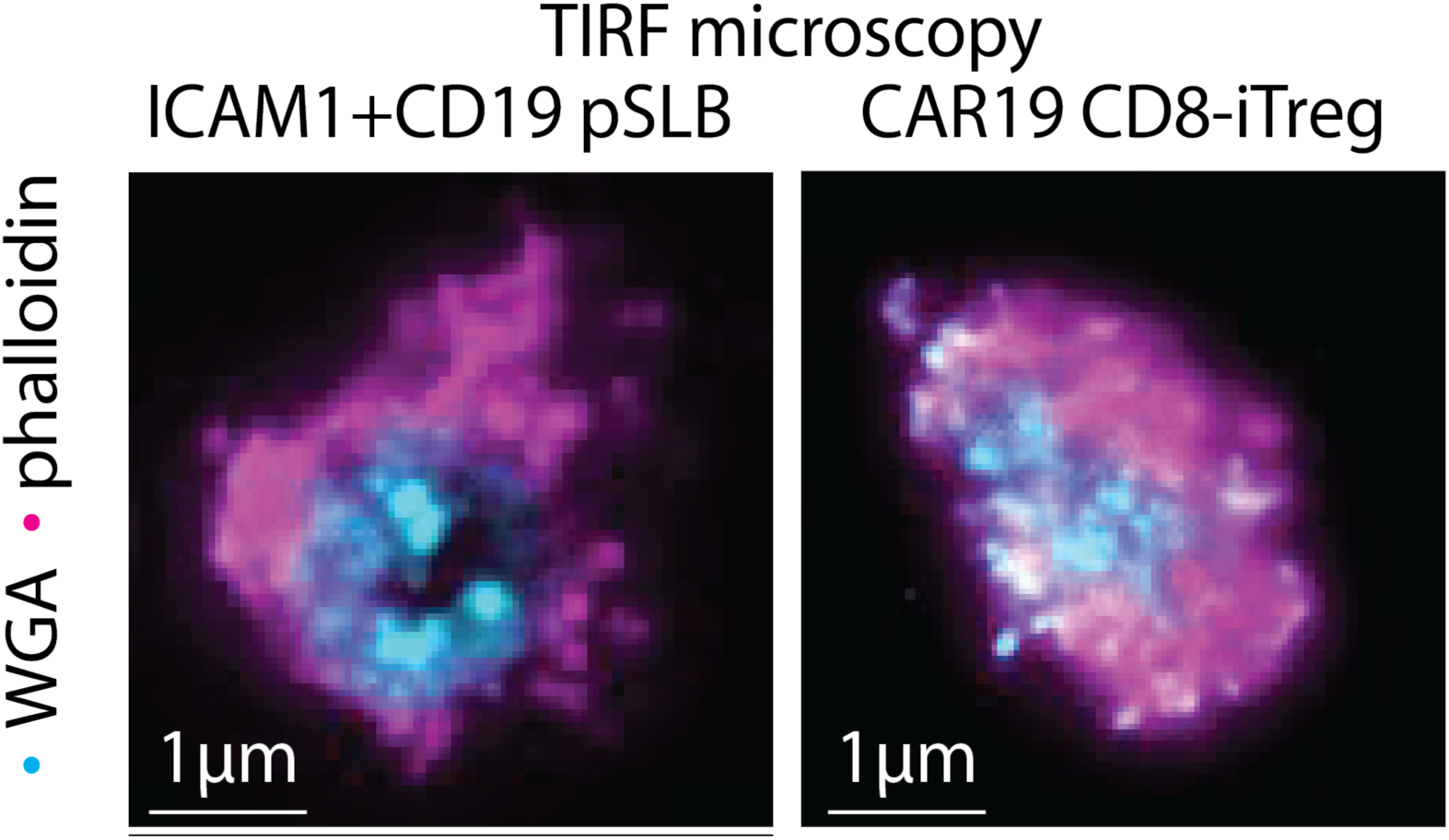
CAR19^+^CD8-iTreg form immunological synapses on planar supported lipid bilayers with his-CD19 and ICAM-1-12his, as determined by actin rings visualized with fluorescent phalloidin and synapse-released glycoproteins by wheat-germ agglutinin.

**Supplemental Figure 9.**
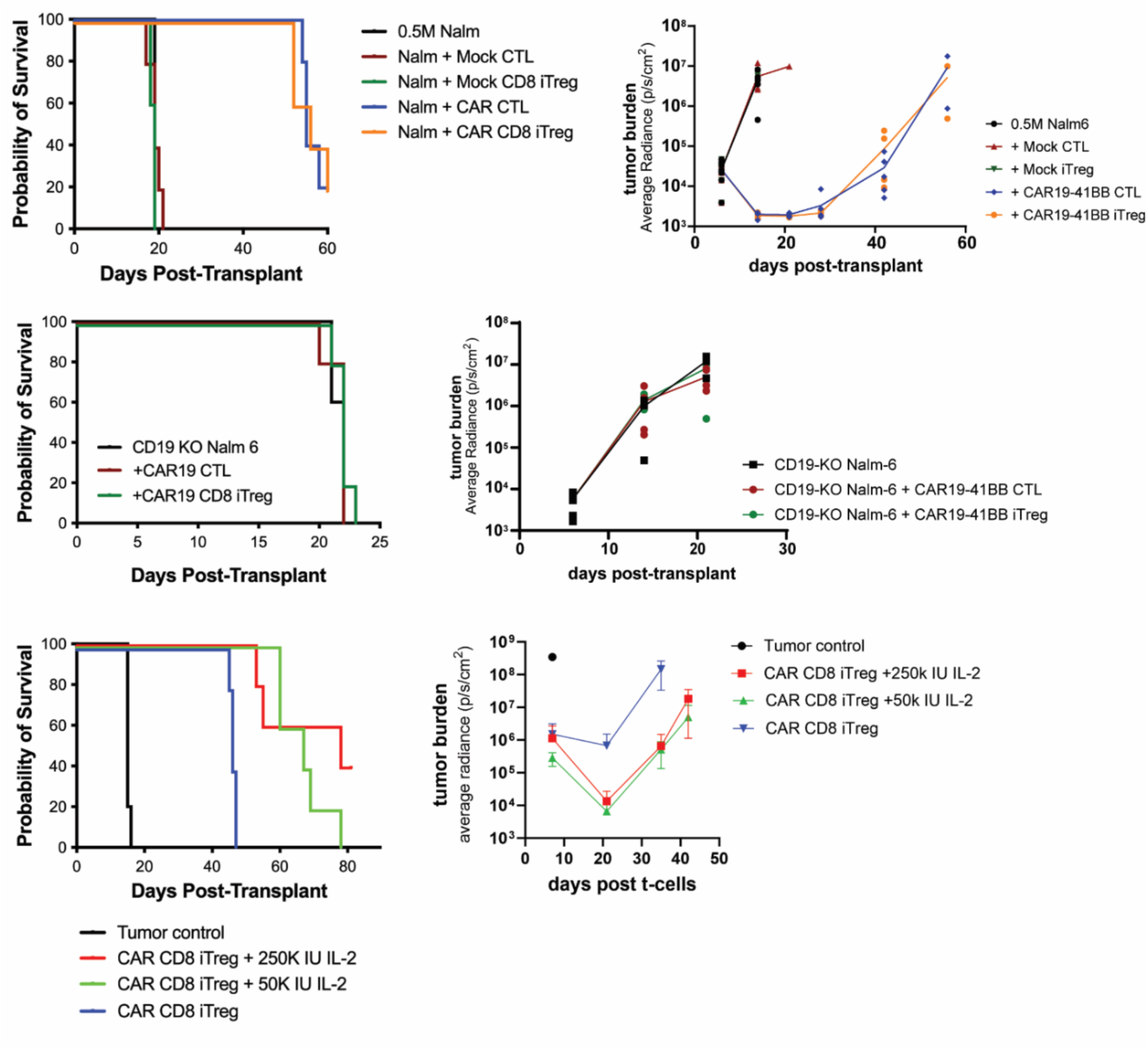
Validation of Nalm-6 xenoGVL model **(A)** Nalm-6 survival and BLI for model infusing 5x10^5^ Nalm-6 tumor D0, followed for CAR19^+^ CD8-iTreg or CD8-CTLs (10^7^/mouse) on D7 with supplemental IL-2 (250kIU/mouse; 3x/week for 21-days). (**B**) Nalm-6 survival and BLI for model infusing 1x10^6^ CD19-KO Nalm-6 tumor targets, followed for CAR19^+^ CD8-iTreg or CD8-CTLs (10^7^/mouse) on D7 with supplemental IL-2 (250kIU/mouse; 3x/week for 21-days). (**C**) Nalm-6 survival and BLI for model infusing 1x10^6^ Nalm-6 tumor targets on D0, followed by D7 infusion of CAR19^+^ CD8-iTreg (10^7^/mouse) with mice receiving 250k IU/mouse IL-2 vs 50k IU/mouse IL-2 vs vehicle control 3x/week for 21-days.

## SUPPLEMENTAL MATERIALS AND METHODS

### Flow cytometry

Single-cell suspensions were recorded on a BD LSRFortessa flow cytometer. Multiparameter spectral flow cytometry was recorded on a Cytek Aurora. FCS3.0 files were analyzed using FlowJoV10.9 (Treestar Inc.). Anti-human antibodies were sourced as follows: from BioLegend, anti-Foxp3 (259d), CD25 (M-A251), CD103 (Ber-ACT8), CTLA-4 (BNI3), GITR (621), TIGIT (A15153G), CD8 (SK1), CD4 (A161A1), GzmK (GM26E7), GNLY (DH2), Perforin (dG9), IL-1β (JK1B-1), TNFα (MAb11), IFNγ (4S.B3), IL-10 (JES3-9D7), IL-6 (MQ2-13A5), CD19 (HIB19), ICOS (15F9), EGFR (AY13), GARP (7B11 and LRRC32)Tim-3 (F38-2E2), CD95 (DX2), GARP (7B11 and LRRC32), CD158b/j (KIR2DL2/L3/S2), CD158e1 (KIR3DL1; DX9), CD158b (KIR2DL2/L3; DX27), CCR7 (G043H7), CD49α (TS2/7), CD49β (P1E6-C5), CD57 (HNK-1), KLRG1 (SA231A2), CX3CR1 (2A9-1), and CD158f (UP-R1). From BD Biosciences, anti-CD39 (TU66), CD127 (HIL 7R M21), CD28 (M-A251), CD27 (L128), Lag-3 (T47-530), GzmM (4B2G4), CD4 (RPA-T4), CD8 (RPA-T8), CD25 (M-A251), and CD45RA (HI100). Anti-GzmA (GzA-3G8.5), GzmB (NGZB), and CD3 (OKT3) were purchased from Thermo Fisher Scientific. Additional markers included human KIR3DL2/CD158k from R&D Systems and CD158e/k (KIR3DL1/DL2) from Miltenyi Biotec.

Cells were stained with either Zombie NIR Fixable Viability Kit (BioLegend) or Live/dead fixable yellow dead cell stain kit (ThermoFisher) for all experiments. Fixation and intracellular/intranuclear staining was achieved by either fixing cell samples with 4% PFA for 20mins and subsequently permeabilizing with 0.1% /Triton X-100 for 5mins prior to intranuclear staining or using the eBioscience Foxp3 staining kit for transcription factor and cytokine analysis.

### Suppression assays

*In vitro* suppressive capacity of expanded cultures was assessed with either Celltrace Far Red or Celltrace CFSE Cell Proliferation Kits (ThermoFisher) (40). PB mononuclear cells (PBMNC) were purified, labeled with CFSE (InVitrogen) and stimulated with anti-CD3/CD28 mAb-coated beads (Dynal) ±cultured iTreg, or CTL (effector-to-PBMNC at ratios of 1:2–1:32), without adjustment for Foxp3 content in the product. On day-3, cells were stained with anti-CD8^+^ and anti-CD4 mAb. Acquired data were analyzed using the proliferation platform in FlowJo, and suppression determined from the Division Index. No attempt was made to HLA match PBMNC with Treg product.

### Killing assays

*In vitro* tumor killing capacity of expanded cultures was assessed by co-culturing CD19^+^ and/or CD19-KO Nalm-6 tumor cells with CD8-iTreg or CD8-CTLs (5:1 effector-to-tumor ratio). To determine total killing capacity, on day-3, cells were stained with anti-CD3 mAb and yellow dead cell dye (ThermoScientific). Acquired data were analyzed using FlowJo, and the frequency of dead Nalm-6 tumor cells normalized for spontaneous death was quantified. Continuous Incucyte killing assays were performed by co-culturing Nalm-6 and Teff cells (as above) on Poly-D-Lysine (ThermoScientific) coated flat-bottom 96-well plates in complete RPMI media without phenol red (Gibco). Images were captured and quantified every 2hrs over 48-72hrs. Tumor cell number was normalized for spontaneous death on tumor control.

### Human single cell RNA sequencing Sample Preparation and Analysis

CD8^+^CD25^-^ T-cells were enriched from non-mobilized PB apheresis products and differentiated into either CD8+ CTLs or CD8-iTreg as described above. Triplicate biological samples were Hashtagged with TotalSeq-C0251 antibodies (BioLegend) before being pooled. Replicate pools were prepared from CD8^+^CD25^-^ T-cell inputs as well as from both CD8^+^ CTL and CD8-iTreg outputs. Samples were prepared with MOJO dead-cell remover to improve viability before encapsulation with 10x Chromium. A 5’ gene expression and CITEseq(60) library were prepared before sequencing on the Illumina NovaSeq X platform to a read depth of 2.5 billion reads. Sequencing quality control, alignment to the GRCh38-2024-A genome(61), UMI counting, and gene expression matrix generation were performed using CellRanger v9.0.1(62). CITE-seq Count(63) was used to process hashtag oligo library for demultiplexing. Samples were deconvoluted using a consensus majority-vote assessment via HTODemux, GMM-Demux(64), MULTI-seq(65), and BFF-Cluster(66) before data was loaded into Seurat v5(67) for quality control filtering, and data normalization. In Seurat dimensional reduction and universal manifold projection was performed and Louvain algorithm was used for clustering which revealed the CD103^+^ CD8iTreg subset. In addition to Seurat biomarker discovery method cells were pseudobulked by cell class and biological replicate for differential expression analysis via DESeq2(68) and gene set enrichment analysis against the Gene Ontology (GO)(69), KEGG(70), MsigDB Hallmark(71), and MsigDB Immunological signatures databases using clusterProfiler(72). For RNA Velocity and trajectory inference Velocyto(73) was used to separate spliced from unspliced gene matrices and scVelo(74) was used to determine decay kinetics and splicing ratios. Finally, CellRank(75) and Palantir(76) were used to determine terminal populations and predict lineage specific pseudotime from developmental trajectories. Analysis code available upon request to the author.

### mRNA Production and CD8-iTreg Electroporation for Cryo-CLXT

Single clones of bacterial cultures containing the pGzmb-mCherry-SEpHluorin plasmid (kindly gifted by Prof. Salvatore Valitutti Lab, INSERM), were expanded, and pGzmb-mCherry-SEpHluorin plasmids were isolated using either by using QIAprep Spin Miniprep Kit (Qiagen) or PureLink HiPure Plasmid Filter Maxiprep Kit (Invitrogen) as per manufacturer’s protocol. The plasmid DNA was linearized by Notl-HF (NEB) digestion, and the linearized DNA was used to produce Gzmb-mCherry-SEpHluorin mRNA using the mMESSAGE mMACHINE T7 Ultra Transcription Kit (Invitrogen) as per manufacturer’s protocol.

Generated mRNA was electroporated to the CD8-iTreg using ECM 830 Square Wave Electroporation System (BTX). In brief, the cells were washed thoroughly with OptiMEM Reduced Serum Medium (Gibco), and 1 x10^6^ cells were resuspended in 100μl OptiMEM with added 5μg mRNA per reaction. The mixture would be transferred to a 2mm electrode gap electroporation cuvettes (BTX), and the sample would be electroporated at the ECM 830 at 300V, for 2ms, with 1 pulse. The electroporated cells were immediately topped up with 20% FBS R10, transferred to a tissue culture plate and recovery at 37°C, 5% CO_2_ followed. Transfection efficiency was tested by FACS at Fortessa X-20.

### Sample Preparation for CD8-iTreg SMAP Characterization by CLXT

CD8-iTreg, cultured and 2-3 days after thawing, were transfected with 5 μg per 10^6^ cells GzmB-mCherrySEpHLuorin mRNA, by electroporation, as previously described. After 2 recovery days (4-5 days after thawing) they were used for CLXT at the B24 Beamline, Diamond Light Source (DLS, UK’s National Synchrotron Facility).

In brief R2/2 G200F1 (Quantifoil) grids were coated by 0.01% Poly-L-Lysine (PLL), for 30 min, and after washing with sterile water (Thermo Scientific), they were coated by combinations of 2.5μg/ml hICAM-1/CD54-Fc (R&D Systems), 5μg/ml purified anti-human CD3ε (OKT3) (BioLegend), before washing with PBS. CD8-iTreg were pre-labelled for 2hrs with 10μg/ml WGA-Alexa Fluor 647 and for 20 min with 150nM LysoTracker Blue (Invitrogen) and concentrated in 50μl conditioned medium grid per condition. The grids were placed in 2-well co-culture µ-slides (ibidi), fitting 1 grid per well, and the concentrated cells were deposited on them. For released particle imaging, after 90 min incubation, the cells were flushed out of the grids with 4°C PBS. Plunge-freezing in liquid nitrogen (LN)-cooled liquid ethane with a LeicaEMGP2 plunge freezer followed, as previously described(77), and the frozen samples were stored in LN.

### Cryo-Structured Illumination Microscopy (Cryo-SIM) at B24

A custom 3D-cryo-SIM system for the B24 beamline was used for plunge-frozen sample fluorescence imaging, performed as previously described (78, 79). In brief, the samples were loaded on a cryo-SIM microscope-attached cryo-stage (Linkam Scientific), and after the sample was located using the Cockpit Software (Micron Oxford), a bright-field mosaic of the grid would be collected. Regions of interest (ROIs) were identified based on cell morphology and fluorescence, and their locations were marked and stitched to the mosaic with StitchM (DLS). The data were collected in pairs based on the microscope camera set-up, since the emissions from the 405 nm and 488 nm lasers are both collected from camera 1, and emissions from the 561 nm and 647 nm lasers are both collected from camera 2(78). For each laser fluorescence emission, there were collected 4 different phase shifts of the sinusoidal pattern, at 3 different orientations, repeated for every Z position in a full 3D stack, using a nematic liquid-crystal-on-silicon spatial light modulator (SLM, Meadowlark optics, 512 x512 SLM Model PDM512), enabling grayscale phase modulation(78, 79). The data were reconstructed by the beamline stuff using a tailored 3D SIM reconstruction software, softWoRx (GE Healthcare). Chromatic shift correction would be performed using TetraSpeck-generated reference files, with Chromagnon V.089.

### Cryo-Soft-X-ray Tomography (Cryo-SXT) at B24

Tilted-series of X-ray projections were collected using a synchrotron endstation UltraXRM-S/L220c X-ray microscope (Carl Zeiss X-ray Microscopy, Inc.) installed at the B24 beamline, DLS, imaging by absorption contrast at the “water window” of the X-ray spectrum (46), at 500 eV with a 25 nm zone plate (Xradia), and a Pixis 1024 B charge-coupled device (CCD) camera (Princeton instruments). The samples were loaded into the X-ray microscope, and collection of a VLM mosaic using a 20x light objective followed. ROIs were identified based on the cryo-SIM data, and 2D X-ray Mosaic Maps were generated for the selected ROIs. Tilted-series collection followed, in a range of -60° to +60°, using at increments of 0.2° or 0.5°, using the XRM Data Explorer software.

### Cryo-SXT Data Reconstruction on IMOD

Using the Cygwin unix-like environment, .tiff data were transformed into .st files, and cryo-SXT data reconstruction followed as per manufacturer’s protocol, using Etomo, IMOD (University of Colorado)(80). Fiducial models were generated manually, with fine alignment mean residual error being reduced gradually, reaching a value below ≤ 0.2 nm where possible. Tomogram generation was performed using the back-projection method, and in cases simultaneous iterative reconstruction technique (SIRT) tomogram generation was additionally performed, in a computational system supporting CUDA (NVIDIA) (B24, DLS).

### Correlation of Cryo-SIM and Cryo-SXT Data on EC-CLEM, Icy

Correlation between reconstructed cryo-SIM and their respective cryo-SXT was performed as previously described (79) using EC-CLEM, Icy(81). In brief, the chromatic shift-corrected cryo-SIM data, were corrected for stage shifts by 3D registrations, using intermediate channels between collections. Once the data were corrected for both chromatic and stage shifts, sequential 2D data registrations corresponding to the cryo-SIM data to the cryo-SXT location were performed, and after the 3D cryo-SIM data were transformed to fit the 3D cryo-SXT location, 3D registration between them was performed, correlating data on the z-plane.

### Lentiviral transduction

CD8^+^ T-cells were stimulated for 2 days with anti-CD3 and anti-CD28 mAb microbeads prior to transduction with Lentivirus. Day of transduction, cells are harvested and counted before being reseeded at 1x10^6^ cells per ml of X-Vivo complete to which titrated lentivirus with 300IU IL-2 are added to each well. Each well is then then gently pipetted to thoroughly mix. Cells are counted and split 48-72hrs post virus addition and seeded at 0.2x10^6^ cells per ml X-Vivo complete. D5 cells are harvested and counted with 0.1x10^6^ being stained with an anti-EGFR labeled fluorochrome for transduction purity. Remaining cells are labeled using anti-EGFR PE (BioLegend) and an anti-PE kit (Miltenyi Biotec) for selection of transduced cells using magnetic column enrichment. A portion of enriched cells are then checked for purity using flow-based cytometry. Enriched cells are then either frozen using CryoStor CS10 (Millipore Sigma) or directly added to Treg induction conditions.

### siRNA treatment

CD8-iTreg and CTLs were transfected by electroporation as described (82, 83). Briefly, Treg were transfected with 10µM of either granzyme K siRNA or non-targeting Control siRNA-A (Santa Cruz). Electroporation was performed using the P3 primary cell 4D-nucleofector kit (Lonza) and a Nucleofector-4D machine (Lonza) on setting EO-115. After electroporation, transfected cells were transferred to 6-well tissue-culture treated plate containing 2ml of pre-warmed complete-RPMI with 300IU/ml of rH-IL-2. Cells were rested at 37°C/5%CO_2_ for 2hrs. After resting, wells were supplemented with 8ml complete-RPMIc with 300U/ml IL-2 and 10ng/ml TGFβ. Assessment of protein knockdown and function were completed 48-72hr after electroporation.

### CRISPR/Cas9a RNP

The ablation of genes of interests was achieved by electroporation with pre-assembled CRISPR/Cas9 RNP complexes. Liquid nitrogen stored CD8-iTreg were thawed and recovered overnight (1M/ml) in presence of rhIL-2 (300U/ml), washed three times with 10 volumes of pre-warmed Opti-MEM (ThermoFisher Scientific). Cells were resuspended to a final 1 × 10^6^ cells in 50 µL of Opti-MEM suspension. In parallel, RNP complexes were assembled *in vitro* in two steps. First, 150 pmol of Alt-R CRISPR-Cas9 tracrRNA (200 µM stock; Integrated DNA Technologies (IDT)) were mixed with 150 pmol of Alt-R CRISPR-Cas9 predesigned crRNA (200 µM stock) and then incubated at 95 °C for 5 min and the resultant duplex guide RNA allowed to cool to room temperature. All Alt-R CRISPR-Cas9 crRNA sequences and cat# are given in Supplementary Table (all on- and off-target scores were optimized by IDT). The duplex gRNA was then mixed with 150 pmol of Alt-R *S. pyogenes* CRISPR-Cas9 Nuclease V3 (IDT, 20 µM stock) and incubated at 37 °C for 15 min. The preformed RNPs were allowed to cool to room temperature and then supplemented with 150 pmol of Alt-R Cas9 Electroporation Enhancer (IDT, 200 µM stock). Cells were then mixed with the RNP solution and immediately transferred to a 2-mm cuvette (Bio-Rad) and electroporated at 300 V for 2 ms using an ECM830 Square Wave electroporator. Immediately after transfection, cells were recovered with prewarmed, IL-2 supplemented RPMI 1640 media, recovered and hit again 24hrs later, followed by another 24hr recovery prior to the experiments.

### Supported lipid bilayer (SLB)

Preparation of liposomes and mobile SLB formation were described in detail elsewhere^37^. In brief, SLB were formed by incubation with mixtures of small unilamellar vesicles to generate a final lipid composition of 12.5 mol% 1,2-dioleoyl-sn-glycero-3-[(N-(5-amino-1 carboxypentyl) iminodiacetic acid) succinyl] (DOGS-NTA) and 87.5 mol% 1,2-dioleoyl-sn-glycero-3-phosphocholine supplemented (DOPC) to yield 30 molecules/μm2 anti-CD3e(UCHT1)-Fab-6his and 200 molecules/µm2 mouse ICAM-1-12his at a total lipid concentration of 0.4 mM. Lipid droplets were deposited onto clean glass coverslips (SCHOTT; #1472315) of the flow chamber (sticky-Slide VI 0.4, Ibidi; #80608). After 20 min incubation, the flow chamber was flooded with HEPES Buffered Saline 1X pH 7.2 (0.2 mM HEPES, 1.37 mM NaCl, 50 nM KCl, 7 nM Na2HPO4, 60 nM D-glucose, 1 mM CaCl2, 2 mM MgCl2) supplemented with 0.1 % Human Serum Albumin (HSA) (Merck Millipore; #12667-50ml) and flushed to remove excess liposomes. After blocking for 20 minutes with 5% BSA in HEPES Buffered Saline containing 100 mM NiSO4 to saturate NTA sites, unbound proteins were flushed out by HSA/HBS, and the SLB was ready to be used. SLB were uniformly fluid as determined by fluorescence recovery after photobleaching. Protein concentrations required to achieve desired densities on bilayers were calculated from calibration curves constructed from flow cytometric measurements of bilayer-associated fluorescence of attached proteins on bilayers formed on glass beads, compared with reference beads containing known numbers of the appropriate fluorophore (Bangs Laboratories; #647-A). All lipids were purchased from Avanti Polar Lipids, Inc.

### Total internal reflection fluorescence (TIRF) imaging

For immunological synapse formation on activating surfaces, T-cells were plated onto activating ICAM1 + anti-CD3e Fab’ surfaces (SLB, see protocol above) for 15 min, washed then fixed with 3.7% paraformaldehyde (PFA) in PBS 1X (v/v) for 15 min at 37C, followed by a PBS wash. For SMAP analysis, T-cells were plated onto same surfaces for 90 minutes at 37C, followed by flushing T-cells off by pushing PBS through the IBIDI channels, followed by fixation with 3.7% PFA and 0.25% glutaraldehyde for 15 minutes at 37C, and a final PBS wash. After fixation, samples that had been activated on SLBs were washed twice with 0.1% BSA/HBS, blocked with 5% BSA/HBS, and treated with Image-IT FX Signal Enhancer for 30 min at RT. Followed by staining with 5µg/ml antibodies (anti-TCRab (IP26; BioLegend); Granzyme B (GB11; BioLegend); Perforin DB-48 (BioLegend); Granzyme K (GM26E7; BioLegend); Granzyme A (CB9; BioLegend); or 10µg/ml thrombospondin-1 (A6.1; Thermo Fisher); thrombospondin-4 (PA5-68467; Thermo Fisher) overnight incubation at 4C, or Wheat Germ Agglutinin directly labelled from Thermo Fisher and stained for 10 minutes at RT. TIRF microscopy was completed using a Nikon ECLIPSE Ti2-E microscope equipped with a Yokogawa CSU-W1 SoRA spinning disk confocal unit and a Photometrics Prime BSI (Nikon). A 60X/1.49 NA oil immersion objective was used for image acquisition.

### Direct Stochastic Optical Reconstruction Microscopy (dSTORM) imaging

The For dSTORM, CD8-iTreg were incubated on activating pSLBs coated with mICAM-1-his and anti-CD3ε (UCHT-1)-Fab-his for 90 minutes, followed by a PBS wash to flush of cells and leave released particles behind on the pSLB. These particles were chemically fixed with 3.7% paraformaldehyde and 0.25% glutaraldehyde to preserve their ultrastructure, followed by staining for putative SMAP components. Oxford NanoImager (ONI) microscope was used to generate 2D SMLM images using dSTORM. A 100X objective lens/1.4 NA in oil-immersion was used. 405 nm, 488 nm, and 640 nm lasers were used with 2 channels (with a dichroic mirror split at 640 nm). All calibrations, controls, and images were performed at 27°C with a supercritical TIRF angle of 54°. Samples were imaged in dSTORM buffer (320 µl 1x PBS, 40 µl 50% glucose, 40 µl cysteamine (MEA), 4 µl glucose-oxidase). First, the 640 nm laser was used at 40% laser power to excite UCHT1-AF647, then the 488 nm laser at 60% to excite AF488. The 405 nm laser was used to promote fluorophore blinking. 3,000 frames were acquired per fluorophore. An exposure of 30ms was used. Calibration channel mapping performed using TetraSpec Microspheres (Thermo) of 100 nm were used to align all channels and generate the final dSTORM image using the Oxford NanoImager (ONI) analysis platform, CODI.

### Direct Stochastic Optical Reconstruction Microscopy (dSTORM) image analysis

Localization, acquisition, and mapping files were uploaded for each image taken. Drift correction was applied using the DME algorithm. Frames were filtered first according to acquisition program settings, then to exclude the initial excitation peaks, the first 100 frames are removed for each laser. The steady state of localizations per frame was included as true blinking localizations. Filtering parameters around the sigma peak were then applied to include a sigma range of approximately 50-200 nm (Gaussian fit to localize single molecules). Localization precision filtering was then applied to include localization with precision less than or equal to 20 nm. Micrographs of T-cell immunological synapses were depicted with a display sigma of 5 nm using fixed representation. Files were downloaded from CODI in TIFF and PNG formats, then cropped and converted to PNG micrographs for figures.

### Airyscan microscop

Airyscan imaging of pSLBs was performed on a confocal laser-scanning microscope Zeiss LSM 880 equipped with Airyscan detection module (Zeiss, Oberkochen, Germany) using the Plan-Apochromat 63×/1.46 Oil objective (Zeiss, Oberkochen, Germany). The emission signals were collected on the 32 channel GaAsP-PMT Airy detector. ZEN Airyscan software (Zeiss) was used to process the acquired data sets. This software processes each of the 32 Airy detector channels separately by performing filtering, deconvolution and pixel reassignment in order to obtain images with enhanced resolution and improved signal to noise ratio. The value of Wiener filter in ZEN software was chosen in accordance with the value in ‘auto’ reconstruction modality and was set around 7, to ensure the absence of deconvolution artefacts.

### Cell lysates and immunoblots

Cells were washed once with ice-cold 1X PBS and lysed in buffer containing 20 mM Tris-HCl (pH 8), 150 mM NaCl and 1% Triton X-100, supplemented with Protease Inhibitor Cocktail Set III (Calbiochem) and 0.2 mg sodium orthovanadate. Homogenates were incubated on ice for 5 minutes, then centrifuged at 16,200 x g for 20 minutes at 4°C, and the soluble fractions were collected. Protein concentration in the post-nuclear supernatants was determined using the Quantum Protein Bicinchoninic Acid (BCA) Assay (Euroclone), with absorbance measured at 570 nm using an iMark™ microplate reader (Bio-Rad Laboratories). 10 µg of protein extract were denatured in Bolt™ Sample Reducing Agent supplemented with Bolt™ LDS Sample Buffer (Invitrogen™) for 5 minutes at 100°C. Samples were resolved on precast 4-12% Bis-Tris polyacrylamide gradient gels (1.0 mm) in 1X Bolt™ MES SDS Running Buffer at a constant voltage of 150 V. Separated proteins were transferred onto nitrocellulose membranes (Amersham Protran^®^) in a transfer buffer containing 20 mM Tris, 0.2 M glycine, and 20% (v/v) ethanol for 75 minutes at a constant current of 250 mA. Membranes were blocked in 4% (w/v) non-fat dry milk in 1× PBS (140 mM NaCl, 2.7 mM KCl, 10 mM phosphate buffer, pH 7.4) supplemented with 0.02% (v/v) Tween-20 (PBS-T), then incubated with anti-Thrombospondin-4 (F7; Santa Cruz Biotechnology) and anti-actin (C4; EMD Millipore) primary antibodies and horseradish peroxidase-conjugated goat anti-mouse IgG secondary antibodies (Jackson ImmunoResearch Inc). Immunoreactive bands were detected using a chemiluminescent substrate and signals were acquired using the Alliance Q9-ATOM imaging system with NineAlliance x64 software (UVITEC).

## REFERENCES

1. Bolaños-Meade J, Hamadani M, Wu J, Al Malki MM, Martens MJ, Runaas L, et al. Post-transplantation cyclophosphamide-based graft-versus-host disease prophylaxis. New England Journal of Medicine. 2023;388(25):2338–48.

2. McCurdy SR, Luznik L. Relapse after allogeneic transplantation with post-transplant cyclophosphamide: Shattering myths and evolving insight. Blood Reviews. 2023:101093.

3. Bolanos-Meade J, Reshef R, Fraser R, Fei M, Abhyankar S, Al-Kadhimi Z, et al. Three prophylaxis regimens (tacrolimus, mycophenolate mofetil, and cyclophosphamide; tacrolimus, methotrexate, and bortezomib; or tacrolimus, methotrexate, and maraviroc) versus tacrolimus and methotrexate for prevention of graft-versus-host disease with haemopoietic cell transplantation with reduced-intensity conditioning: a randomised phase 2 trial with a non-randomised contemporaneous control group (BMT CTN 1203). Lancet Haematol. 2019;6(3):e132–e43.

4. McDonald-Hyman C, Turka LA, Blazar BR. Advances and challenges in immunotherapy for solid organ and hematopoietic stem cell transplantation. Science translational medicine. 2015;7(280):280rv2–rv2.

5. Meyer EH, Salhotra A, Gandhi AP, Pantin J, Patel SS, Hoeg RT, et al. Orca-T vs allogeneic hematopoietic stem cell transplantation (Precision-T): a multicenter, randomized phase 3 trial. Blood. 2026;147(11):1168–77.

6. Siegel RL, Miller KD, Wagle NS, Jemal A. Cancer statistics, 2023. CA Cancer J Clin. 2023;73(1):17–48.

7. Lennmyr E, Karlsson K, Ahlberg L, Garelius H, Hulegardh E, Izarra AS, et al. Survival in adult acute lymphoblastic leukaemia (ALL): A report from the Swedish ALL Registry. Eur J Haematol. 2019;103(2):88–98.

8. Frigault MJ, Maus MV. State of the art in CAR T cell therapy for CD19+ B cell malignancies. The Journal of Clinical Investigation. 2020;130(4):1586–94.

9. Majzner RG, Mackall CL. Clinical lessons learned from the first leg of the CAR T cell journey. Nature medicine. 2019;25(9):1341–55.

10. Martino M, Alati C, Canale FA, Musuraca G, Martinelli G, Cerchione C. A Review of Clinical Outcomes of CAR T-Cell Therapies for B-Acute Lymphoblastic Leukemia. Int J Mol Sci. 2021;22(4).

11. Schmidts A, Wehrli M, Maus MV. Toward Better Understanding and Management of CAR-T Cell–Associated Toxicity. Annual Review of Medicine. 2021;72:365–82.

12. Siegler EL, Kenderian SS. Neurotoxicity and cytokine release syndrome after chimeric antigen receptor T cell therapy: insights into mechanisms and novel therapies. Frontiers in immunology. 2020;11:1973.

13. Brudno JN, Kochenderfer JN. Toxicities of chimeric antigen receptor T cells: recognition and management. Blood, The Journal of the American Society of Hematology. 2016;127(26):3321–30.

14. Hay KA, Hanafi LA, Li D, Gust J, Liles WC, Wurfel MM, et al. Kinetics and biomarkers of severe cytokine release syndrome after CD19 chimeric antigen receptor-modified T-cell therapy. Blood. 2017;130(21):2295–306.

15. Bader CS, Meyer EH, Negrin RS. Regulatory T cell approaches for graft-versus-host disease prevention. Curr Opin Immunol. 2026;98:102685.

16. Bolivar-Wagers S, Loschi ML, Jin S, Thangavelu G, Larson JH, McDonald-Hyman CS, et al. Murine CAR19 Tregs suppress acute graft-versus-host disease and maintain graft-versus-tumor responses. JCI insight. 2022;7(17).

17. Zhao DM, Thornton AM, DiPaolo RJ, Shevach EM. Activated CD4+CD25+ T cells selectively kill B lymphocytes. Blood. 2006;107(10):3925–32.

18. Gondek DC, Lu LF, Quezada SA, Sakaguchi S, Noelle RJ. Cutting edge: Contact-mediated suppression by CD4+CD25+ regulatory cells involves a granzyme B-dependent, perforin-independent mechanism. J Immunol. 2005;174(4):1783–6.

19. Gondek DC, Devries V, Nowak EC, Lu LF, Bennett KA, Scott ZA, et al. Transplantation survival is maintained by granzyme B+ regulatory cells and adaptive regulatory T cells. J Immunol. 2008;181(7):4752–60.

20. Bolivar-Wagers S, Larson JH, Jin S, Blazar BR. Cytolytic CD4+ and CD8+ regulatory T-cells and implications for developing immunotherapies to combat graft-versus-host disease. Frontiers in immunology. 2022;13.

21. Boroughs AC, Larson RC, Choi BD, Bouffard AA, Riley LS, Schiferle E, et al. Chimeric antigen receptor costimulation domains modulate human regulatory T cell function. JCI Insight. 2019;5(8).

22. Heinrichs J, Li J, Nguyen H, Wu Y, Bastian D, Daethanasanmak A, et al. CD8(+) Tregs promote GVHD prevention and overcome the impaired GVL effect mediated by CD4(+) Tregs in mice. Oncoimmunology. 2016;5(6):e1146842.

23. Suzuki M, Jagger AL, Konya C, Shimojima Y, Pryshchep S, Goronzy JJ, et al. CD8+ CD45RA+ CCR7+ FOXP3+ T cells with immunosuppressive properties: a novel subset of inducible human regulatory T cells. The Journal of Immunology. 2012;189(5):2118–30.

24. Ligocki AJ, Niederkorn JY. Advances on Non-CD4+ Foxp3+ T regulatory cells: CD8+, Tr1, and double negative T regulatory cells in organ transplantation. Transplantation. 2015;99(8):1553.

25. Schmidt A, Eriksson M, Shang M-M, Weyd H, Tegnér J. Comparative analysis of protocols to induce human CD4+ Foxp3+ regulatory T cells by combinations of IL-2, TGF-beta, retinoic acid, rapamycin and butyrate. PloS one. 2016;11(2):e0148474.

26. Zheng J, Liu Y, Liu Y, Liu M, Xiang Z, Lam K-T, et al. Human CD8+ regulatory T cells inhibit GVHD and preserve general immunity in humanized mice. Science translational medicine. 2013;5(168):168ra9–ra9.

27. Schmitt EG, Williams CB. Generation and function of induced regulatory T cells. Frontiers in immunology. 2013;4:152.

28. Bilate AM, Lafaille JJ. Induced CD4+ Foxp3+ regulatory T cells in immune tolerance. Annual review of immunology. 2012;30:733–58.

29. Vieyra-Lobato MR, Vela-Ojeda J, Montiel-Cervantes L, López-Santiago R, Moreno-Lafont MC. Description of CD8+ regulatory T lymphocytes and their specific intervention in graft-versus-host and infectious diseases, autoimmunity, and cancer. Journal of immunology research. 2018;2018.

30. Koch SD, Uss E, van Lier RA, ten Berge IJ. Alloantigen-induced regulatory CD8+ CD103+ T cells. Human immunology. 2008;69(11):737–44.

31. Cosmi L, Liotta F, Lazzeri E, Francalanci M, Angeli R, Mazzinghi B, et al. Human CD8+CD25+ thymocytes share phenotypic and functional features with CD4+CD25+ regulatory thymocytes. Blood. 2003;102(12):4107–14.

32. Grakoui A, Bromley SK, Sumen C, Davis MM, Shaw AS, Allen PM, et al. The immunological synapse: a molecular machine controlling T cell activation. Science. 1999;285(5425):221–7.

33. Shi L, Kam CM, Powers JC, Aebersold R, Greenberg AH. Purification of three cytotoxic lymphocyte granule serine proteases that induce apoptosis through distinct substrate and target cell interactions. J Exp Med. 1992;176(6):1521–9.

34. Balint S, Muller S, Fischer R, Kessler BM, Harkiolaki M, Valitutti S, et al. Supramolecular attack particles are autonomous killing entities released from cytotoxic T cells. Science. 2020;368(6493):897–901.

35. Cassioli C, Capitani N, Staton CC, Schirra C, Finetti F, Onnis A, et al. Activation-induced thrombospondin-4 works with thrombospondin-1 to build cytotoxic supramolecular attack particles. Proceedings of the National Academy of Sciences. 2025;122(6):e2413866122.

36. Hippen K, Merkel S, Schirm D, Nelson C, Tennis N, Riley J, et al. Generation and Large-Scale Expansion of Human Inducible Regulatory T Cells That Suppress Graft-Versus-Host Disease. American Journal of Transplantation. 2011;11(6):1148–57.

37. Brunstein CG, Miller JS, McKenna DH, Hippen KL, DeFor TE, Sumstad D, et al. Umbilical cord blood–derived T regulatory cells to prevent GVHD: kinetics, toxicity profile, and clinical effect. Blood, The Journal of the American Society of Hematology. 2016;127(8):1044–51.

38. Hori S, Nomura T, Sakaguchi S. Control of regulatory T cell development by the transcription factor Foxp3. Science. 2003;299(5609):1057–61.

39. Joeris T, Gomez-Casado C, Holmkvist P, Tavernier SJ, Silva-Sanchez A, Klotz L, et al. Intestinal cDC1 drive cross-tolerance to epithelial-derived antigen via induction of FoxP3(+)CD8(+) T(regs). Sci Immunol. 2021;6(60).

40. Hippen KL, Harker-Murray P, Porter SB, Merkel SC, Londer A, Taylor DK, et al. Umbilical cord blood regulatory T-cell expansion and functional effects of tumor necrosis factor receptor family members OX40 and 4-1BB expressed on artificial antigen-presenting cells. Blood. 2008;112(7):2847–57.

41. Hoffmann P, Hofmeister R, Brischwein K, Brandl C, Crommer S, Bargou R, et al. Serial killing of tumor cells by cytotoxic T cells redirected with a CD19-/CD3-bispecific single-chain antibody construct. Int J Cancer. 2005;115(1):98–104.

42. Hippen KL, Merkel SC, Schirm DK, Sieben CM, Sumstad D, Kadidlo DM, et al. Massive ex vivo expansion of human natural regulatory T cells (Tregs) with minimal loss of in vivo functional activity. Science translational medicine. 2011;3(83):83ra41–83ra41.

43. Li J, Zaslavsky M, Su Y, Guo J, Sikora MJ, van Unen V, et al. KIR+ CD8+ T cells suppress pathogenic T cells and are active in autoimmune diseases and COVID-19. Science. 2022:eabi9591.

44. Li J, Zaslavsky M, Su Y, Sikora MJ, van Unen V, Christophersen A, et al. Human KIR+ CD8+ T cells target pathogenic T cells in Celiac disease and are active in autoimmune diseases and COVID-19. bioRxiv. 2021:2021.12. 23.473930.

45. Gao Y, Liu R, Shi J, Shan W, Zhou H, Chen Z, et al. Clonal GZMK+ CD8+ T cells are identified as a hallmark of the pathogenesis of cGVHD-induced bronchiolitis obliterans syndrome after allogeneic hematopoietic stem cell transplantation. EBioMedicine. 2025;112.

46. Harkiolaki M, Darrow MC, Spink MC, Kosior E, Dent K, Duke E. Cryo-soft X-ray tomography: using soft X-rays to explore the ultrastructure of whole cells. Emerging topics in life sciences. 2018;2(1):81–92.

47. Jacobsen C. Soft x-ray microscopy. Trends in cell biology. 1999;9(2):44–7.

48. Chang HF, Schirra C, Ninov M, Hahn U, Ravichandran K, Krause E, et al. Identification of distinct cytotoxic granules as the origin of supramolecular attack particles in T lymphocytes. Nat Commun. 2022;13(1):1029.

49. Zeiser R, Blazar BR. Acute Graft-versus-Host Disease - Biologic Process, Prevention, and Therapy. N Engl J Med. 2017;377(22):2167–79.

50. Shah BD, Bishop MR, Oluwole OO, Logan A, Baer MR, Donnellan WB, et al. End of phase I results of ZUMA-3, a phase 1/2 study of KTE-X19, anti-CD19 chimeric antigen receptor (CAR) T cell therapy, in adult patients (pts) with relapsed/refractory (R/R) acute lymphoblastic leukemia (ALL). American Society of Clinical Oncology; 2019.

51. Siegel RL, Miller KD, Jemal A. Cancer statistics, 2019. CA: a cancer journal for clinicians. 2019;69(1):7–34.

52. Allen CE, Marsh R, Dawson P, Bollard CM, Shenoy S, Roehrs P, et al. Reduced-intensity conditioning for hematopoietic cell transplant for HLH and primary immune deficiencies. Blood. 2018;132(13):1438–51.

53. Steinbach A, Clark SM, Clemmons AB. Bosutinib: a novel src/abl kinase inhibitor for chronic myelogenous leukemia. J Adv Pract Oncol. 2013;4(6):451–5.

54. Philipson BI, O’Connor RS, May MJ, June CH, Albelda SM, Milone MC. 4-1BB costimulation promotes CAR T cell survival through noncanonical NF-κB signaling. Sci Signal. 2020;13(625).

55. Suhoski MM, Golovina TN, Aqui NA, Tai VC, Varela-Rohena A, Milone MC, et al. Engineering artificial antigen-presenting cells to express a diverse array of co-stimulatory molecules. Molecular Therapy. 2007;15(5):981–8.

56. Hippen KL, Bucher C, Schirm DK, Bearl AM, Brender T, Mink KA, et al. Blocking IL-21 signaling ameliorates xenogeneic GVHD induced by human lymphocytes. Blood. 2012;119(2):619–28.

57. Parmar S, Liu X, Tung SS, Robinson SN, Rodriguez G, Cooper LJ, et al. Third-party umbilical cord blood-derived regulatory T cells prevent xenogenic graft-versus-host disease. Cytotherapy. 2014;16(1):90–100.

58. Ali N, Flutter B, Sanchez Rodriguez R, Sharif-Paghaleh E, Barber LD, Lombardi G, et al. Xenogeneic Graft-versus-Host-Disease in NOD-scid IL-2RγNull Mice Display a T-Effector Memory Phenotype. PLoS ONE. 2012;7(8):219–29.

59. Lu Y, Hippen KL, Lemire AL, Gu J, Wang W, Ni X, et al. miR-146b antagomir–treated human Tregs acquire increased GVHD inhibitory potency. Blood. 2016;128(10):1424–35.

60. Stoeckius M, Hafemeister C, Stephenson W, Houck-Loomis B, Chattopadhyay PK, Swerdlow H, et al. Simultaneous epitope and transcriptome measurement in single cells.

61. Frankish A, Carbonell-Sala S, Diekhans M, Jungreis I, Loveland JE, Mudge JM, et al. GENCODE: reference annotation for the human and mouse genomes in 2023. Nucleic acids research. 2023;51(D1):D942–D9.

62. Zheng GXY, Terry JM, Belgrader P, Ryvkin P, Bent ZW, Wilson R, et al. Massively parallel digital transcriptional profiling of single cells.

63. Roelli P, Flynn B, Gui G. Hoohm/CITE-seq-Count: 1.4. 2. zenodo. 2019.

64. Xin H, Lian Q, Jiang Y, Luo J, Wang X, Erb C, et al. GMM-Demux: sample demultiplexing, multiplet detection, experiment planning, and novel cell-type verification in single cell sequencing. Genome Biol. 2020;21(1):188.

65. McGinnis CS, Patterson DM, Winkler J, Conrad DN, Hein MY, Srivastava V, et al. MULTI-seq: sample multiplexing for single-cell RNA sequencing using lipid-tagged indices. Nat Methods. 2019;16(7):619–26.

66. Boggy GJ, McElfresh GW, Mahyari E, Ventura AB, Hansen SG, Picker LJ, et al. BFF and cellhashR: analysis tools for accurate demultiplexing of cell hashing data. Bioinformatics. 2022;38(10):2791–801.

67. Hao Y, Stuart T, Kowalski MH, Choudhary S, Hoffman P, Hartman A, et al. Dictionary learning for integrative, multimodal and scalable single-cell analysis.

68. Love MI, Huber W, Anders S. Moderated estimation of fold change and dispersion for RNA-seq data with DESeq2. Genome Biol. 2014;15(12):550.

69. Consortium TGO. The Gene Ontology knowledgebase in 2026. Nucleic Acids Research. 2025;54(D1):D1779–D92.

70. Kanehisa M, Goto S. KEGG: kyoto encyclopedia of genes and genomes. Nucleic Acids Res. 2000;28(1):27–30.

71. Liberzon A, Birger C, Thorvaldsdóttir H, Ghandi M, Mesirov JP, Tamayo P. The Molecular Signatures Database (MSigDB) hallmark gene set collection. Cell Syst. 2015;1(6):417–25.

72. Yu G, Wang LG, Han Y, He QY. clusterProfiler: an R package for comparing biological themes among gene clusters. Omics. 2012;16(5):284–7.

73. La Manno G, Soldatov R, Zeisel A, Braun E, Hochgerner H, Petukhov V, et al. RNA velocity of single cells. Nature. 2018;560(7719):494–8.

74. Bergen V, Lange M, Peidli S, Wolf FA, Theis FJ. Generalizing RNA velocity to transient cell states through dynamical modeling. Nature biotechnology. 2020;38(12):1408–14.

75. Lange M, Bergen V, Klein M, Setty M, Reuter B, Bakhti M, et al. CellRank for directed single-cell fate mapping. Nature methods. 2022;19(2):159–70.

76. Setty M, Kiseliovas V, Levine J, Gayoso A, Mazutis L, Pe’Er D. Characterization of cell fate probabilities in single-cell data with Palantir. Nature biotechnology. 2019;37(4):451–60.

77. Okolo CA, Fish TM, Nahas KL, Jadhav AC, Vyas N, Taylor A, et al., editors. A combination of soft X-ray and laser light sources offer 3D high content information on the native state of the cellular environment. Journal of Physics: Conference Series; 2022: IOP Publishing.

78. Phillips MA, Harkiolaki M, Susano Pinto DM, Parton RM, Palanca A, Garcia-Moreno M, et al. CryoSIM: super-resolution 3D structured illumination cryogenic fluorescence microscopy for correlated ultrastructural imaging. Optica. 2020;7(7):802–12.

79. Vyas N, Kunne S, Fish TM, Dobbie IM, Harkiolaki M, Paul-Gilloteaux P. Protocol for image registration of correlative soft X-ray tomography and super-resolution structured illumination microscopy images. STAR protocols. 2021;2(2).

80. Kremer JR, Mastronarde DN, McIntosh JR. Computer visualization of three-dimensional image data using IMOD. Journal of structural biology. 1996;116(1):71–6.

81. Paul-Gilloteaux P, Heiligenstein X, Belle M, Domart M-C, Larijani B, Collinson L, et al. eC-CLEM: flexible multidimensional registration software for correlative microscopies. Nature methods. 2017;14(2):102–3.

82. Zanin-Zhorov A, Ding Y, Kumari S, Attur M, Hippen KL, Brown M, et al. Protein Kinase C-Theta Mediates Negative Feedback on Regulatory T Cell Function. Science. 2010;328(5976):372–6.

83. McDonald-Hyman C, Muller JT, Loschi M, Thangavelu G, Saha A, Kumari S, et al. The vimentin intermediate filament network restrains regulatory T cell suppression of graft-versus-host disease. J Clin Invest. 2018;128(10):4604–21.

