## Supplemental Results and Methods for "Human CD8-iTreg are potent GVHD suppressors and tumoricidal effectors by release of Granzyme-K^+^ Supramolecular Attack Particles"

**SUPPLEMENTAL FIGURES**

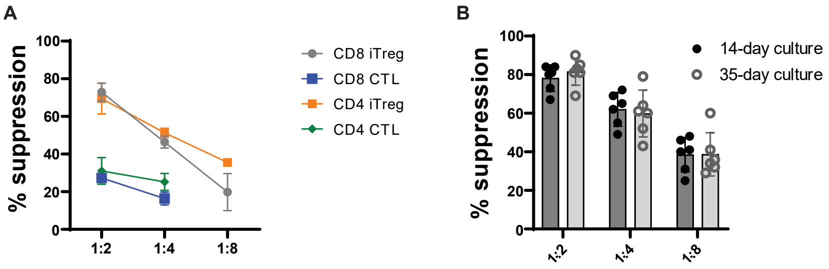


***Supplemental Figure 1. (A)*** *In vitro suppression of activated PBMC proliferation by donor-matched CD8-iTreg vs CD8-CTLs vs CD4$^+$ iTreg vs CD4$^+$ CTLs. CD8 and CD4 iTreg and CTLs were generated from donor-matched PB CD8$^+$CD25$^-$ or CD4+CD25$^-$ T-cells, respectively, stimulated for 14-days with either IL-2 alone or IL-2, Rapamycin, and TGFβ.* ***(B)*** *In vitro suppression of activated PBMC proliferation by 14-day vs 35-day cultured CD8-iTreg at different effector-to-PBMC ratios (1:2, 1:4, 1:8) vs PBMC-only controls (0:1).*

***Supplemental Table 1. Comprehensive 32-Parameter Spectral Flow Cytometry Panels.*** *List of markers analyzed across two experimental panels evaluating T-cell activation, cytotoxicity, immunosuppression, and canonical CD4*$^{+}$ *and CD8*$^{+}$ *nTreg phenotypes.*

| PANEL 1 | MARKER | CLONE | PANEL 2 | MARKER | CLONE |
|---|---|---|---|---|---|
| | **CD103** | Ber-ACT8 | | **GzmB** | GzA-3G8.5 |
| | **CCR7** | G043H7 | | **CCR7** | G043H7 |
| | **CD127** | HIL 7R M21 | | **CD103** | Ber-ACT8 |
| | **CD158e,k** | KIR3DL1/DL2 | | **CD27** | L128 |
| | **CD25** | M-A251 | | **CD28** | M-A251 |
| | **CD27** | L128 | | **CD3** | OKT3 |
| | **CD28** | M-A251 | | **CD4** | RPA-T4 |
| | **CD39** | TU66 | | **CD49α** | TS2/7 |
| | **CD4** | RPA-T4 | | **CD49β** | P1E6-C5 |
| | **CD8** | SK1 | | **CD57** | HNK-1 |
| | **CD95 (FAS)** | DX2 | | **CD8** | SK1 |
| | **Foxp3** | 259d | | **CD95 (FAS)** | DX2 |
| | **GAPR** | 7B11 | | **CTLA-4** | BNI3 |
| | **GITR** | 621 | | **CX3CR1** | 2A9-1 |
| | **GNLY** | DH2 | | **GzmM** | GM26E7 |
| | **GzmB** | GzA-3G8.5 | | **GznK** | NGZB |
| | **GzmK** | NGZB | | **KLRG1** | SA231A2 |
| | **GzmM** | GM26E7 | | **KLRG1** | SA231A2 |
| | **HLA-E** | 4B2G4 | | **Perforin** | dG9 |
| | **Lag-3** | T47-530 | | | |
| | **PD-1** | J105 | | | |
| | **TIGIT** | A15153G | | | |

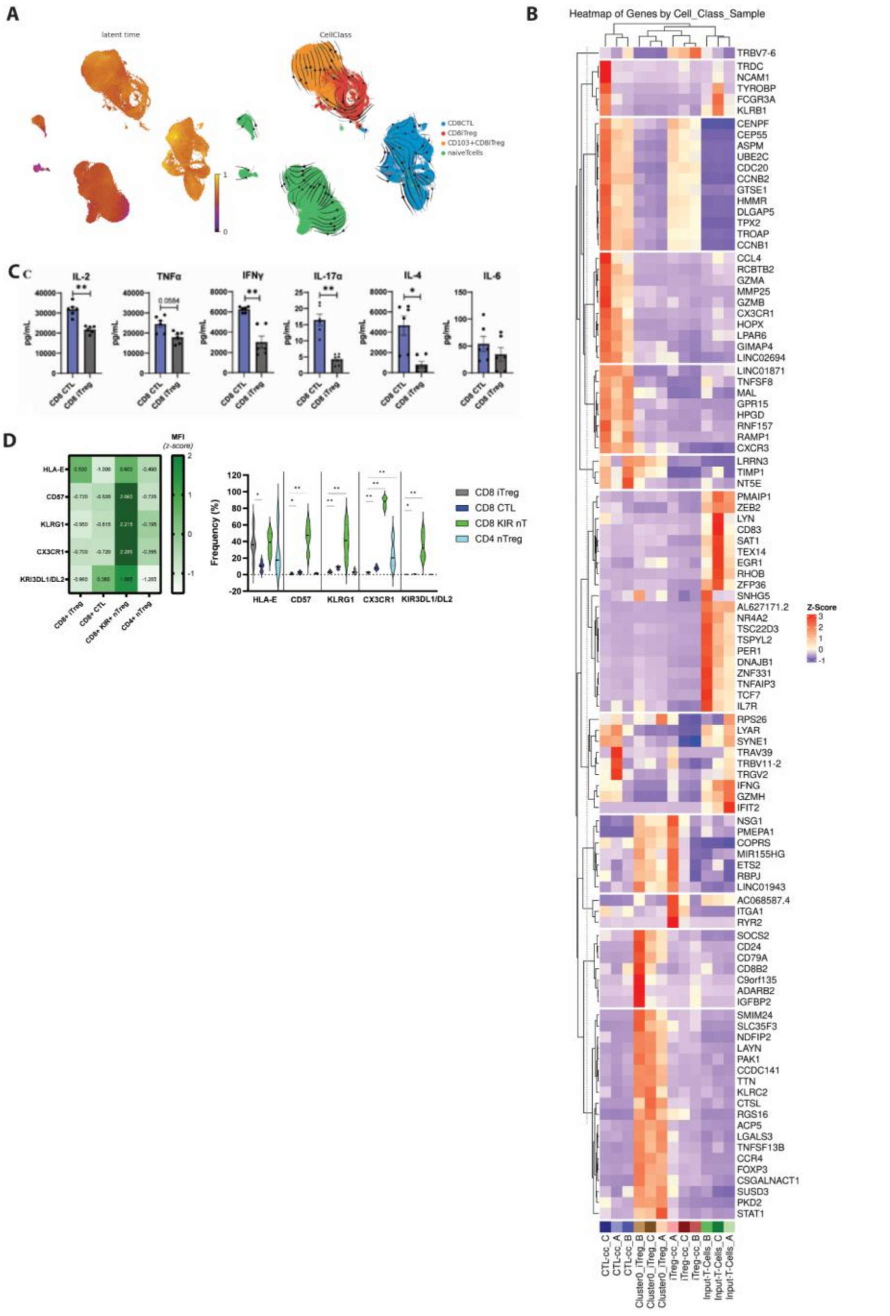
A
latent time
CellClass
CD8CTL
CD8iTreg
CD103+CD8iTreg
naiveTcells
B
Heatmap of Genes by Cell_Class_Sample
Z-Score
C
IL-2
TNFα
IFNγ
IL-17α
IL-4
IL-6
pg/mL
0.0584
CD8 CTL
CD8 iTreg
D
MFI (z-score)
HLA-E
CD57
KLRG1
CX3CR1
KIR3DL1/DL2
CD8+ iTreg
CD8+ CTL
CD8+ KIR+ nTreg
CD4+ nTreg
Frequency (%)
CD8 iTreg
CD8 CTL
CD8 KIR nT
CD4 nTreg
TRBV7-6
TRDC
NCAM1
TYROBP
FCGR3A
KLRB1
CENPF
CEP55
ASPM
UBE2C
CDC20
CCNB2
GTSE1
HMMR
DLGAP5
TPX2
TROAP
CCNB1
CCL4
RCBTB2
GZMA
MMP25
GZMB
CX3CR1
HOPX
LPAR6
GIMAP4
LINC02694
LINC01871
TNFSF8
MAL
GPR15
HPGD
RNF157
RAMP1
CXCR3
LRRN3
TIMP1
NT5E
PMAIP1
ZEB2
LYN
CD83
SAT1
TEX14
EGR1
RHOB
ZFP36
SNHG5
AL627171.2
NR4A2
TSC22D3
TSPYL2
PER1
DNAJB1
ZNF331
TNFAIP3
TCF7
IL7R
RPS26
LYAR
SYNE1
TRAV39
TRBV11-2
TRGV2
IFNG
GZMH
IFIT2
NSG1
PMEPA1
COPRS
MIR155HG
ETS2
RBPJ
LINC01943
AC068587.4
ITGA1
RYR2
SOCS2
CD24
CD79A
CD8B2
C9orf135
ADARB2
IGFBP2
SMIM24
SLC35F3
NDFIP2
LAYN
PAK1
CCDC141
TTN
KLRC2
CTSL
RGS16
ACP5
LGALS3
TNFSF13B
CCR4
FOXP3
CSGALNACT1
SUSD3
PKD2
STAT1
CTL-cc_C
CTL-cc_A
CTL-cc_B
Cluster0_iTreg_B
Cluster0_iTreg_C
Cluster0_iTreg_A
iTreg-cc_A
iTreg-cc_C
iTreg-cc_B
Input-T-Cells_B
Input-T-Cells_C
Input-T-Cells_A

***Supplemental Figure 2.*** *(**A**) Results of scVelo RNA velocity analysis. (Left) Calculated cellular latent time heatmap. Earliest latent time point appear in purple in naïve T-cell associated macroclusters. Latest latent time point appear in $CD103^+$ CD8iTreg and CTL macroclusters in yellow. (Right) RNA velocity stream visualization generated from scVelo. (**B**) Heatmap of top 25 DEGs identified for the four major cell types: $CD103^{hi}$ CD8-iTreg (orange) vs $CD103^{lo}$ CD8-iTreg (red) vs CTLs (blue) vs $CD8^+CD25^-$ input cells (green). (**C**) Total cytokine production (IL-2, TNFα, INFγ, IL-17α, IL-4, and IL-6) from CD8 CTLs (dark blue) vs CD8 iTreg (grey) after 4hr stimulation with PMA/Ionomycin, quantified by RayPlex® Human Inflammation Bead Array. **(D)** Relative expression and population frequency of $KIR^+$ CD8 Treg associated markers in CD8-iTreg compared to donor-matched, $CD8^+$ CTLs and $hCD4^+$ Treg (light blue).*

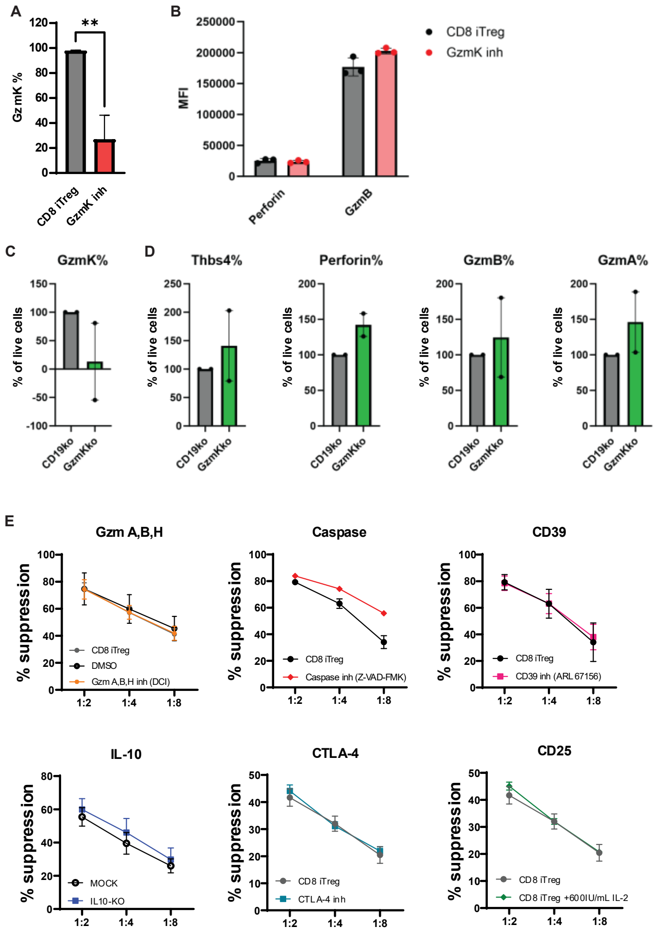


***Supplemental Figure 3. (A)*** *Frequency ±SD of GzmK$^{+}$ CD8-iTreg following treatment with 10µM of Bosutinib for 48hrs vs untreated control (n=3 biological replicates).* ***(B)*** *Expression of Perforin and Granzyme B in CD8-iTreg after 48hr treatment with 10uM Bosutinib (n=3 biological replicates). Frequency of* ***(C)*** *GzmK, and* ***(D)*** *Thbs4, Perforin, GzmB and GzmA in CD8-iTreg electroporated with GzmK CRISPR/Cas9 RNPs vs irreverent CD19-ko RNPs (n=2 biological replicates).* ***(E)*** *In vitro suppression of activated PBMC proliferation at different effector-to-*

**Commented [BRB1]:** check to see if supplemental figures are shown in their proper sequence as related to the figures so that Fig 1 has supplemental Fig 1; not sure

*PBMC ratios (1:2, 1:4, 1:8) vs PBMC-only controls (0:1) by CD8-iTreg (grey) vs CD8-iTreg following inhibition or blockade of GzmA/B/H, Caspase, CD39, IL-10, CTLA-4, or CD25, respectively; shown as mean ±SD for each ratio (n=3 biological replicates).*

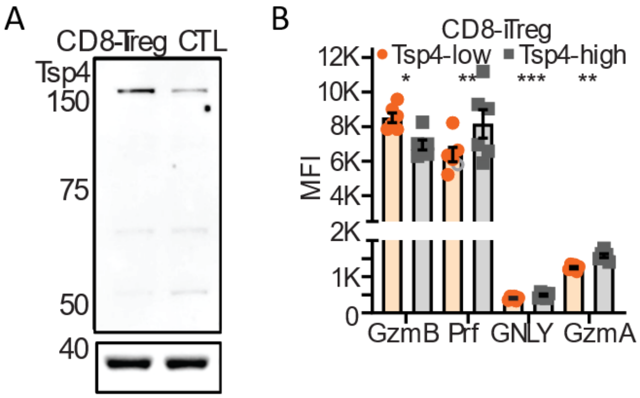


***Supplemental Figure 4. (A)*** *Western blot analysis of Thrombospondin-4 expression in CD8-iTreg and donor-matched CTLSMAPs, released on activating Quantifoil grids, coated with b-actin as loading control. (**B**) Flow cytometry analysis reporting mean fluorescent intensity (MFI)poly-l-lysine, anti-CD3ε (OKT3) and recombinant human ICAM-1-Fc/CD54; shown as mean ±SD (n=6 biological replicates). GzmB-mCherry-pHluorin mRNA-transfected CD8-iTreg were incubated on the activating Quantifoil grids for 90 min, followed by cell flushing of GzmB, GzmA, perforinthe cells out of the activating grids with ice cold PBS, and GNLY in plunge-freezing of the sample in liquid ethane.*

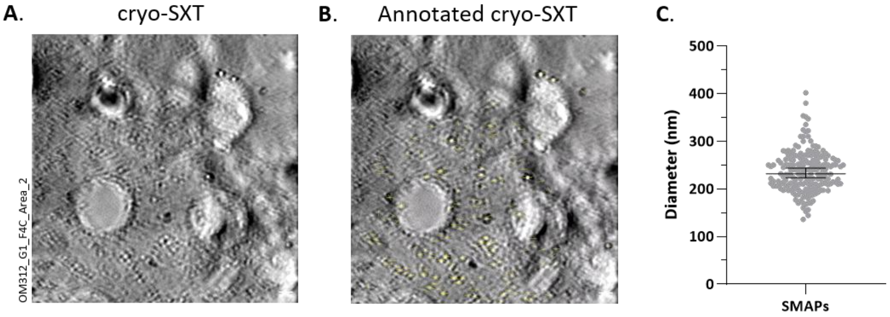


***Supplemental Figure 5.*** *(**A**) Manually reconstructed cryo-tomogram of CD8-iTreg SMAPs, released on activating Quantifoil grids, coated with poly-l-lysine, anti-CD3ε (OKT3) and recombinant human ICAM-1-Fc/CD54. GzmB-mCherry-SEpHluorin mRNA-transfected CD8-iTreg were incubated on the activating Quantifoil grids for 90 min, followed by cell flushing of the cells out of the activating grids with ice cold PBS, and plunge-freezing of the sample in liquid ethane. Imaging by correlative cryo-Structured Illumination Microscopy (cryo-SIM) with cryo-SXT followed under liquid nitrogen. Tomogram generation was performed in IMOD, with a manual fiducial model and Simultaneous Iterative Reconstruction Technique (SIRT) tomogram generation method with 12 iterations. (**B**) Annotations of the CD8-iTreg released SMAP diameters. (**C**) SMAPs released on activating Quantifoil grids, CD8-iTreg SMAP diameters by cryo-SXT.*

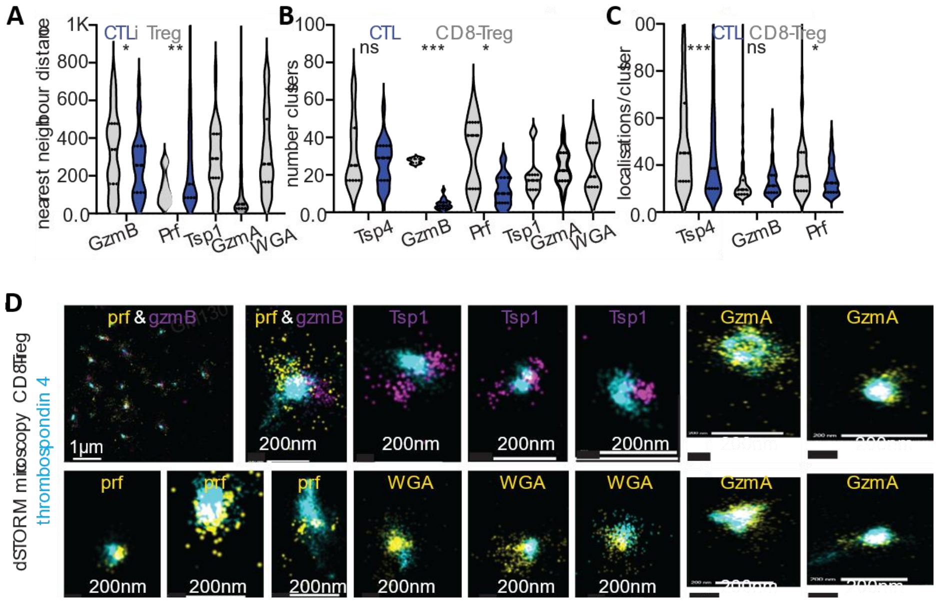


***Supplementary Figure 6. dSTORM analysis of SMAPs.*** *Two- or three-colour dSTORM datasets were analysed using the Oxford NanoImager (ONI) CODI platform. Localization, acquisition, and mapping files were processed with drift correction (DME algorithm; TetraSpeck bead reference). Frames were filtered to remove initial excitation peaks, retaining steady-state blinking events. Localizations were filtered by Gaussian sigma (~50–250 nm), photon counts (~300–500), and precision (≤20 nm). SMAP radius and localization number were quantified using clustering within regions of interest at the c-SMAC. Analyses included (**A**) nearest-neighbour distance, (**B**) cluster number (DBSCAN), (**C**) localizations per cluster, and (**D**) representative images with scale bars.*

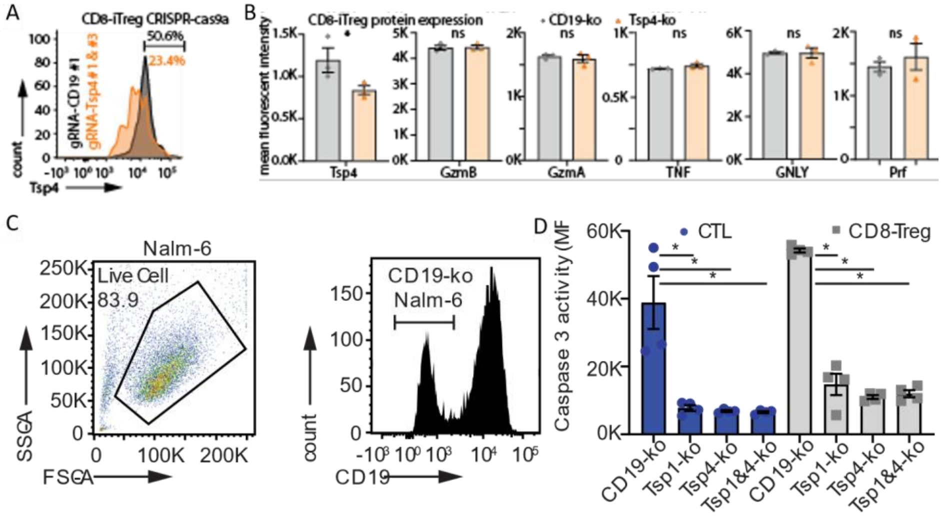


***Supplemental Figure 7.*** *Electroporation of CD8-iTreg with CRISPR/Cas9a Ribonucleoprotein complexes (RNPs) targeting (**A**) Thrombospondin-4 with 2 different gRNAs 24hrs apart, significantly reduces Tsp-4 expression, (**B**) without altering GzmB, A, Prf, TNF, and GNLY expression, compared to once targeted with control CD19 targeting RNP; shown as mean ±SD (n=2 biological replicates). (**C**) CD19 negative Nalm-6 were generated by hitting them once with CD19-RNP, followed by sorting of CD19 negative population. (**D**) Targeting Tsp-1, Tsp-4 or both, in $CAR19^+$CD8-iTreg significantly reduces their tumoricidal activity as determined by caspase 3 activity in $CD19^+$ Nalm6 4hrs post co-culture.*

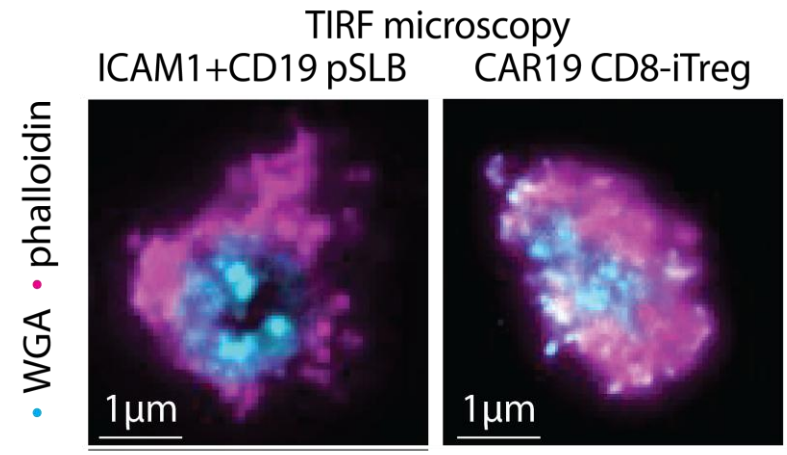


***Supplemental Figure 8.*** *CAR19⁺CD8-iTreg form immunological synapses on planar supported lipid bilayers with his-CD19 and ICAM-1-12his, as determined by actin rings visualized with fluorescent phalloidin and synapse-released glycoproteins by wheat-germ agglutinin.*

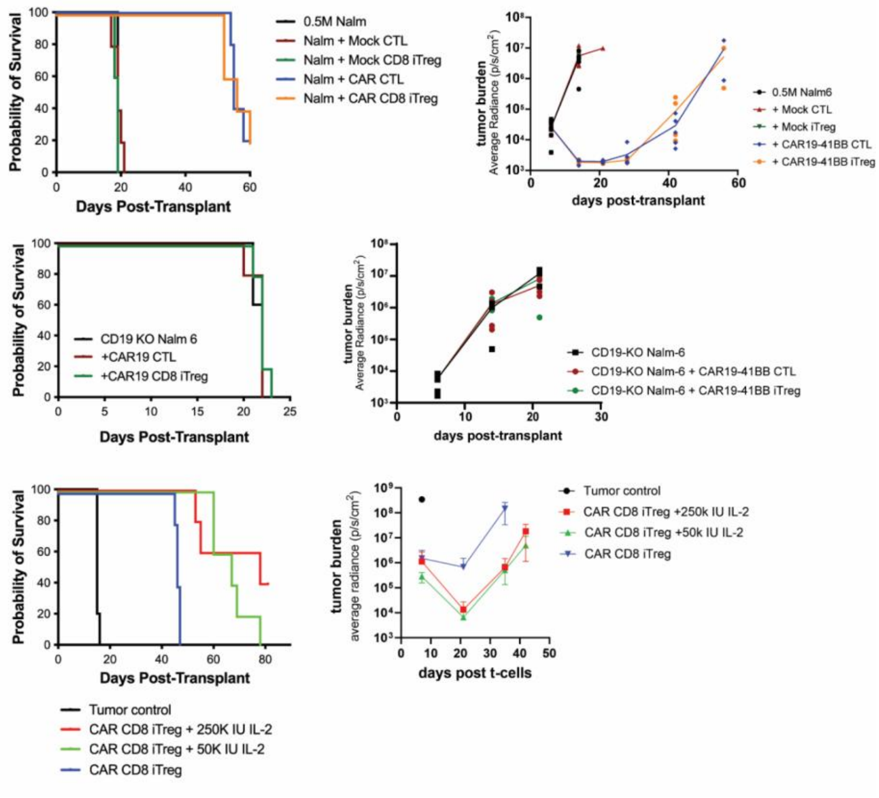


***Supplemental Figure 9.*** *Validation of Nalm-6 xenoGVL model* ***(A)*** *Nalm-6 survival and BLI for model infusing 5x10$^5$ Nalm-6 tumor D0, followed for CAR19$^+$ CD8-iTreg or CD8-CTLs (10$^7$/mouse) on D7 with supplemental IL-2 (250kIU/mouse; 3x/week for 21-days). (**B**) Nalm-6 survival and BLI for model infusing 1x10$^6$ CD19-KO Nalm-6 tumor targets, followed for CAR19$^+$ CD8-iTreg or CD8-CTLs (10$^7$/mouse) on D7 with supplemental IL-2 (250kIU/mouse; 3x/week for 21-days). (**C**) Nalm-6 survival and BLI for model infusing 1x10$^6$ Nalm-6 tumor targets on D0, followed by D7 infusion of CAR19$^+$ CD8-iTreg (10$^7$/mouse) with mice receiving 250k IU/mouse IL-2 vs 50k IU/mouse IL-2 vs vehicle control 3x/week for 21-days.*

**Human single cell RNA sequencing Sample Preparation and Analysis.** CD8$^+$CD25$^-$ T-cells were enriched from non-mobilized PB apheresis products and differentiated into either CD8+ CTLs or CD8-iTreg as described above. Triplicate biological samples were Hashtagged with TotalSeq-C0251 antibodies (BioLegend) before being pooled. Replicate pools were prepared from CD8$^+$CD25$^-$ T-cell inputs as well as from both CD8$^+$ CTL and CD8-iTreg outputs. Samples were prepared with MOJO dead-cell remover to improve viability before encapsulation with 10x Chromium. A 5' gene expression and CITEseq(*62*) library were prepared before sequencing on the Illumina NovaSeq X platform to a read depth of 2.5 billion reads. Sequencing quality control, alignment to the GRCh38-2024-A genome(*63*), UMI counting, and gene expression matrix generation were performed using CellRanger v9.0.1(*64*). CITE-seq Count(*65*) was used to process hashtag oligo library for demultiplexing. Samples were deconvoluted using a consensus majority-vote assessment via HTODemux, GMM-Demux(*66*), MULTI-seq(*67*), and BFF-Cluster(*68*) before data was loaded into Seurat v5(*69*) for quality control filtering, and data normalization. In Seurat dimensional reduction and universal manifold projection was performed and Louvain algorithm was used for clustering which revealed the $CD103^{+}$ CD8iTreg subset. In addition to Seurat biomarker discovery method cells were pseudobulked by cell class and biological replicate for differential expression analysis via DESeq2(*70*) and gene set enrichment analysis against the Gene Ontology (GO)(*71*), KEGG(*72*), MsigDB Hallmark(*73*), and MsigDB Immunological signatures databases using clusterProfiler(*74*). For RNA Velocity and trajectory inference Velocyto(*75*) was used to separate spliced from unspliced gene matrices and scVelo(*76*) was used to determine decay kinetics and splicing ratios. Finally, CellRank(*77*) and Palantir(*78*) were used to determine terminal populations and predict lineage specific pseudotime from developmental trajectories. Analysis code available upon request to the author.
